# Emerging transient dynamics at finite observation scales

**DOI:** 10.1101/2023.12.27.573307

**Authors:** Sergio Cobo-López, Lucas J. Carbajal, Matthew Witt, Forest L. Rohwer, Antoni Luque

**Affiliations:** Viral Information Institute, San Diego State University, San Diego, CA 92182, USA; Department of Biology, San Diego State University, San Diego, CA 92182, USA; Department d’Enginyeria Química, Universitat Rovira i Virgili, Tarragona, Catalonia, Spain; Department of Biology, University of Miami, Miami, FL 33146, USA; Department of Physics, San Diego State University, San Diego, CA 92182, USA; Holland Artificial Intelligence, San Diego, CA, USA; Department of Physics, University of Miami, Miami, FL 33146, USA; Department of Chemical, Environmental and Materials Engineering, University of Miami, Miami, FL 33146, USA

**Keywords:** Transient dynamics, dynamic regimes, dominant processes, observation timescale, Boolean dynamics, predator-prey systems, bacteriophages

## Abstract

Biogeochemical, ecological, and astronomical systems are transient. Yet, predicting dynamical shifts remains challenging because of inferential uncertainties in datasets and the limitations of asymptotic-dependent theories. To address this problem, we developed a theoretical framework that builds on the finite nature of observations. This framework assesses the relative importance of processes, defined as the mechanisms that contribute to the rate of change of the system’s dynamic variables; this allows prediction of switching thresholds, that is, the values of the dynamic variables at which a process becomes relevant or irrelevant, switching the dynamics into a new dynamic regime. The framework predicts that the number of possible observable dynamic regimes increases exponentially with the number of processes. Yet observers are expected to experience only a subset of those dynamic regimes, determined by which processes remain relevant throughout the observation. The framework was tested on a case study of a classic predator-prey system comprising four processes parameterized for bacteria (prey) and lytic bacteriophages (predator). The analysis recovered the sixteen dynamic regimes predicted by the framework, including two non-trivial quasi-equilibrium dynamics. The classical quasi-equilibrium solution, with all processes being relevant, was observed in only about one in ten simulated trajectories that covered the observational window. An adaptive Boolean model, using only relevant observable processes, validated the framework’s accuracy by recovering the full model’s dynamics within the observer’s tolerance. The observational framework introduced here provides a strategy for identifying the processes and conditions that lead to significant switches in dynamic regimes. The framework complements asymptotic approaches to transient dynamics by focusing on the observer’s specific, finite context rather than the system’s asymptotic states. It can also guide experimental design to measure a process’s impact in a dynamical system and suggests actions to suppress it.

**AUTHOR SUMMARY:** Identifying the causes behind dynamic transitions in ecological, climatic, and biological systems remains a major challenge. This study introduces a mathematical framework based on the premise that observing significant switches in a system’s dynamics depends on the observer’s finite empirical scales, including finite observation time and perceptual resolution. By assessing the relative importance of underlying processes (the mechanisms driving changes in dynamic variables), our framework predicts the threshold values at which the dynamics switch into new observable dynamic regimes. These regime switches differ from classical tipping points: they do not necessarily change the system’s attractors, but they change which processes influence the dynamics an observer can detect.

We demonstrate its rigor and value in a predator-prey system (bacteria as prey, lytic bacteriophages as predators), recovering 16 predicted dynamic regimes, including two non-trivial quasi-equilibria. An adaptive Boolean model using only relevant processes validated the framework’s accuracy by recovering the full model’s dynamics. This observer-centric approach offers a practical tool for identifying the processes that dominate the observed dynamics of complex systems. It also anticipates when new processes will become relevant and how to mitigate the switch. The case study focuses on microbial ecology, but the computational approach applies to any physical or biological system.

## INTRODUCTION

Natural systems are transient. The emergence of marine photosynthetic bacteria triggered a transition from low to high oxygen in the atmosphere 2.5 billion years ago [1,2]. The disruption of the thermohaline circulation in the Atlantic during the Younger Dryas period reversed the recovery from the last Glaciation period 12,000 years ago [3,4]. The increase in overfishing and eutrophication is favoring fleshy algae to outcompete calcifying corals and coralline algae, transforming the trophic structure in coral reefs towards a microbialized ecosystem [5–7]. These are just a few examples of the prevalence of transient dynamics in ecological, climate, and social systems [3,8–10]. However, predicting such major transitions and their consequences remains a pressing challenge.

A promising strategy for predicting transitions in natural systems is the use of early warning signals, based on statistical patterns in dynamic variables, as a proxy for an upcoming transition [11–14]. The majority of these methods are data-driven and use statistical and machine learning approaches to exploit or identify signals that can be informative, such as mean rates, conditional heteroskedasticity, or deep-learning-based patterns [11,15–19]. However, these early warning indicators strongly depend on the dataset used, leading to issues with reliability and uncertainty quantification [20]. Using physical-based frameworks grounded in the bifurcation-based theory of critical transitions offers a more generalizable approach as a guide for early warning signals [14,21]. This approach is based on the idea that transitions arise when slowly changing parameters cross bifurcations that alter the asymptotic attractors of the system, which need not be equilibrium states [9]. However, empirical tests have demonstrated that the assumptions underlying the application of critical transition theory do not always hold for natural systems [22,23].

A significant limitation of current theoretical frameworks that study transient dynamics is their reliance on concepts rooted in asymptotic dynamics and equilibrium states [3,9]. In this article, transient dynamics are defined as non-asymptotic dynamics, following Hastings et al. [9]. The influence of ideas and methods associated with asymptotic time comes from the fact that these are well-established and fruitful approaches in the mathematical theory of dynamical systems and their application in theoretical physics and mathematical biology [24,25]. Because asymptotic methods can obtain elegant theoretical results describing transitions when fast and slow processes co-occur, they provide a persuasive argument for their use in characterizing transient dynamics [25,26]. However, most studies that use mechanistic models to investigate the underlying processes causing transitions in dynamical systems rely on ad hoc analyses of the dynamic trajectories in the regions of interest [27–32]. Consequently, interpretations strongly depend on specific parameter values and are difficult to generalize, limiting the ability to predict when unexpected transitions occur. Sophisticated approaches that can extract transient dynamic regimes, like Lyapunov exponents, sloppy model theory, and low-dimensional manifolds, are not trivial to implement and do not necessarily lead to theoretical expressions revealing the underpinning causes of the transitions [33–38]. Thus, both data-driven and asymptotic-based mechanistic methods are currently limited in their ability to generalize insights from transitions in specific systems, posing a challenge for predicting the conditions that could drive future transitions in natural systems.

An underlying premise across current methodologies investigating transient dynamics is the assumption that the transient regimes perceived in a system, and the conditions under which they change, are intrinsic properties of the system that await being characterized or inferred [3,8–10]. What is often overlooked, however, is that it is the observer who ultimately distinguishes, based on measurements, perception, and prior knowledge, what is interpreted as a significant change in the dynamics of a system [39–41]. The temperature during a conference may be considered constant by the audience as long as it remains within a certain degree Celsius range, regardless of fluctuations below that threshold [42–44]. However, a one-degree change in laboratory conditions can result in a significant change in chemical reactions [45,46]. The same temperature oscillations may thus be discarded as noise in one context and carefully monitored and controlled in another. This perspective calls for methods that investigate transient dynamics over the scales of interest to an observer, that is, the finite observational time and resolution, rather than over asymptotic or supposedly intrinsic system properties. This article addresses this gap in transient-dynamics theory.

The framework introduced here builds on two empirical principles (Figure 1). First, any observer interacts with or measures a system for a finite amount of time. This defines the *observational time*, Δt_obs_. For example, if one collects data on bacterial growth in the laboratory over a 14-hour period, then Δt_obs_ =14 h. If one samples chemical factors over two weeks on an oceanographic cruise, then Δt_obs_ = 2 weeks or 336 h. If one aims to forecast the epidemiology of a virus outbreak over a year, then Δt_obs_ = 1 year or 8766 h. If a scientific community makes a multi-generational effort to monitor the dynamics of a lake over 200 years, then Δt_obs_ = 200 years or ∼1.75·10^6^ h. The observational time is thus the main characteristic time that an observer (an individual or a community) can, or is interested in, studying or predicting the dynamics of a system. Second, any observer is subject to an implicit finite (technological) perception that defines a meaningful change of a measurable or existing dynamic variable (such as temperature in the previous example) or an agent, like a virus population. In some cases, this idealized finite *observable change*, ΔA_obs_, may be limited by available instrumentation and experimental design, for example, the precision of a thermometer. It may also be associated with the known variance in the measurement due to environmental factors, for instance, not being able to reduce the resolution to less than 10% when comparing cell counts across samples. Only changes above this meaningful observable change, ΔA> ΔA_obs_, are considered perceptible or of interest in that observer’s context. A consequence of these two finite principles is that their ratio defines a reference rate of change or *relevant observable rate*, R_A, obs_ = ΔA_obs_ / Δt_obs_, of a variable that determines what is relevant in the context of the observer (Figure 1). Our hypothesis is that, during the system’s dynamics, only processes whose rates exceed the relevant observable rate at a given time can meaningfully contribute to the system’s trajectory from the observer’s viewpoint (the *observer’s reference frame;* Figure 1). We also propose that, when the rate of a process crosses the relevant observable rate, the transient dynamics switch to a new observable dynamic regime. The change in the process rate might reflect an internal change if it depends on a dynamic variable in the system, or external factors that may alter the parameter values that define the process.

**Figure 1.**
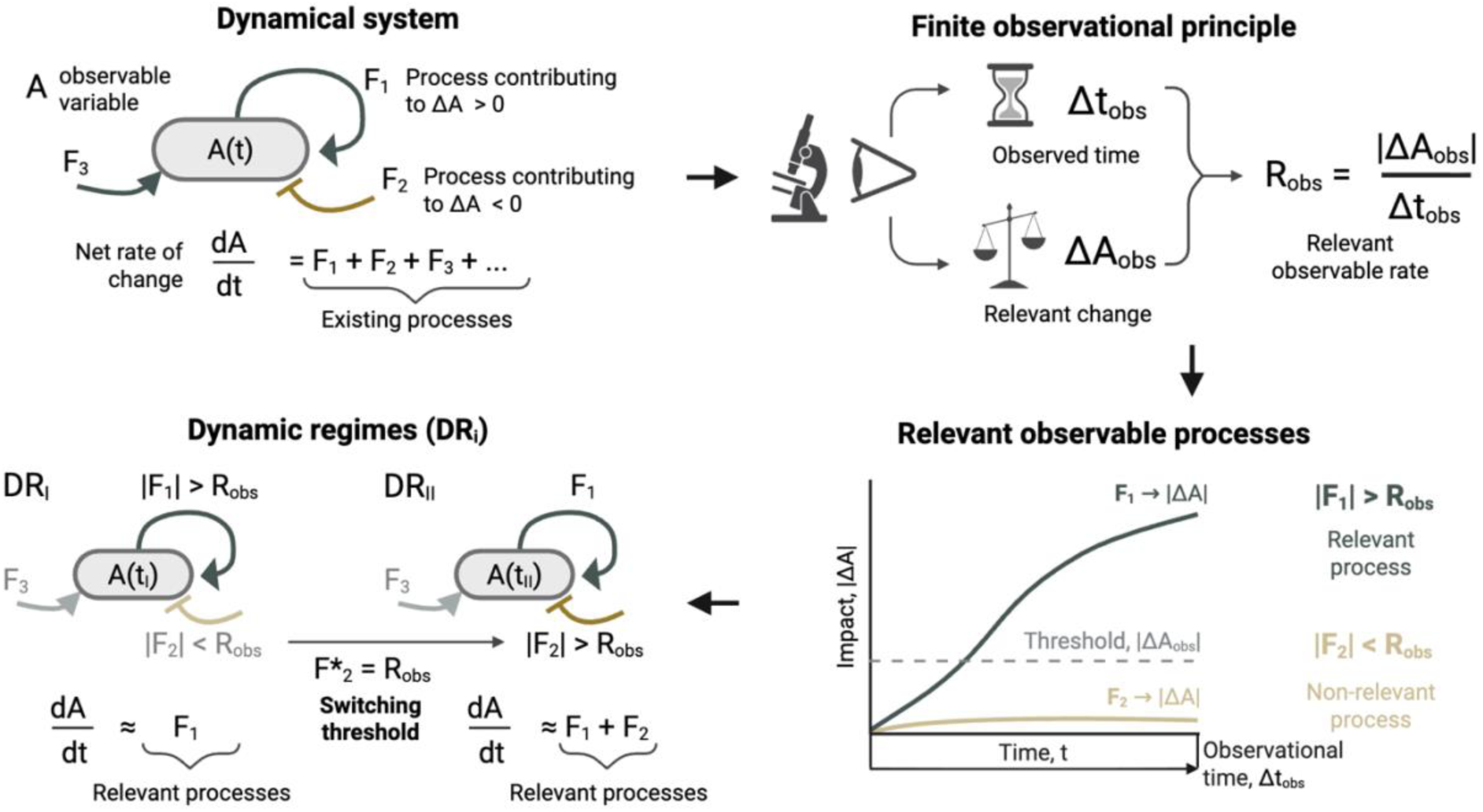
Dynamic regimes in the finite observational framework. The description below follows the same order as the arrows connecting the panels. (Top left) Diagram of a dynamical system illustrating the impact of different processes, F_i_, on the observable dynamic agent or variable, A(t). A few processes contributing to a positive change, ΔA>0 (dark blue arrow), or negative change, ΔA<0 (gold flat-head arrow), are illustrated. The equation at the bottom displays the contribution of the processes to the net rate of change, dA/dt. (Top right) Diagram illustrating the two finite observational principles: the finite observed time, Δt*_obs_*, and the relevant perceived change, ΔA*_obs_*. Their ratio leads to the relevant observable rate *R_obs_*. (Bottom right) Graph illustrating the individual contribution of two processes to the change of A over the observed time. The dashed horizontal grey line indicates the threshold associated with the relevant perceived change ΔA*_obs_*. Processes above this threshold are displayed as relevant (opaque) and below as non-relevant (semi-transparent). Their relationship with the relevant observable rate R^obs^ is displayed. (Bottom left) Diagram displaying a regime switch between dynamic regimes DR_I_ and DR_II_, each characterized by different relevant processes. The switching condition responsible for the regime switch is displayed as F*_i_. Figure created in BioRender as open access: https://BioRender.com/5g3o22z.

Here, this general framework was applied to a classic predator-prey system as a proof of concept. The predicted dynamic regimes were examined in the context of bacteria (prey) and bacteriophages (predators). The theoretical framework predicted transient regimes that had been previously overlooked, and the predictions were tested by measuring the error between the full dynamical system and the trajectory of the system including only the predicted observable processes. The addition of a new process, associated with the population’s carrying capacity, was used to test the framework’s ability to predict the conditions under which an additional process would be observable. The accuracy of these predictions was then tested across the empirical parameter space and five orders of magnitude of observer tolerance. The discussion section explains how insights from the finite observable framework guide diagnosis and prediction of transient dynamics.

## MATERIALS AND METHODS

### Identification of dynamic regimes and switching thresholds

The observational framework was defined based on the relevant observational time Δ*t_obs_* and relevant observational change Δ*A_i_*_,*obs*_ for each agent (or observable dynamic variable) *A_i_*. Their ratio defined the relevant observable rate 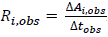. The dynamics of each agent *A_i_* was defined as the net rate of change, summing the processes *F_ij_* (Figure 1A). Normalizing the rates using the relevant observable rate yielded the weight of each process 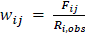. This weight is calculated at each time *t*, that is, *w* (*t*). A process was considered dynamically relevant if, at time *t*, its rate exceeded the observable rate, which, in terms of weights, led to the condition |*w_ij_*(*t*)| ≥ *w_t_*_ℎ_ = 1, with *w_t_*_ℎ_ being the relevance threshold. This condition is necessary, but not sufficient, to be significantly relevant because if a process’ weight above the threshold is short-lived and is not much larger than one, then the process’ contribution to the overall dynamics might not be that significant. Nonetheless, the condition serves as a practical, conservative operational strategy, allowing the observer to identify which processes may be relevant at time t, before further measurements, to support decision-making or action if needed or possible. It can also be applied to models that are not differentiable. A process, *F_ij_*, might depend on the system’s dynamic variables, that is, *F_ij_({A_k_})*, and so it can adjust its weight, *w_ij_({A_k’_})*. The discrepancy in the specific variables, indicated by the indices k and k’, is because the relevant observational rate, R_obs_, can be dynamic when, for example, relative observational changes are convenient, as shown later. In those cases, when dividing the process *F_ij_*by the observable rate *R_obs_* to obtain the weight *w_ij_*, some dependencies on dynamic variables can change. The values of the dynamic variables at which a weight reaches the relevance threshold define the switching threshold, *A_i,c_*. This captures the fact that changes in internal variables can drive dynamic switches even in the absence of external changes. A dynamic regime is defined as the region of state and parameter space in which a unique set of processes is relevant; a regime switch occurs when a trajectory crosses a switching threshold. The key steps in this process are illustrated in Figure 1.

### Analysis of dynamic regimes and switching thresholds in a Lotka-Volterra system for phage and bacteria

The dynamics of bacteria (prey), *A_1_* = *B*, and a lytic bacteriophage (predator), *A_2_* = *P*, were approximated using a standard Lotka-Volterra model [47–49]. The net rate of change of the bacterial population was defined by two processes: the bacteria’s growth rate, *F_11_ = F_g_*, and the phage infection or predation pressure, *F_12_ = F_p_* (see differential equations in Figure 2a). The net rate of change of the phage population was also defined by two processes: the phage production or burst when lysing an infected cell, *F_21_ = F_b_*, and the phage decay rate, *F_22_ = F_d_* (Figure 2a). The observational time was defined as *τ* = Δ*t_obs_*, while the relevant observational change of phage and bacteria was set, respectively, to Δ*B_obs_* = *α* · *B* and Δ*P_obs_* = *α* · *P*. Here *α* was the fraction of the population change considered to be relevant in the observation. This *α* can be interpreted as the observer’s relative tolerance. The rationale for choosing a dynamic observational change relative to the population at time t is that the dynamics in this system can be quite explosive, impacting the populations by orders of magnitude. Using a relative rather than an absolute change helps adjust the resolution dynamically to the current state. A fixed absolute threshold can be advantageous when dynamics vary less, or when measurement-error variance is absolute rather than relative, for example near a detection limit. The weights of each process and the switching thresholds (threshold concentrations) associated with regime switches were obtained by following the prescriptions described in the previous section and are presented in the Results. The dynamic regimes were defined by different combinations of relevant processes for the observer.

**Figure 2.**
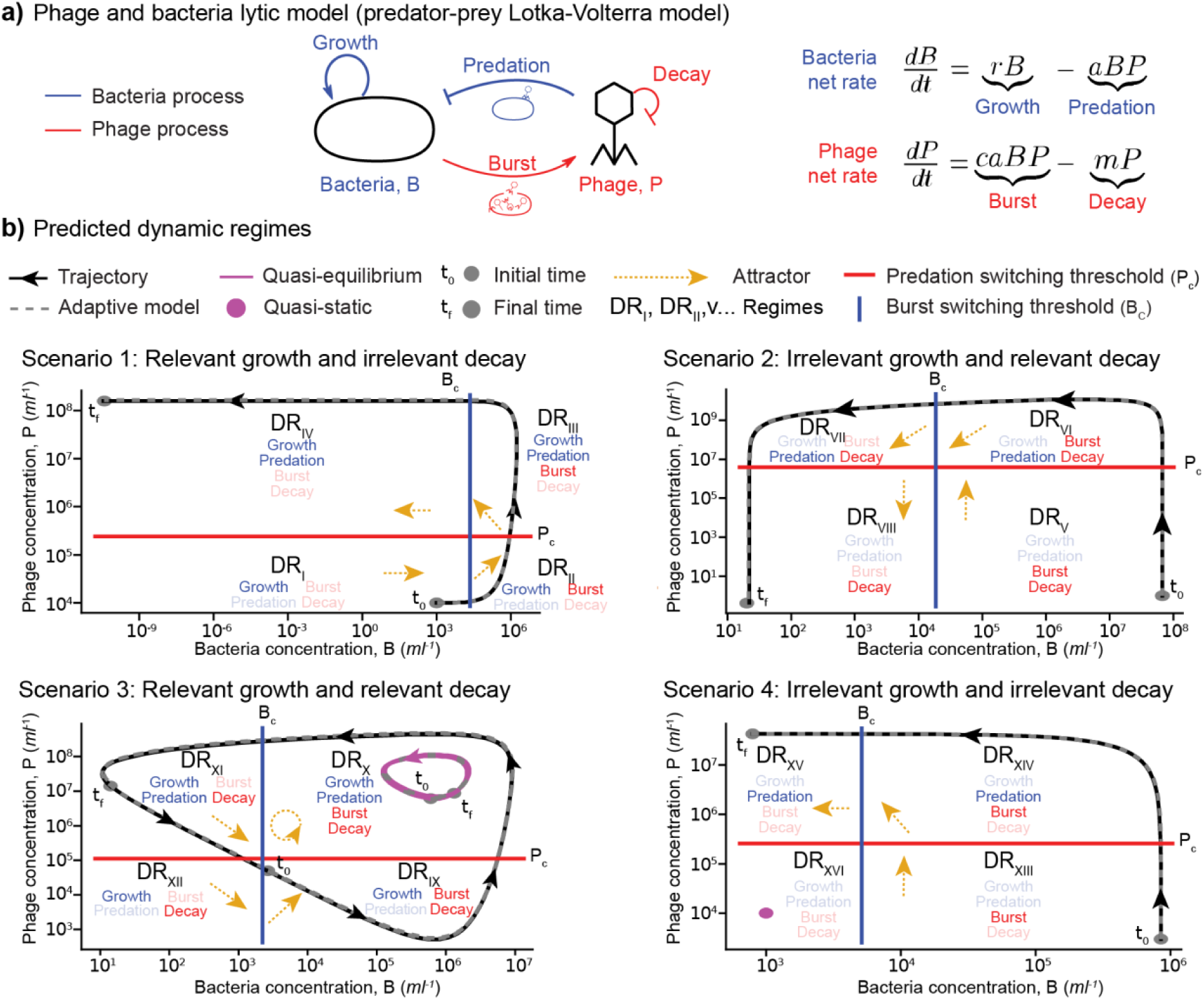
Dynamic regimes for a phage and bacteria lytic model. **a)** Diagram displaying the processes impacting the bacteria, B (blue), and phage, P (red), populations. The mathematical terms of each process are displayed in the net rate of change of each population, respectively, *dB/dt* and *dP/dt*. The parameters are *r* (growth rate constant), *a* (adsorption-infection rate constant), *c* (burst size), and *m* (decay rate constant). **b)** Dynamic regimes, DR_i_, are defined for four scenarios. These scenarios are predicted from an observational-principle-based transient-dynamics analysis and depend on whether bacterial growth or phage decay is relevant (or irrelevant). Each plot displays the phage concentration versus the bacteria concentration (black curve), with the black arrows indicating the time direction. The grey dots are the initial and end times. The magenta curves display quasi-equilibrium and quasi-static trajectories. The vertical (blue) and horizontal (red) lines represent the switching thresholds of bacteria (*B_c_*) and phage (*P_c_*). Each regime (quadrant) displays the labels of the relevant (opaque) and non-relevant (semi-transparent) processes. The yellow arrows indicate the local asymptotic attractors for each regime. The dashed grey curves display the approximate trajectories from the adaptive Boolean model, simulating only relevant processes. The parameters for the trajectories are listed in Table 1.

**Table 1.** Model parametrization. The columns display the values for the model parameters, bacteria and phage concentrations, and observational (obser.) time. The units are in parentheses. The rows distinguish the ranges used in the robustness analysis (empirical ranges for the parameters and observational time; sampled ranges for the initial concentrations) and the values used in each dynamic scenario. For each scenario, the concentrations of phage and bacteria correspond to the initial conditions used. (*) For the agents’ concentrations, the scenarios correspond to the initial values; scenarios with two lines of initial values correspond to the transient dynamics case (tr.) and the quasi-equilibrium case (q.-eq.). Observer finite scales: observational time, τ = Δt_obs_ (last column); finite resolutions, ΔB_obs_ = α B(t) and ΔP_obs_ = αP(t), where α = 0.1 (relative 10% tolerance) in all scenarios; observational reference rates, R_obs,B_ = ΔB_obs_/τ and R_obs,P_ = ΔP_obs_/τ. The Supplemental Material [50] (S2. Model Parametrization) includes how different references contributed to building these ranges [55–62].

|  | Growth<br>rate,<br>*r* (h<sup>−1</sup>) | Infection<br>rate,<br>*a* (ml·h<sup>-1</sup>) | Burst<br>size,<br>c (phages) | Decay<br>rate,<br>m (h<sup>-1</sup>) | \*Bacteria<br>concentration,<br>*B* (cells·ml<sup>-1</sup>) | \*Phage<br>concentration<br>*P* (phages·ml<sup>-1</sup>) | Obser.<br>time, *τ*<br>(h) |
|---|---|---|---|---|---|---|---|
| Ranges | 3·10<sup>−5</sup>–6 | 7·10<sup>-10</sup>–1·10<sup>-6</sup> | 2–5,000 | 10<sup>-3</sup>–0.8 | 1·10<sup>3</sup>–1·10<sup>12</sup> | 1·10<sup>3</sup>–1·10<sup>14</sup> | 0.8 – 16,800 |
| Scenario 1 | Based on laboratory values associated with fast growing bacteria (*E. coli*-like*)* and lytic phage (T4-like) |  |  |  |  |  |  |
|  | 0.9 | 3·10<sup>-8</sup> | 50 | 1.5·10<sup>-3</sup> | 10<sup>3</sup> | 10<sup>4</sup> | 14 |
| Scenario 2 | Based on environmental marine communities of slow growing bacteria and fast decaying phage (UV-exposed) |  |  |  |  |  |  |
|  | 3.2·10<sup>-5</sup> | 1·10<sup>-10</sup> | 225 | 1·10<sup>-1</sup> | 6.7·10<sup>7</sup> | 1·10<sup>0</sup> | 260 |
| Scenario 3 | Based on environmental communities (coastal-like) of relatively fast-growing bacteria and fast decaying phage |  |  |  |  |  |  |
|  | 1·10<sup>-1</sup> | 3·10<sup>-9</sup> | 50 | 1·10<sup>-1</sup> | 6·10<sup>5</sup> (q-eq)<br>2.7·10<sup>3</sup> (tr) | 6·10<sup>6</sup> (q-eq)<br>4.9·10<sup>4</sup> (tr) | 300 |
| Scenario 4 | Based on deep ocean-like communities of slow growing bacteria and slow decaying phage |  |  |  |  |  |  |
|  | 3.2·10<sup>-5</sup> | 3·10<sup>-8</sup> | 50 | 1.5·10<sup>-3</sup> | 1·10<sup>3</sup> (q-eq)<br>8.5·10<sup>5</sup> (tr) | 1·10<sup>4</sup> (q-eq)<br>3·10<sup>3</sup> (tr) | 13 |

#### Asymptotic analysis of the dynamic regimes

The theoretical attractors were derived as the asymptotic solution, that is, for infinite observational time, for each subset of differential equations within each dynamic regime. Each subset defines the reduced model of the regime. These regime attractors are not necessarily stable equilibria, and they describe the reduced model rather than the attractors of the full system, which do not change across regimes. If a fixed point existed (making the rates of the system equal to zero), then the linear stability of the fixed point was evaluated by the eigenvalues of the Jacobian, following standard practices [25]. The analysis is available in the Supplemental Materials [50] (S1. Asymptotic Analysis).

#### Model parametrization

The range of values for the agents, parameters, and typical observational times in the model was obtained from the literature for both laboratory experiments and environmental studies, as summarized in Table 1 and expanded in the Supplemental Material [50] (S2. Model Parametrization). In the numerical analysis, the observer’s tolerance (or relative resolution) was set to *α* = 0.1, that is, a 10% change in the population. This threshold was chosen because, in most cases, a 10% change provides a good resolution for measuring changes in environmental phage and bacterial populations, as well as in rapidly changing laboratory populations. Changing the tolerance, ⍺, makes an observer more or less sensitive to changes. The actual value of ⍺ depends on the context, prior knowledge, and technology available. In empirical studies, α can be estimated from replicate measurements, for example, as the coefficient of variation of replicate counts, and τ is the planned duration of the experiment, survey, or forecast. Because the switching thresholds and the minimum observation times scale linearly with α, the effect of a different resolution can be obtained analytically (see Results). The impact of the tolerance, α, was also tested numerically as described in the Robustness analysis section below.

#### Numerical simulations

Case studies illustrating all predicted dynamic regimes in the observational framework were explored following the theoretical analysis shared in the Results section. This resulted in four sets of parameter values (one for each scenario), which were selected based on a rationale that incorporated laboratory and environmental values (Table 1). For scenarios predicted to produce quasi-equilibrium solutions, two sets of initial conditions were used, capturing both quasi-equilibrium and transient trajectories. The system of differential equations was coded in Python and integrated with the *solve_ivp* function in the *SciPy* library [51]. This function relies on the LSODA solver from the FORTRAN library odepack [52], which automatically switches between non-stiff and stiff solvers depending on the behavior of the equations [53]. The simulations were performed with the developed transient-dynamics code (v0.2.3-alpha) [54], written in Python using NumPy, SciPy, pandas, and Matplotlib. The simulated dynamics, including populations, weights, and relevant processes, are provided in the Supplemental Material [50] (S3. Simulated Dynamics).

#### Error analysis of the predicted regimes

To test the accuracy of the observational framework, the full phage-bacteria model (Figure 2a) was compared with a dynamically adaptive Boolean model that integrated only the relevant processes in a regime [63,64]. The adaptive Boolean model was obtained by multiplying each process of the original model by a Heaviside step function *θ_i_*(*w_i_* − *w_t_*_ℎ_) = 1 for *w_i_* ≥ *w_t_*_ℎ_ and 0 otherwise, where *w_i_* was the weight of process *i* and *w_t_*_ℎ_ = 1 was the relevance threshold. This differential equation was integrated for the same initial conditions, parameters, and methods as the full model. The error for each population was obtained as the absolute difference between the value from the full and adaptive Boolean models at each time step. The relative error was defined as the absolute error divided by the value of the full model. The combined error was defined as the average of the relative errors of the populations at each time step. Summary statistics were obtained for each regime, scenario, and overall simulation dynamics.

### Analyzing the impact of adding a new process

The phage-bacteria model, defined by the Lotka-Volterra equations, was modified by incorporating a carrying capacity process as described in the Results section. This process accounted for the competition between bacteria for resources [24,47]. The weight of the carrying capacity process *w_K_*was obtained using the observational framework. Conditions associated with the quasi-equilibrium solution of the standard model were used to assess the impact of the process on the system (Table 1). The observational framework was used to predict the minimum observational time needed to observe the effect of the carrying capacity process (*w_K_ ≥ w_th_=*1). The full and adaptive Boolean models were simulated and compared to assess the errors of the observational framework.

### Robustness analysis

The full and Boolean phage–bacteria models, with and without carrying capacity, were integrated with *solve_ivp* (LSODA) on 10,000 evenly spaced time points over the observational time, τ. Model parameters (r, a, c, m) and τ were sampled within the ranges in Table 1, initial concentrations within 10^3^–10^12^ bacteria ml^−1^ (B_0_) and 10^3^–10^14^ phages ml^−1^ (P_0_), and the carrying capacity within K = 10^4^–10^13^ cells ml^−1^ with B_0_ ≤ K, using log-uniform Latin hypercube sampling (scipy.stats.qmc; seed 2026). Each sample was simulated with the same parameters across tolerances α = 10^−5^, 10^−4^, …, 1. A simulation was stopped when either population, in either model, fell below 10^−3^ ml^−1^ or exceeded the physical limits of 10^15^ bacteria ml^−1^ or 10^18^ phages ml^−1^ (the volumes of the smallest bacterium, about 10^−3^ µm^3^, and phage, about 10^−6^ µm^3^, packed into 1 ml); both models were then compared up to the earliest stop time. The sample size was doubled (2,000, 4,000, …) until every tolerance, α, had at least 1,000 simulations covering the full observational window (16,000 samples without and 8,000 with carrying capacity); 1,000 of these were selected at random for each α; stopped simulations with at least two compared time points were analyzed separately. The share of the sampled space with all processes relevant was calculated as the number of samples that covered the full window with all processes relevant throughout, divided by the total number of samples. The accuracy of the Boolean model was measured as the mean combined relative error, the time average of ½(|B_full_ − B_Boolean_|/B_full_ + |P_full_ − P_Boolean_|/P_full_) × 100. For each simulation, the fraction of time spent with 0 to N relevant processes (N = 4 without and 5 with carrying capacity) and whether all processes were relevant throughout was recorded. Extreme outliers were identified with Tukey’s far-out test on log_10_ error (values above Q3 + 3 × IQR) and excluded from the summary statistics, and 95% confidence intervals for proportions were calculated using the Wilson score interval. Scripts, tables, and figures for the robustness analysis are in results/alpha-robustness-v2 of the code archive [54].

## RESULTS

### General predictions derived from the finite observational reference framework

#### The identified dynamic regimes depend on the observation scales

The first relevant result of the observational framework approach outlined in the methods is the condition for a dynamic process to be considered relevant for a finite observer:

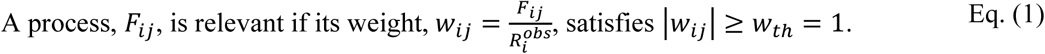

Here, *w_t_*_ℎ_ was the relevance threshold. The switching thresholds, that is, the values of the variables and parameters leading to 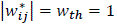 determine when a process switches from relevant to irrelevant or vice versa (Figure 1). Two observers might be subject to different observational rates, 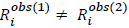 either through the observational time Δ*t^obs^* or through the relevant observational change 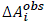.

In general, these observers will perceive different weights for the same process, 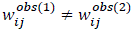, and, thus, different switching thresholds and dynamic regimes. Importantly, an asymptotic observational time, Δ*t^obs^* → ∞, or infinitely precise observational change, Δ*A^obs^* → 0, yields infinite weights *w_ij_* → ∞, in which case any process that is not strictly null is always relevant, no switching thresholds exist, and the classical asymptotic description is recovered. Therefore, the switching thresholds and dynamic regimes emerge from the interplay between the system and the observer’s finite observational reference framework. In contrast, the attractors and bifurcations of the full system, including classical tipping points, remain intrinsic properties of the system and do not depend on the observer.

### The dynamic regimes increase exponentially with the number of processes

Relevant processes driving the dynamics can push variables across switching thresholds, making previously irrelevant processes relevant, or vice versa. (Figure 1). Each process in a dynamical system can be either relevant or irrelevant, so the number of dynamical regimes D that an observer can measure is predicted to grow exponentially with the number of processes M in the system:

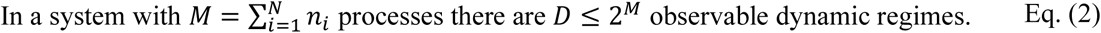

Systems with more processes, thus, can lead to exponentially more complex observable dynamics. Suppose a system remains in the same dynamic regime during the observational time. In that case, the attractor of the reduced model of that regime will remain constant during the observation, and the observer will perceive the dynamics as more predictable or *quasi-steady*. Suppose the dynamics stay in the same regime, and the variables oscillate around or converge towards a constant value. In that case, the observer may interpret the system as being in a *quasi-equilibrium state*. In the limit case, where the value of dynamic variables remains within the relevant observable change, the observer may perceive the system as *quasi-static*.

### The case study of a classic predator-prey system in the context of phage and bacteria confirmed the general predictions

#### Sixteen dynamic regimes were observed, distributed in four observational scenarios

The normalization of the phage and bacteria (Lotka-Volterra) model (Figure 2a) using the observational framework yielded the equation rates in terms of the weights, *w*, for the four processes (*M* = 4) in the system, that is, bacterial growth (*w_g_*) and phage predation (*w_p_*) for the bacterial rate of change, and the phage production or burst (*w_b_*) and phage decay (*w_d_*) for the phage rate of change:

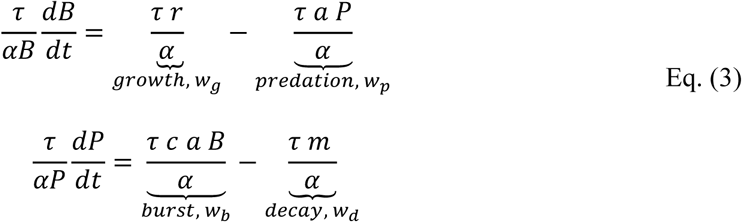

Here, *r* was the bacterial intrinsic growth rate constant, *a* the adsorption-infection rate constant, *c* the phage burst size, and *m* the phage decay rate constant. The parameter *α* was the relative observational resolution or tolerance used in the relevant observational changes Δ*B^obs^* = *αB* and Δ*P^obs^* = *αP*. Thus, the observational relevant rates, R^obs^, are dynamic and proportional to B(t) and P(t), respectively. Since the growth rate and decay rate processes are proportional to the bacteria and phage, respectively, *F_g_* ∝ B(t) and *F_d_* ∝ P(t), rescaling the dynamical system using the dynamic relevant observable rates led to weights for the growth rate, *w_g_*, and phage decay, *w_d_*, that were independent of the state variables, B and P, Eq. (3). These processes were predicted to be observable (*w ≥ w_th_*=1) for observational times *τ* ≥ *τ_g_* = *α*/*r* and *τ* ≥ *τ_d_* = *α*/*m*, respectively. Thus, once a tolerance, α, is fixed, the observational times depend only on the intrinsic growth rate, r, and intrinsic decay rate, m, respectively. These parameters can vary over time depending on external factors, like temperature [55,57]. However, for the sake of simplicity but without losing generality, these parameters were kept constant in the current analysis. The case for dynamic parameters (non-autonomous dynamical system) is explored in the Discussion section. The weights for the predation, *w_p_*, and phage burst (or production), *w_b_*, depended dynamically on the phage population, *P*, and the bacterial population, *B*, respectively, Eq. (3). Thus, the relevance threshold, *w_t_*_ℎ_ = 1, predicted the switching thresholds (threshold concentrations) *P_c_* = *α*/(*τ a*) and *B_c_* = *α*/(*τ c a*) for the predation and phage burst to become relevant processes, respectively. Thus, depending on the dynamic trajectories of the bacteria and phage populations, the system is expected to switch dynamic regimes when they cross these threshold concentrations. These thresholds depend on the adsorption-infection rate, *a*, and the burst size times the adsorption-infection rate, *c·a*. These properties depend on factors that can change over time, like the viscoelasticity and affinity of the fluid (e.g., mucus) with the agents [65,66] or the nutrients impacting the host metabolism [47,57]. However, as mentioned above, these parameters were kept constant, and the non-autonomous case is addressed in the Discussion section.

The predator-prey system is constituted by four processes, M=4 (Figure 2). Thus, the observer’s reference framework predicts up to D ≤ 2^M^ = 16 measurable dynamic regimes, Eq. (2). To observe all regimes, it is necessary that all possible combinations of relevant/irrelevant process states are accessible. In this case, since the weights for the growth, *w_g_*, and decay, *w_d_*, were constant in the reference observational framework used, it was possible to explore the dynamic regimes in four observable scenarios (Figure 2b). Scenario 1 is defined by relevant growth and irrelevant decay, Scenario 2 by irrelevant growth and relevant decay, Scenario 3 by relevant growth and decay, and Scenario 4 by irrelevant growth and decay. The theoretical asymptotic analysis of each regime (dynamic subsystem with a unique set of relevant processes) identified the attractors of the reduced model of each regime for each observable scenario, as indicated in Tables S1 (Scenario 1), S2 (Scenario 2), S3 (Scenario 3), and S4 (Scenario 4) in the Supplemental Material [50] (S1. Asymptotic Analysis). The attractor analysis for the different scenarios predicted quasi-equilibrium when either all processes remained relevant (magenta trajectory in Scenario 3) or all processes remained irrelevant (magenta trajectory in Scenario 4).

The numerical exploration of the scenarios illustrated in Figure 2b was associated with different qualitative biological situations as described in Table 1. The weight analysis and error of the observational framework for each scenario are explored below and displayed in Figures 3 (Scenario 1), S1 (Scenario 2), 4 (Scenario 3), and S2 (Scenario 4). One important result in this case is that it was possible to generate dynamic trajectories that recover the four different dynamic regimes in each observable scenario. That is, the sixteen dynamic regimes could be measured. This exhaustive exploration may not be possible or recommended for systems with a larger number of processes, given that the number of possible regimes is predicted to increase exponentially, Eq. (2). Nonetheless, the specific scenarios described below illustrate how, for specific finite observational scales, the framework can characterize the processes that are relevant and what conditions will lead to transition switches associated with a distinct measurable dynamic regime.

**Figure 3.**
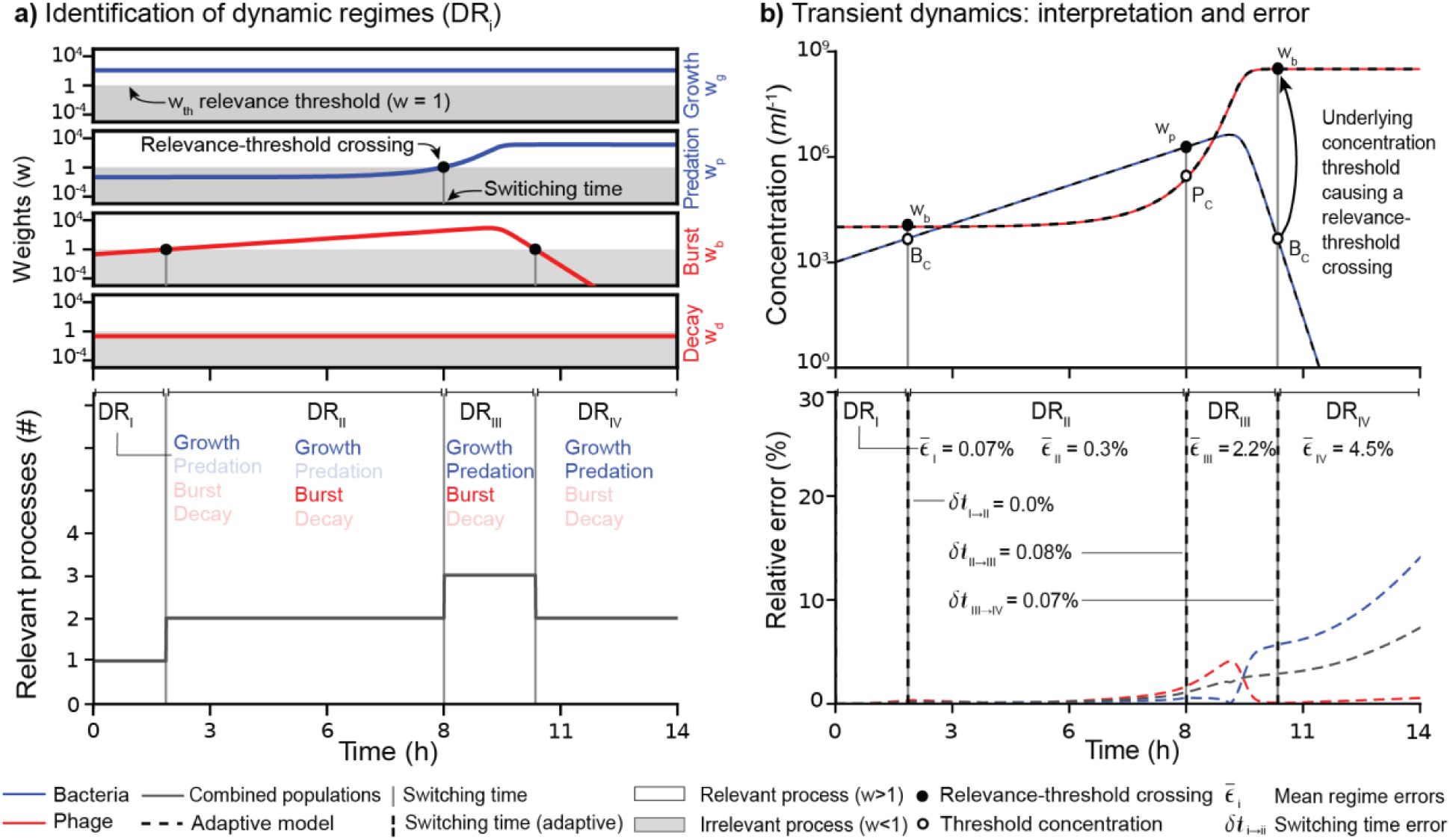
Transient dynamics analysis using the finite observational framework. Dynamic regimes and transient dynamics for the case of relevant growth (wg > 1) and irrelevant decay (wd < 1), corresponding to Scenario 1 in Figure 2. The model parameterization is given in Table 1. **a)** The dimensionless weights for each process (top panel) and the number of relevant processes (bottom panel) are plotted as a function of time. Processes associated with bacteria and phage are shown in blue and red, respectively. Black circles indicate crossings of the relevance threshold, and vertical grey lines indicate the switching times, separating regimes, DR_i_. **b)** The top panel shows the dynamics of bacteria (blue) and phage (red) as well as the Boolean model (dashed black lines), while the bottom panel shows the relative error of the adaptive Boolean model. The mean relative errors for each regime, 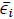, and the error in the switching times, *δt_i→ii_*, are displayed. Concentrations are shown on a logarithmic scale; populations decline toward zero and do not become negative.

#### Processes predicted to be relevant accurately reproduce transient dynamics

The first biological scenario, related to Figure 2b, Scenario 1 (relevant growth and irrelevant decay), was investigated using parameters and observational times consistent with laboratory experiments for fast-growing bacteria, such as *E. coli*, and lytic phages, like T4 phage (Table 1). The observational time used was *τ* = 14 h, which relates to the usual duration of a lab experiment. The resolution for the relevant observational change was 10% of the population (*α* = 0.1), given that the populations can change several orders of magnitude during the observational period. The weight analysis identified four dynamic regimes based on the combination of relevant processes (Figure 3a): bacterial growth (DR_I_), bacterial growth and phage burst (DR_II_), bacterial growth, phage burst, and phage predation (DR_III_), and bacterial growth and phage predation (DR_IV_). The fixed parameters in this scenario analytically predicted a bacterial switching threshold of *B_c_* = 4.8·10^3^ cells/ml, regulating the phage production (or burst) process, and a phage switching threshold *P_c_* = 2.4·10^5^ phages/ml, regulating the predation process. These regimes and their associated switching thresholds distilled the dynamics of the full bacteria and phage model (Figure 3b top). The bacterial population initially increased exponentially, driven by bacterial growth, while the phage population remained relatively constant. The crossing of the bacterial switching threshold (*B > B_c_*) at 1.73 h made the phage burst relevant. The crossing of the phage switching threshold (*P > P_c_*) at 8.40 h rendered phage predation relevant, which rapidly outweighed the bacterial growth (*w_p_ >> w_g_* = 126). This yielded a rapid decline of bacteria, which crossed back its switching threshold (*B < B_c_*) at 10.59 h, making the burst process irrelevant and stalling phage production, which remained relatively constant because the phage decay process was irrelevant throughout the dynamics. The dynamics of the adaptive Boolean model, which integrated only relevant processes (*w ≥ w_th_*), were visually indistinguishable from those of the full model (Figure 3b top). The error analysis indicated that the accuracy varied across the regimes, yielding an average relative error (combined populations) of 1.58% (Figure 3b bottom), with the bacterial population’s most significant error in the fourth regime (mean error = 8.76%) and the phage in the third regime (mean error = 2.09%) (see S4. Relative Error). The main source of discrepancy between the full model and the Boolean model comes from the fact that they do not necessarily reach the switching thresholds at the same time because of the processes that are not considered relevant in the Boolean dynamics. However, since in each switch the populations can be in an upward or downward trajectory, the error does not necessarily accumulate; it instead fluctuates across regimes. It is important to note that the Boolean model was run independently of the full model; there was no information transfer between them.

It is worth noting two important observations that may apply to the other scenarios and systems. First, for a set of initial conditions, the combination of the attractors of each regime (yellow arrows in Figure 2b) and the switching concentrations (vertical and horizontal lines in Figure 2b) makes it possible to predict the qualitative trajectory of the system without the need for numerical simulations. For example, in observational scenario 1 (relevant growth and irrelevant decay) just covered (Figure 2b and Figure 3), at the initial time, *t_0_*, one starts in dynamic regime I, DR_I_. The attractor points towards an increase in the bacterial population, which eventually crosses the bacterial switching concentration, *B_c_*, and the burst becomes relevant. That moves the system to dynamic regime II, DR_II_, where the attractor predicts “infinite” growth of both phage and bacteria. Thus, the phage population eventually crosses the phage switching concentration, *P_c_*, and predation becomes relevant, moving the system to dynamic regime III, DR_III_. Since the weight for the growth is constant and the weight of predation increases with the phage concentration, this leads inevitably to an attractor that pulls the system to further increase the phage population while decimating the bacteria population. This eventually leads the bacterial population to cross the switching threshold, rendering the burst process irrelevant and moving the system to dynamic regime IV, DR_IV_. In this regime, the attractor maintains the phage population constant because no process is relevant affecting this population. Because this locks the phage concentration above the switching threshold, predation keeps driving down the bacterial population. For a finite population, this will inevitably lead to extinction, unless a mutant is resistant to phage infection, which is often what occurs experimentally [28], but goes beyond the scope of this study. It is possible to carry out a similar graphical qualitative analysis for all cases in this study (Figure 2). A key point is that one does not need to investigate all possible dynamic regimes in advance. Instead, in a given dynamic regime, one needs to ask in what direction the local attractor is pulling the variables, check what processes (relevant or irrelevant) depend on those variables, and determine if any switching thresholds could be crossed. That provides the recipe to be prepared for the next dynamic regime and opens a window of time for the observer to act and influence the system. For example, in the scenario described above, a key switch that eventually leads to bacterial extinction occurs when the phage population exceeds the concentration threshold, *P* > *P_c_*, making predation relevant in the dynamic regime DR_III_. Since this concentration threshold, *P_c_* = *α*/(*τ a*), is inversely proportional to the adsorption-infection parameter, *a*, any cofactor (natural or engineered) that interferes with phage adsorption, for example, a protein that binds the host-receptor-binding domain on the phage, would decrease *a* and increase *P_c_*, preventing predation from becoming relevant. This, in turn, will increase the bacterial threshold concentration, *B_c_* = *α*/(*τ c a*), delaying the switch to make the burst process relevant between the dynamic regimes DR_I_ and DR_II_. If one aims to grow the bacteria and reach a relatively constant culture where the phage does not significantly impact the population during the observational time, and assuming the media has a carrying capacity *K*, then if B_c_ > K, the bacteria will never reach the concentration necessary for the burst process to produce phage significantly during the observational time. This would require a modified adsorption-infection constant satisfying *a*′ < *α*/(*τ c K*). Using the values for Scenario 1 and aiming for a population of K ≈ 10^8^ cells/ml, then a’ < 1.43·10^−12^ ml/h, which is four orders of magnitude smaller than the original adsorption-infection constant, *a*, in this scenario (Table 1). To prevent the phage burst from being relevant at longer observational times, the adsorption-infection constant would need to be reduced accordingly. This, thus, provides the specific actions and targets necessary for the observer to act on the system to alter the dynamics.

A second important observation involves the decoupling of causality and the relevance of the processes on the dynamics. In the first dynamic regime, DR_I_, only bacterial growth is relevant (Figure 3). When the bacterial population reaches the switch concentration, *B_c_*, the burst process becomes relevant (DR_II_). The burst is a consequence of the phage infecting and lysing (killing) bacteria, yet this predation process is not relevant in the observer reference framework. Thus, the observer can measure an increase in the phage population but cannot infer that this increase results from bacteria dying in the process. After the phage population reaches the switch concentration, *P_c_*, the predation effect becomes apparent. This apparent violation of the mechanism’s causality is because each infection releases c=50 infective phages (Table 1), amplifying the burst signal relative to predation. This highlights the need to investigate the process weights in each reference framework, as they determine whether the effect might be measurable before the associated cause.

Returning to the analysis of the other dynamic regimes, the opposite transient dynamic scenario, that is, one with irrelevant bacterial growth and relevant phage decay, as shown in Figure 2b (Scenario 2), utilized parameters related to marine systems, featuring slow-growing bacteria and fast phage decay occurring at the sea surface due to sunlight effects (see Table 1). The observational time was set to *τ* = 260 h (about 11 days), and the tolerance or resolution in the observational change was again 10% (*α* = 0.1). The parameters used yielded the switching thresholds *B_c_* = 1.7·10^4^ cells/ml and *P_c_* = 3.8·10^6^ phages/ml for bacteria and phage, respectively. The weight and trajectory dynamic analysis captured four expected regimes (Figure S1): DR_V_ (burst and decay), DR_VI_ (predation, burst, and decay), DR_VII_ (predation and decay), and DR_VIII_ (decay). Initially, phage burst production exceeded decay, driving rapid phage growth and the subsequent decline of the bacterial population. After the bacterial population dropped below its switching threshold, phage burst became irrelevant, resulting in exponential phage decay. Note that this trajectory can be qualitatively sketched graphically following the attractors in each regime (Figure 2b). The adaptive Boolean model yielded a mean (combined) error of 0.31% with maximum errors of 0.76% for bacteria and 0.31% for phage (see S4. Relative Error).

### Two observable quasi-equilibrium regimes in the Lotka-Volterra dynamics

The quasi-equilibrium dynamic predicted when all processes were relevant (Table S3) was recovered in Scenario 3 (Figure 2b and Figure 4). The parameters for Scenario 3 were similar to those for the marine populations in Scenario 2, but assumed a faster-growing bacterial population, which is possible in enriched environments such as inhabited coastal ecosystems (Table 1). The observational time was *τ* = 300 h (nearly two weeks), and the tolerance to change was again 10% (*α* = 0.1). The parameters used yielded the switching thresholds *B_c_* = 2.2·10^3^ cells/ml and *P_c_* = 1.1·10^5^ phages/ml for bacteria and phage, respectively. The initial conditions were above the switching thresholds for both bacteria and phage, and the constant weights, growth (*w_g_*) and decay (*w_d_*) (see Eq. 3), were of the same order, 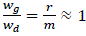, due to similar values of the rate constant (*r*) and decay rate constant (*m*) (Table 1, Scenario 3). These two conditions yielded oscillations around the classical theoretical equilibrium of the Lotka-Volterra system for bacteria and phage, respectively, 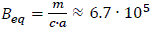 cells/ml and 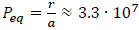 phages/ml, with all processes remaining relevant throughout the dynamics (Figure 4a), resulting in no errors in the adaptive Boolean model (see S4. Relative Error).

**Figure 4.**
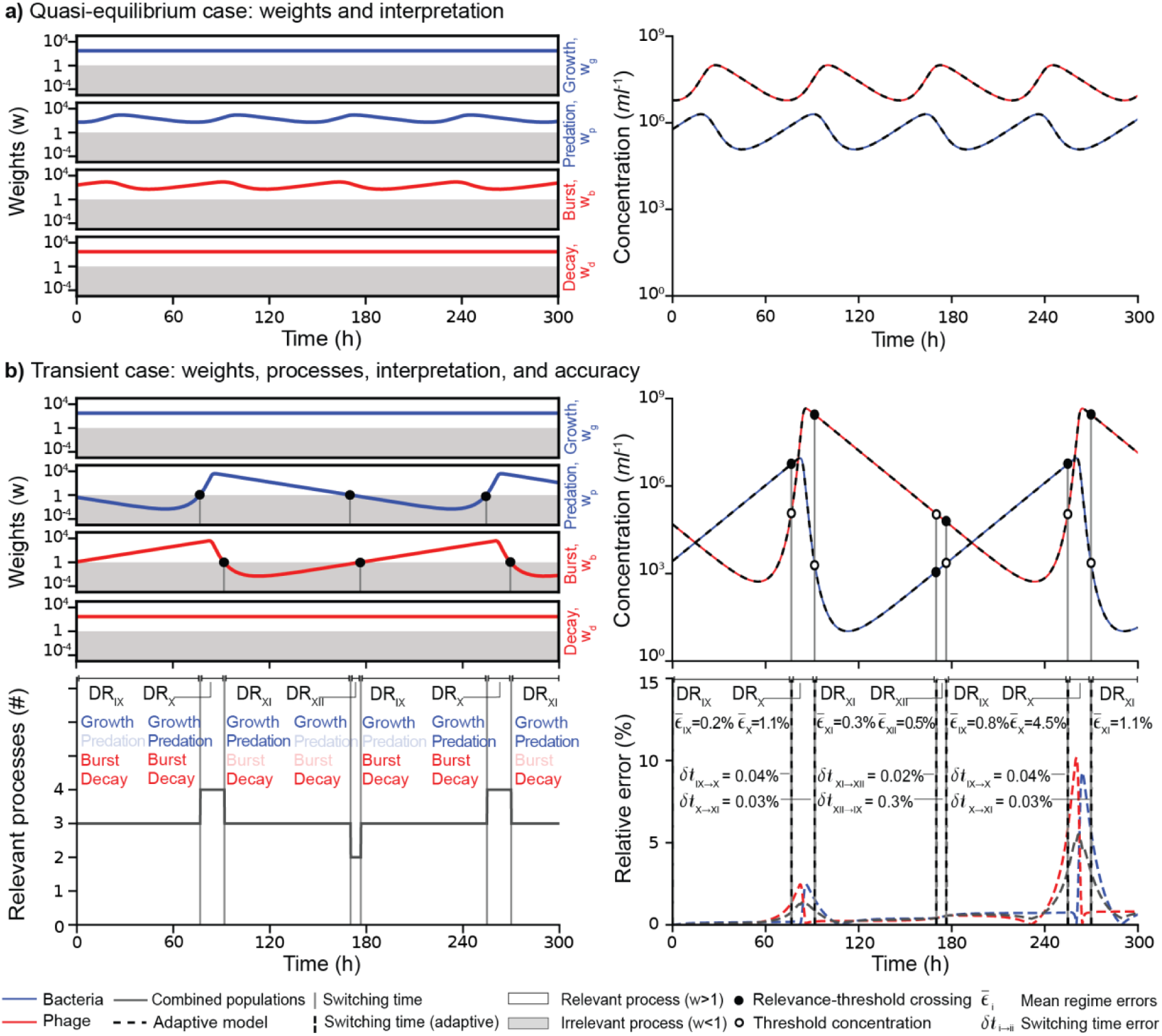
Scenario displaying quasi-equilibrium. Dynamics in the scenario with relevant growth (*w_g_>1*) and relevant decay (*w_d_>1*), corresponding to Scenario 3 in Figure 2. Model parameters are given in Table 1. The color coding and symbols are analogous to Figure 3. **a)** The left panel shows the weights for phage (red) and bacterial (blue) processes over time. The right panel shows the dynamics of phage and bacterial concentrations. The dashed lines represent the adaptive Boolean model. Because all processes remain relevant, its predictions coincide with those of the full model, and no error plot is included. **b)** The top two plots are analogous to those in **a),** but with initial conditions that lead to transient dynamics. The bottom plots include the relevant processes over time (left) and the relative error of the adaptive Boolean model (right). The solid circles indicate crossings of the relevance threshold, and the vertical lines indicate switching times. The mean relative errors for each regime, 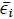, and the error in the switching times, *δt_i→ii_*, are displayed. The dynamic regimes, DR_i_, are displayed in both bottom panels following the same notations as in Figure 2.

Initial conditions where at least one of the populations was below the switching threshold were predicted to make one of the variable processes (predation or burst) initially irrelevant. The conditions used in this case rendered predation initially irrelevant and fueled transient dynamics that alternated between four regimes (Figure 4b, left): DR_IX_ (growth, burst, and decay), DR_X_ (all relevant), DR_XI_ (growth, predation, and decay), and DR_XII_ (growth and decay). The phage and bacterial populations oscillated abruptly, crossing their switching thresholds repeatedly (Figure 4b, right), which was associated with the broader cycle in the phase diagram (Figure 2b, Scenario 3). The adaptive Boolean model, which integrated only relevant processes, accurately reproduced the dynamics of the full model (Figure 4b, right). The mean combined error was 0.74%, and the errors in the bacterial and phage populations increased locally during the short regimes with abrupt population changes (Figure 4b, right-bottom).

The second quasi-equilibrium situation was predicted when all processes were irrelevant, resulting in all populations remaining quasi-static (Table S4). This prediction was captured in the case with irrelevant growth and decay, corresponding to Scenario 4 (Figure 2b, magenta trajectory). The parameters used resembled the environmental values of Scenario 2, using slow phage decay instead, a plausible situation in the deep ocean (Table 1). The observational time was shortened to be similar to laboratory conditions *τ* = 13 h, as in Scenario 1. The tolerance to change was again 10% (*α* = 0.1). The switching thresholds were *B_c_* = 5.1·10^3^ cells/ml and *P_c_* = 2.6·10^5^ phages/ml for phage. Initial conditions below the switching thresholds (*B_0_* = 1·10^3^ cells/ml < *B_c_* and *P_0_* = 1·10^4^ phages/ml < *P_c_*) rendered all processes irrelevant, yielding relatively constant concentrations of phage and bacteria for the observer. The adaptive Boolean model yielded a combined average error of 0.09% (see S4. Relative Error). Using initial conditions above the bacterial switching threshold (*B_0_* = 8.5·10^5^ cells/ml > *B_c_*) made the burst process relevant (Figure S2a). This increased the phage concentration and eventually decimated the bacterial population below the switching threshold (Figure S2b). The adaptive Boolean model yielded an error of 2.06% with respect to the full model (Figures S2b and S4. Relative Error).

#### The observational framework accurately predicts when new processes become relevant

In general, when modeling the interaction of agents in dynamical systems, it is difficult to judge what details of the system might be important to capture its dynamics in a given context. To illustrate how the observational reference framework can help in deciding what the potential relevance of additional processes that may be considered of interest is, a carrying capacity term was added to illustrate the competition of bacteria for resources, − 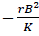, where *K* was the carrying capacity (Figure 5a). This term introduced an asymptotically globally stable equilibrium 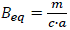 *and* 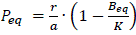. However, the observational framework predicts that the effect of this process would be only observable if its weight 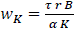 is relevant (*w_K_* ≥ *w_th_* = 1). Using the conditions for quasi-equilibrium from Scenario 3 and setting up the carrying capacity to 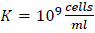 (a reasonable upper value in coastal conditions) yielded a weight below the threshold *w_K_* = 0.18 < *w_t_*_ℎ_ = 1 for the carrying capacity process at the initial conditions (and at most 0.6 along the oscillations). Therefore, the process remained irrelevant throughout the dynamics (Figure 5b). The system, thus, displayed the same periodic quasi-equilibrium oscillations already observed in the model without carrying capacity (compare Figures 4a and 5b). The adaptive Boolean model produced a dynamic that was visually indistinguishable from the full model and yielded a mean combined error of 1.0% (Figure 5b and S4. Relative Error).

**Figure 5.**
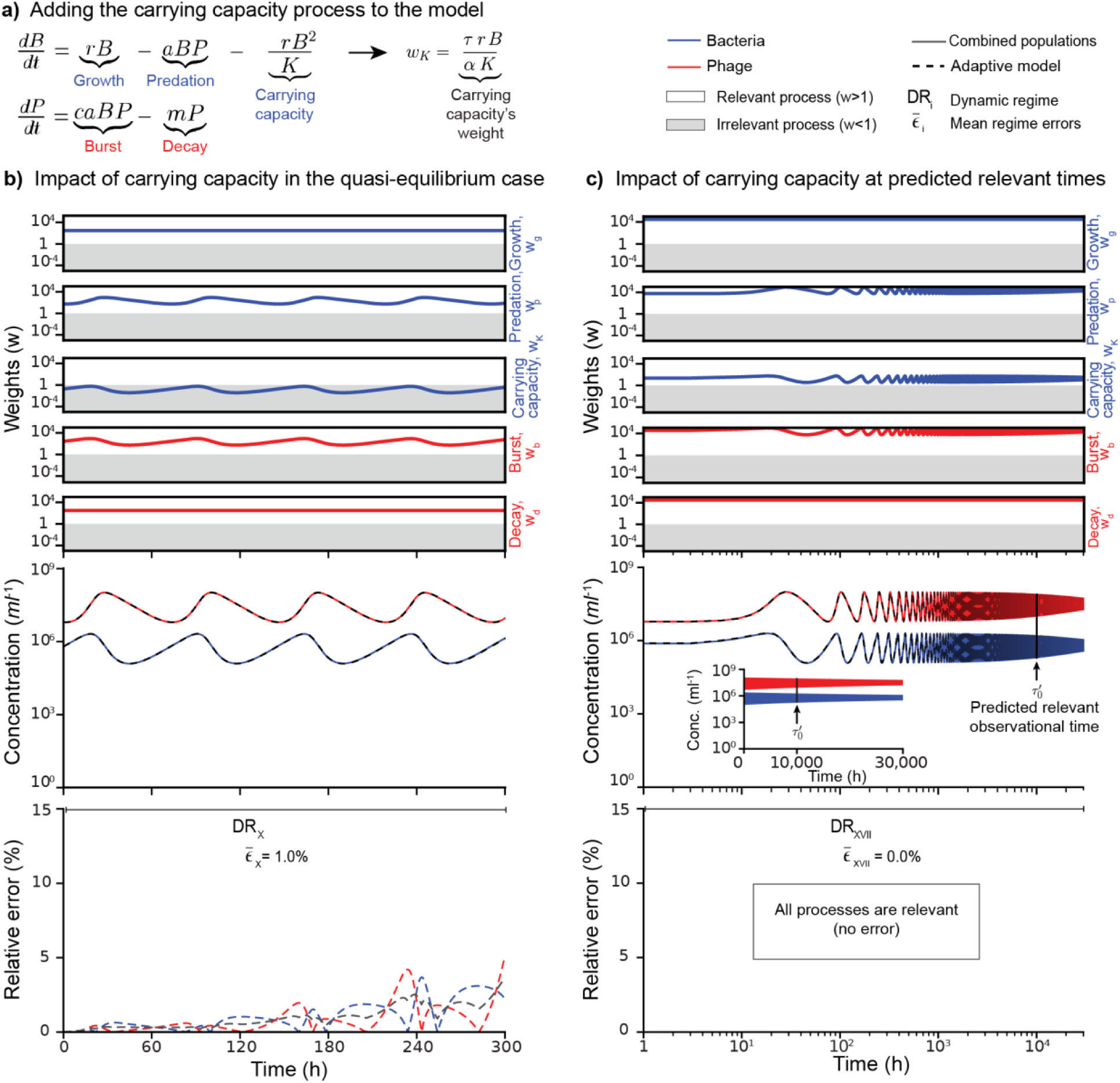
Impact of adding a new process to the dynamics: the carrying capacity. The original model was modified to include the carrying capacity process, which limits bacterial growth. The model parametrization is shared in Table 1. **a)** Phage and bacteria model including the carrying capacity. The weight of the carrying capacity process is displayed as *w_K_*. The color coding and symbols in this figure are analogous to Figures 2 to 4; the subset used is displayed to the right. **b)** Transient dynamic analysis for the same quasi-equilibrium case (relevant growth and decay, Scenario 3) displayed in Figure 4a. The plot for the weights includes the carrying capacity process. The population dynamics of phage and bacteria (red and blue) and the adaptive Boolean model’s (dashed lines) error are plotted. The regime and mean error are displayed. **c)** Transient dynamics for the same case as in **b)** but for a larger observational time. The time is displayed on a logarithmic scale. The inset plot displays the time on a linear scale. The vertical arrows indicate the predicted minimum observational time, τ_0_’, to observe the impact of the carrying capacity. DR_XVII_ denotes the regime in which all five processes, including the carrying capacity, are relevant.

One classical interpretation of the irrelevance of the carrying capacity term introduced above is that bacterial population concentration remains far below carrying capacity throughout the dynamics (B << K), so the carrying capacity term is not of leading order and removing it from the dynamics might just have a minor effect. However, the observer reference framework provides an additional strategy for determining how to observe and measure the impact of bacterial competition through carrying capacity. Assuming the system’s properties (parameters) cannot be changed, the most accessible experimental design strategy is to modify the observational time. The observational framework predicted that the carrying capacity process would become relevant for observational times *τ*′ ≥ *τ*_0_′ = *αK*/(*rB*_min_) ≈ 8.3·10^3^ h, that is, about one year, where *B*_min_ ≈ 1.2·10^5^ cells/ml was the minimum bacterial concentration along the oscillations. Setting an observational time larger than this switching time, *τ*′ = 30,000 h > *τ*_0_′, made the carrying-capacity process relevant throughout the dynamics (*w*_K_ ≥ 3.6; Figure 5c). The predicted global asymptotic value had not been reached yet, but the variations (or oscillations) of the phage and bacteria populations diminished over time, which is the measurable impact of the carrying capacity. Another insight is that the variability of the phage and bacteria populations can be perceived differently by the observer depending on the sampling time. If the observer samples at regular intervals throughout the 30,000 hours of the observational time, this linear sampling will capture a dispersion of phage and bacteria populations that decrease over time (inset in Figure 5c – middle). If the sampling strategy is stratified, combining more frequent sampling at first and more sparse sampling as time progresses, then it would be possible to perceive the oscillations of the populations and how they changed over time (Figure 5c – middle). In any case, for this larger observational time, all processes remained relevant, so the adaptive Boolean model yielded no errors (Figure 5c bottom).

#### The adaptive Boolean error remains within the observer’s tolerance, and only a small fraction of conditions led to all processes being relevant

To test the reliability of the observer’s reference framework under different conditions, the parameter space, that is, the model parameters, initial concentrations, and observational times, was sampled using Latin hypercube sampling, and the adaptive Boolean model was compared with the full model (Figure 6). For a 10% observer tolerance (α = 0.1), the mean combined relative error across 1,000 simulations covering the observational time window remained within the observer’s tolerance in 99.7% of cases, and the errors in the three exceptions were close to the tolerance (Figure 6a). The error distribution was bimodal because a subset of simulations, 9.5% (95 out of 1,000), had all processes relevant throughout the entire window and were errorless. The median error of the remaining 90.5% of simulations was 0.79%, and the 90th percentile was 2.3%, well within the observer’s tolerance (Figure 6a). The simulated conditions produced a wide range of process combinations across the time window, with two or three processes being relevant most often (Figure 6b). The two cases associated with quasi-equilibrium, that is, all processes relevant or irrelevant throughout the time window, represented 9.5% (95 out of 1,000) and 4.6% (46 out of 1,000), respectively (Figure 6b). The exploration of the Boolean approach of relevant processes across observers’ tolerances spanning five orders of magnitude (α = 10^−5^ to 1) showed that the error with respect to the full model increases with the observer’s tolerance α, and the error in most simulations remained well within this tolerance (Figure 6c). As tolerance α decreased, the observer (the Boolean model) became more sensitive, and the fraction of times all processes were relevant increased, leading to a significant drop in the approach’s median error (Figure 6c). The trend was similar for simulations that had to be stopped because populations crossed unphysical values (Figure S3), which represented the majority of simulated cases (∼89.5% across tolerances). The same robustness analysis was performed for the model including the carrying capacity process and yielded similar findings (Figures S4 and S5). The only major difference was the presence of extreme outliers for α = 0.1 (1 out of 1,000) and α = 1 (4 out of 1,000) among the simulations covering the full observational time window (Figure S4c). For the main case of interest in this study, α = 0.1 (10% tolerance), the extreme outlier corresponded to long oscillatory trajectories in which the Boolean model accumulated a persistent delay with respect to the full model, yet the Boolean trajectories reproduced the period and amplitude of the oscillations of the full model (Figure S6).

**Figure 6.**
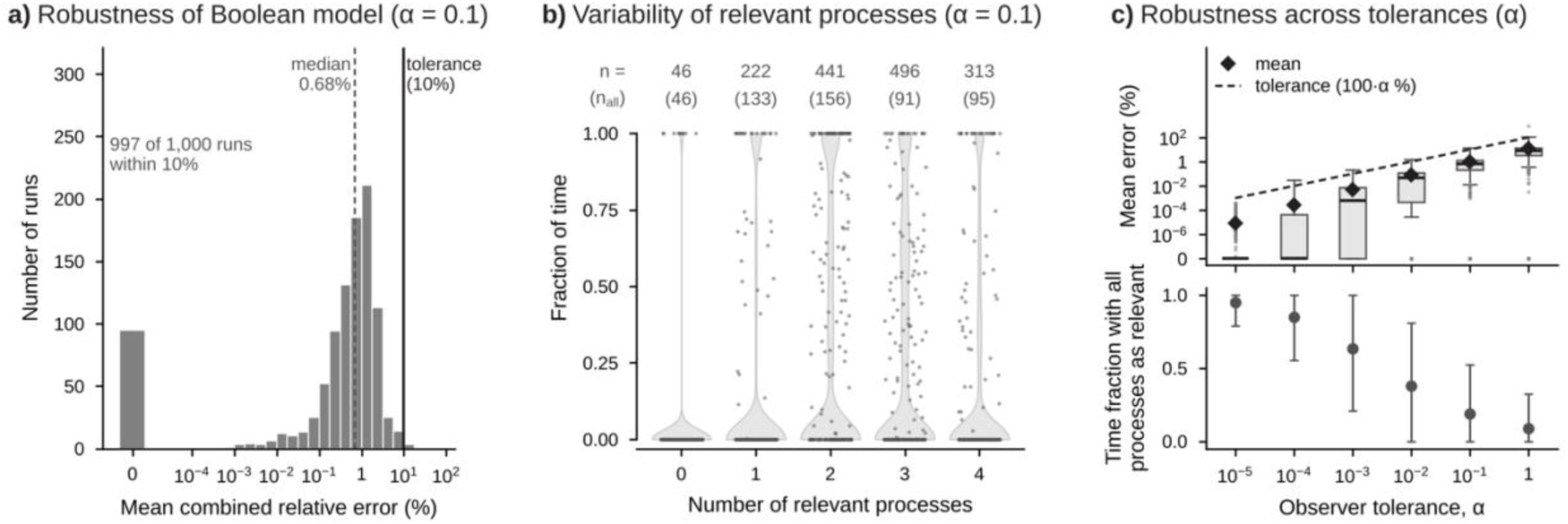
Robustness analysis of the adaptive Boolean model. Latin hypercube sampling (log-uniform) of the parameters, initial concentrations, and observational time of the phage–bacteria (Lotka–Volterra) model; only simulations that remained within the physical bounds for the full observational window are shown (n = 1,000 for α = 0.1). **a)** Histogram of the mean combined relative error (%) between the adaptive Boolean and full models at α = 0.1; the bar labeled 0 counts runs with zero error (all processes relevant throughout), the dashed line marks the median, and the solid black line marks the observer tolerance (10%, corresponding to α = 0.1). **b)** Fraction of the observational time that each run spent with 0 to 4 relevant processes (weight w ≥ 1) at α = 0.1; violins show the distribution across runs, gray points are a random sample of 200 runs, the numbers on top (n) are the runs that spent any time with that number of relevant processes, and the numbers in parentheses (n_all_) are the runs that spent all of the observational time with that number. **c)** Top, box plots of the mean combined relative error for α = 10^−5^ to 1 (box, interquartile range, IQR; line, median; whiskers, 1.5 × IQR; points, values beyond the whiskers), with the mean (◆) and the observer tolerance, 100·α % (dashed line); zero errors are drawn at the bottom of the axis, labeled 0. Bottom, mean ± standard deviation of the fraction of observational time with all 4 processes relevant. Extreme outliers (Tukey’s far-out test, above Q3 + 3 × IQR of log_10_ error) are excluded from the statistics and, when present, shown in ochre.

## DISCUSSION

The application of the finite observational framework introduced here illustrates how a system with a limited number of underlying processes, such as the standard predator-prey Lotka-Volterra system, can display a wide range of dynamical regimes for a real observer (Figure 2). The sixteen dynamic regimes characterized in this system, two of which can sustain quasi-equilibrium trajectories, contrast with the classic asymptotic analysis of the Lotka-Volterra model, which expects a periodic solution and a trivial (null concentrations) equilibrium [24,47]. A real observer, instead, would perceive the classic periodic solution only when all four processes (growth, predation, burst, and decay) are relevant throughout the observational time (Figure 2 and Figure 4). This makes the conditions for quasi-equilibrium a far more stringent set of conditions than in the classic analysis. The concentrations of both agents must remain above their switching thresholds, so the amplitudes of the oscillations do not cross any of them (Figure 4). The exploration of the parameter space for an observer with a 10% tolerance (α = 0.1) indicates that the classic quasi-equilibrium scenario, with all processes relevant throughout the dynamics, occurred in only 9.5% of the simulated trajectories that covered the observational window, and in about 1% of the sampled parameter space (Figure 6). When adding the carrying capacity, the case with all processes active reduced even further to 1.3% (Figure S4).

The finite observable framework also predicts scenarios that are not captured in the classic asymptotic analysis, such as the existence of a quasi-static solution when the processes are not relevant (Figure 2) or when a potential additional process, like the carrying capacity, remains irrelevant (Figure 5). Most of these transient regimes, predicted and measured within the finite observable framework, would not have been accessible using published methods for transient dynamics. Inference methods that rely on data would not be able to predict the drastic effects of a new process becoming relevant with a trajectory where its weights until that moment would be negligible within the observer’s tolerance [11,20]. The transient dynamics methods that rely on asymptotic assumptions, such as those based on bifurcation-induced critical transitions, are not designed to recover the dynamic regime switches described here. These switches depend on the impact of the state variables on making processes relevant or irrelevant. Instead, those transitions arise from changes in the attractors of the system, that is, changes in its parameters [3,9,14]. Although this study fixes the parameters in each scenario, the approach can also be applied when parameters change over time. In that case, the switching thresholds *B_c_* and *P_c_* in the example covered here would change over time as a function of the implicit time dependence of the parameters. Nonetheless, the framework specifies that classical asymptotic behavior occurs only if all processes are relevant. Since the switching thresholds change with the parameters, their changes would align with the changes in stable states when the parameters cross a bifurcation. The weights used in the framework relate to classical tools in applied mathematics. A process whose weight is well below the relevance threshold behaves as a regular perturbation that can be neglected at leading order, and it becomes relevant when the dynamics push its weight above the threshold. This is analogous to the change of dominant balance across a boundary layer [26]. The weights are non-dimensionalized mappings of each process, but the reference scales are set by the observer’s finite observation time and resolution rather than by the system’s intrinsic properties. The fact that resolution can be dynamic when relative resolutions are preferred, as shown in the examples above, is an important distinction from classical non-dimensionalization approaches.

The robustness analysis also shows that the adaptive Boolean model error scales with the observer’s tolerance (Figure 6c). A heuristic argument explains this behavior. By definition, an irrelevant process changes the corresponding population by less than the observable change, αB or αP, over the observational time, so neglecting it introduces a relative error of order α or smaller within each regime. The error can accumulate, however, when the trajectory crosses many switching thresholds, as in long oscillatory dynamics, where small delays in the switching times compound from cycle to cycle (Figure S6).

On a separate note, the sloppy model approach could identify that some of the processes are irrelevant, but its underlying dependency on asymptotic manifolds would overlook the fact that some processes can transition from irrelevant to relevant and vice versa [33]. The research presented here demonstrates instead that the local attractors associated with relevant processes at a given time drive the dynamics until a switching threshold is crossed, altering the set of relevant processes (Figure 2 and Table S1). This also supports the drastic effect that local events can have globally on a system [67,68].

The transient regimes are identifiable from the relevant processes, as demonstrated by the fact that the adaptive Boolean model, which simulates only relevant processes and runs independently of the full model, accurately reproduced the dynamics (Figures 2 and 6, and S3-S5). These regimes provide a guide for distilling specific systems, as seen in the case study on phage and bacteria investigated here. If the life traits (model parameters) of a bacterial host and phage pair, along with their initial concentrations, do not fall within the stringent ranges for quasi-equilibrium, transient dynamics are expected. Empirical parameter ranges suggest that the lytic lifestyle often leads to transient phage-host pair dynamics that deviate substantially from classic quasi-equilibrium (Figures 2 and 6). We hypothesize that community effects, such as the evolution of virus-host (predator-prey) networks [69–76], holobiont networks [77,78], and alternative lifestyles, like lysogeny, where the virus integrates into the host [57,79–82], contribute to the diverse range of stable concentrations of viruses and bacteria observed in microbial communities across ecosystems [56,62,83,84]; testing this hypothesis will require applying the framework to multispecies systems. The identification of relevant processes depending on the parameter ranges can help simplify theoretically driven approaches to phage therapy [28,32,85,86].

A key advantage of the framework introduced here is that the calculations are relatively simple, yielding a practical recipe for obtaining conditions associated with regime switches (Figure 1). The case investigated here involved only four processes, making it feasible to explore the sixteen (2^4^) dynamic regimes predicted (Figure 2 and Table S1). However, the number of potential dynamic regimes increases exponentially with the number of processes in the system (Eq. 2), so describing all possible dynamics in most systems is unfeasible. The framework’s strength lies not in enumerating all possible regimes, but in identifying the essential processes in each context and indicating the conditions under which a process becomes relevant or irrelevant throughout the dynamics. In fact, previous work has shown that the detection of critical transitions and their early warning signals depends on the length and resolution of the observational record [20–23]; the framework introduced here complements this work by indicating the observation time and resolution required for a process to be perceived at all. It can also help estimate and interpret parameter fits in dynamical systems. The explicit expressions for the switching thresholds and minimum observation times indicate the parameter ranges in which a process is irrelevant for a given observer, so fitting efforts can focus on parameters of relevant processes. This knowledge also provides information on how to characterize or engineer the dynamics of complex systems that could simplify ongoing efforts [68,87–92]. One possibility, for example, is combining the observer relevance framework introduced here with the theory to identify the strongly connected components (SCCs) in complex systems [93]. That theory builds on the fact that most dynamic variables in a complex system cannot be measured, but that graphical network analysis can identify components that help infer the dynamics of other variables. The theory introduced here argues that the graphical network depends on the observational time, implying that identifying relevant SCCs can depend on the observer’s context. A limitation of both the SCC approach and the framework introduced in this formulation is the need for prior knowledge of the potential processes involved in a system, along with their corresponding parameter ranges. Nonetheless, the concept of weights and finite constraints could be extended to assess the contribution of processes inferred in machine learning and data-driven methods [94], including the interaction strengths estimated with multivariate autoregressive state-space models [95].

Finally, the observational framework makes explicit the role of the observation scales, which is often overlooked in the analysis of dynamical systems [39,40]. The perception and interpretation of the dynamics depend on the observer’s context, such as the observational time (Figure 1). This does not imply that the dynamics are subjective: the attractors and bifurcations are properties of the system, whereas the set of processes that produce a measurable change, and therefore the dynamic regime that describes the observations, depends on the observation scales. Dynamical systems may exhibit well-defined asymptotic characteristics that can be considered universal mathematically, like the global stability introduced by the carrying capacity [96,97]. Yet observers, whose perception is finite, do not have access to such universal properties. Instead, they live in local dynamics that may lead to very different perceptions of the system, as exemplified by observers describing phage and bacteria dynamics over either a two-week or three-year period (Figure 5). The former would describe the system as periodically driven by the four standard processes in Lotka-Volterra dynamics, while the latter would perceive the long-term effect of carrying capacity, which smooths the system’s oscillations. In their contexts, both descriptions are accurate and relevant. Because the framework depends only on the interplay between system dynamics and observational limitations, it does not rely on specific features of any particular model and can therefore be implemented across diverse dynamical systems. One important advantage of the approach is the ability to capture intrinsic switches of dynamic regimes at fixed parameters due to the progression of state variables over time. This provides a solution for a problem that has been challenging to address with prior methods. Exploring its full implications across heterogeneous systems remains a direction for future work. The extrapolation of this result indicates that, in more complex scenarios, the dynamics observed may often be associated with a small number of relevant processes that dominate, providing a theoretical foundation for the prevalence of the Pareto Principle (80/20 rule), the principle of factor sparsity across systems, self-organization, and apparent simplicity in complex systems [68,89,98–101].

## CONCLUSION

The observational framework introduced here is a practical and accurate approach to identifying the dominant processes, dynamic regimes, and regime switches perceived by a finite observer of a dynamical system. It builds on the foundational empirical principle that observers are subject to finite measurements. This principle yields reference observational rates that determine when processes are relevant, shaping the attractors that drive the dynamics the observer perceives. Regime switches occur when dynamic processes become relevant or irrelevant to the observer, thereby changing the reduced model and its attractor that describes the observed dynamics. Different combinations of relevant processes generate a rich range of transients and a few quasi-stable dynamic regimes that are observable but overlooked in classic asymptotic approaches. The observational framework thus indicates that dynamic regimes and their switching thresholds depend on the observation scales. The attractors and bifurcations of the full system do not depend on these scales, yet they might not be observable. The framework thus highlights the need to use the observer’s finite perception as a cornerstone element in describing and forecasting dynamical systems.

## Supporting information

Main text and figures of the supplemental material

parameter ranges covered

Simulation data for different scenarios

## ACKNOWLEDGEMENTS

The authors thank Joan Roughgarden, Alexander V. Alekseyenko (Medical University of South Carolina), and the Biomath working group at San Diego State University, with special thanks to Jim Nulton, for their invaluable insights and fruitful discussions throughout the project. The authors also acknowledge the crucial influence of Matthew C. Witt’s Master’s Thesis, carried out under the supervision of Antoni Luque at San Diego State University [102], and the theoretical inspiration from several key references [3,9,28,88]. Regarding the use of AI, the authors used Claude (Anthropic) to assist with code development for the robustness analysis of the Boolean model and to edit the manuscript during the revision process. The authors reviewed, tested, and verified all AI-assisted code, analyses, and text and take full responsibility for the content.

## DATA AVAILABILITY STATEMENT

All code and data underlying the findings are publicly available. The simulation code (Jupyter notebook), example scenarios, robustness-analysis scripts, simulation outputs, and figure files are archived on Zenodo (release v0.2.3-alpha, https://doi.org/10.5281/zenodo.24014219) [54] and maintained on GitHub (https://github.com/luquelab/transient-dynamics) under the MIT license.

## AUTHOR CONTRIBUTION STATEMENT

Conceptualization: AL. Data curation: SC-L, LJC, MCW. Formal analysis: SC-L, LJC, MCW, AL. Funding acquisition: AL, FLR. Investigation: SC-L, LJC, MCW. Methodology: SC-L, LJC, MCW, AL. Project administration: AL. Resources: AL, FLR. Software: SC-L, LJC, MCW, AL. Supervision: AL, FLR. Validation: LJC, AL. Visualization: SC-L, LJC, AL. Writing – original draft: SC-L. Writing – review & editing: AL, LJC, MCW, FLR.

## SUPPORTING INFORMATION

**S1 Text. Supplemental Material.** Asymptotic analysis of the sixteen dynamic regimes (S1; Tables S1–S4), model parametrization (S2), simulated dynamics of Scenarios 2 and 4 (S3; Figures S1 and S2), summary statistics of the relative error of the adaptive Boolean model (S4; Table S5), and robustness analysis of the adaptive Boolean model across the sampled parameter space and observer tolerances (S5; Figures S3–S6 and Table S6).

**S1 Dataset. Parameter ranges.** Literature values used to build the ranges of the parameters, concentrations, and observational times in Table 1 (supplementary_file_parameter_ranges.xlsx).

**S2 Dataset. Simulation outputs.** Simulated populations, weights, and relevant processes for Scenarios 1–4, including the alternative initial conditions and the model with carrying capacity (supplementary_file_simulations_outputs.zip).

