## Supplementary material for "Emerging transient dynamics at finite observation scales": Main text and figures of the supplemental material

### S1 Text. Supplemental Material

#### S1. Asymptotic Analysis

The tables below present the asymptotic stability analysis for the 16 dynamic regimes associated with the combination of relevant processes in the Lotka-Volterra phage dynamic system (see the differential equation in Figure 2a). As described in the results section, the predicted dynamics were grouped in four scenarios: relevant growth and irrelevant decay (Scenario 1), irrelevant growth and relevant decay (Scenario 2), relevant growth and relevant decay (Scenario 3), and irrelevant growth and irrelevant decay (Scenario 4). In each scenario, four possible dynamic regimes were identified, depending on whether the predation and burst processes were relevant or irrelevant, which in turn depended on the dynamic variables. The asymptotic analysis is provided below in a table associated, respectively, with each scenario: Table S1 (Scenario 1), Table S2 (Scenario 2), Table S3 (Scenario 3), and Table S4 (Scenario 4). The asymptotic analysis for each subset of (relevant) processes was carried out in two steps.

The first step of the analysis determined if there were any equilibrium conditions consistent with the relevant processes in that scenario. For example, the dynamic regimes, $DR_{I}$, $DR_{II}$, and $DR_{III}$ in Table S1 had an apparent mathematical solution for the equilibrium of bacteria (rate zero) for $B* = 0$ as a fixed point. However, this solution was incompatible with the growth process being relevant because if $B = 0$, the weight for the growth would be $w_g = 0 < w_{th} = 1$. This exemplifies a common situation in analyzing the sixteen dynamical subsystems. In other cases, such as $DR_{IV}$ and $DR_{V}$, the equilibrium-imposed constraints in the parameter values compatible with relevant processes require the equilibrium concentration to be above the corresponding switching threshold: $P^* > P_c = \alpha/(\tau \cdot a)$ for $DR_{IV}$ and $B^* = m/(c \cdot a) > B_c = \alpha/(\tau \cdot c \cdot a)$ for $DR_{V}$.

The second step was to determine the dynamics' stability and attractors. For regimes with fixed points compatible with the relevant processes, the eigenvalues of the Jacobian matrix were obtained and evaluated. For regimes without fixed points, the attractors were obtained based on the directionality of the asymptotic limit. For instance, $DR_{I}$ is unstable because the bacterial concentration is growing exponentially. $DR_{IV}$ is unstable. The phage concentration remains approximately constant, yet any perturbation above or below the fixed point, $P^*=r/a$, will either decrease or increase the bacterial population. It will decrease, $B \to 0$, when the predation process is stronger than the growth process, that is, $P > r/a$, and increase, $B \to \infty$, when the growth process is stronger than the predation process, $P < r/a$. Either scenario requires the phage concentration to be above the switching threshold, $P > P_c = \alpha/(\tau \cdot a)$, so the predation term is relevant.

**Table S1: Asymptotic analysis of dynamic regimes in Scenario 1 (relevant growth, irrelevant decay).** Each dynamic regime, DR, is labeled with a Roman numeral as in the main text (Figure 2b) and is described by the reduced model that contains only its relevant processes. The equilibria, eigenvalues, stability, and attractors refer to that reduced model, not to the full system. A regime had a valid equilibrium only if its fixed points were compatible with its relevant processes; the same applied to the eigenvalues of the Jacobian matrix. The stability of each regime was determined based on the attractors.

| Dynamic regime | Reduced model | Equilibrium | Eigenvalues | Attractors | Stability |
|---|---|---|---|---|---|
| $DR_I$ | $\frac{dB}{dt} = rB$ <br> $\frac{dP}{dt} = 0$ | | | $B \to \infty$ <br> $P \sim P_0$ | Unstable |
| $DR_{II}$ | $\frac{dB}{dt} = rB$ <br> $\frac{dP}{dt} = caBP$ | | | $B \to \infty$ <br> $P \to \infty$ | Unstable |
| $DR_{III}$ | $\frac{dB}{dt} = rB - aBP$ <br> $\frac{dP}{dt} = caBP$ | | | $B \to 0$ <br> $P \to \infty$ | Unstable |
| $DR_{IV}$ | $\frac{dB}{dt} = rB - aBP$ <br> $\frac{dP}{dt} = 0$ | $B^* = B_0$ <br> $P^* = \frac{r}{a} > P_c$ | $\lambda_1 = \lambda_2 = 0$ | $B \to 0 \quad P > \frac{r}{a}$ <br> $B \to \infty \quad P < \frac{r}{a}$ <br> $P \sim P_0$ | Unstable |

In Table S1, dynamic regimes $DR_I$, $DR_{II}$, and $DR_{III}$ did not include the trivial fixed point where $B = 0$ because this value is incompatible with the growth process being relevant in the dynamics.

Consequently, the eigenvalues of the Jacobian matrix could not be calculated. Dynamic regime $DR_{IV}$ had a valid equilibrium if $P^*$ was higher than the switching threshold of the predation process, $P_c$. The bacteria had two different attractors depending on the phage concentrations.

**Table S2: Asymptotic analysis of dynamic regimes in Scenario 2 (irrelevant growth, relevant decay).** Each dynamic regime, DR, is labeled with a Roman numeral as in the main text (Figure 2b) and is described by the reduced model that contains only its relevant processes. The equilibria, eigenvalues, stability, and attractors refer to that reduced model, not to the full system. A regime had a valid equilibrium only if its fixed points were compatible with its relevant processes; the same applied to the eigenvalues of the Jacobian matrix. The stability of each regime was determined based on the attractors.

| Dynamic regime | Dynamics | Equilibrium | Eigenvalues | Attractors | Stability |
|---|---|---|---|---|---|
| $DR_{V}$ | $\frac{dB}{dt} = 0$<br>$\frac{dP}{dt} = caBP - mP$ | $B^* = \frac{m}{ca} > B_c$<br>$P^* = P_0$ | $\lambda_1 = \lambda_2 = 0$ | $B \sim B_0$<br>$P \to 0 \quad B < \frac{m}{ca}$<br>$P \to \infty \quad B > \frac{m}{ca}$ | Unstable |
| $DR_{VI}$ | $\frac{dB}{dt} = -aBP$<br>$\frac{dP}{dt} = caBP - mP$ | | | $B \to 0$<br>$P \to 0$ | Unstable |
| $DR_{VII}$ | $\frac{dB}{dt} = -aBP$<br>$\frac{dP}{dt} = -mP$ | | | $B \to 0$<br>$P \to 0$ | Unstable |
| $DR_{VIII}$ | $\frac{dB}{dt} = 0$<br>$\frac{dP}{dt} = -mP$ | | | $B \sim B_0$<br>$P \to 0$ | Unstable |

In Table S2, only dynamic regime $DR_V$ had an equilibrium if $B^* > B_c$, where $B_c = \alpha/(\tau\ c\ a)$. The phage concentration had two different attractors depending on the bacterial concentrations. All other regimes did not have equilibria because their fixed points were incompatible with one or more of their relevant processes. All four regimes were unstable, as shown by the attractors in the last column.

**Table S3: Asymptotic analysis of dynamic regimes in Scenario 3 (relevant growth and decay).** Each dynamic regime, DR, is labeled with a Roman numeral as in the main text (Figure 2b) and is described by the reduced model that contains only its relevant processes. The equilibria, eigenvalues, stability, and attractors refer to that reduced model, not to the full system. A regime had a valid equilibrium only if its fixed points were compatible with its relevant processes; the same applied to the eigenvalues of the Jacobian matrix. The stability of each regime was determined based on the attractors.

| Dynamic regime | Dynamics | Equilibrium | Eigenvalues | Attractors | Stability |
|---|---|---|---|---|---|
| $DR_{IX}$ | $\frac{dB}{dt} = rB$ <br> $\frac{dP}{dt} = caBP - mP$ | | | $B \to \infty$ <br> $P \to \infty$ | Unstable |
| $DR_{X}$ | $\frac{dB}{dt} = rB - aBP$ <br> $\frac{dP}{dt} = caBP - mP$ | $B^* = m/ca$ <br> $P^* = r/a$ | $\lambda_1 = i\sqrt{mr}$ <br> $\lambda_2 = -i\sqrt{mr}$ | $\overline{B} \to B^*$ <br> $\overline{P} \to P^*$ | Center |
| $DR_{XI}$ | $\frac{dB}{dt} = rB - aBP$ <br> $\frac{dP}{dt} = -mP$ | | | $B \to \infty$ <br> $P \to 0$ | Unstable |
| $DR_{XII}$ | $\frac{dB}{dt} = rB$ <br> $\frac{dP}{dt} = -mP$ | | | $B \to \infty$ <br> $P \to 0$ | Unstable |

In Table S3, only dynamic regime $DR_X$ (all processes relevant, that is, the full Lotka-Volterra model) had an equilibrium. The stability analysis predicted the well-known center around the fixed points $B^* = m/ca$ and $P^* = r/a$. All other regimes were unstable.

**Table S4: Asymptotic analysis of dynamic regimes in Scenario 4 (irrelevant growth and decay).** Each dynamic regime, DR, is labeled with a Roman numeral as in the main text (Figure 2b) and is described by the reduced model that contains only its relevant processes. The equilibria, eigenvalues, stability, and attractors refer to that reduced model, not to the full system. A regime had a valid equilibrium only if its fixed points were compatible with its relevant processes; the same applied to the eigenvalues of the Jacobian matrix. The stability of each regime was determined based on the attractors.

| Dynamic regime | Dynamics | Equilibrium | Eigenvalues | Attractors | Stability |
|---|---|---|---|---|---|
| $DR_{XIII}$ | $\frac{dB}{dt} = 0$<br>$\frac{dP}{dt} = caBP$ | | | $B \to 0$<br>$P \to \infty$ | Unstable |
| $DR_{XIV}$ | $\frac{dB}{dt} = -aBP$<br>$\frac{dP}{dt} = caBP$ | | | $B \to 0$<br>$P \to \infty$ | Unstable |
| $DR_{XV}$ | $\frac{dB}{dt} = -aBP$<br>$\frac{dP}{dt} = 0$ | | | $B \to 0$<br>$P \sim P_0$ | Unstable |
| $DR_{XVI}$ | $\frac{dB}{dt} = 0$<br>$\frac{dP}{dt} = 0$ | $B^* = B_0$<br>$P^* = P_0$ | $\lambda_1 = \lambda_2 = 0$ | $B \sim B_0$<br>$P \sim P_0$ | Quasi-stable |

Table S4 has only one quasi-stable regime. It corresponds to dynamic regime $DR_{XVI}$, where no process is relevant, and the populations remain quasi-static. The remaining regimes in this scenario were unstable and did not have valid equilibria.

#### S2. Model Parametrization

The values for the model's parametrization are included in the parameter-ranges supplementary file, "supplementary_file_parameter_ranges.xlsx" (S1 Dataset). The references are cited in the main text. The file consists of two tabs: "Descriptors" provides descriptions for each column; "Table_values" contains the values. These are the columns included: Reference, DOI, source of data, growth rate range (and units), decay rate range (and units), adsorption rate (and units), burst size (and units), phage abundance range (and units), bacterial abundance range (and units), and observational time range (and units).

#### S3. Simulated Dynamics

The data obtained from the numerical simulations are available in the supplementary file "supplementary_file_simulations_outputs.zip" (S2 Dataset). The decompressed folder includes a dictionary text file ("data_output_dict.md") describing the data files. Briefly, the data files include the simulations for scenarios 1, 2, 3, and 4 (see main text), including those for alternative initial conditions or processes (like the presence of the carrying capacity).

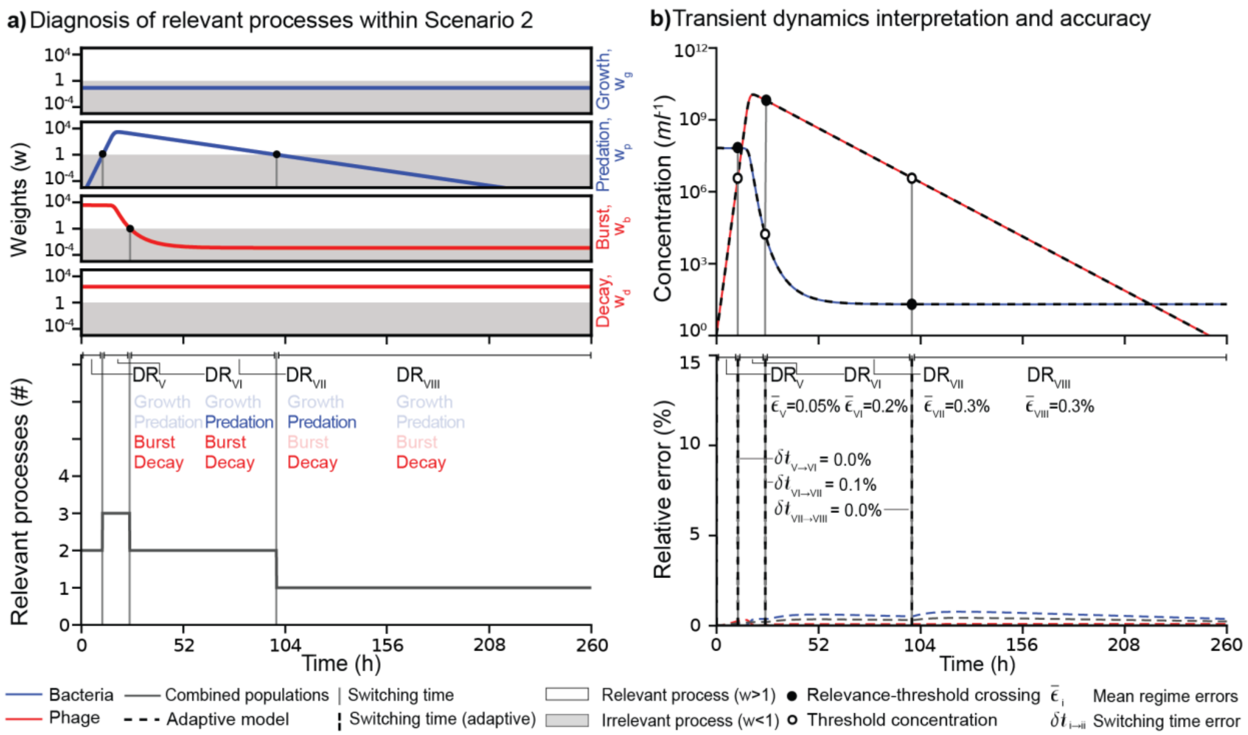


**Figure S1. Transient dynamics for Scenario 2.** a-b) Weights, relevant processes, dynamics, and relative errors for Scenario 2 (irrelevant growth rate and relevant decay rate). The symbols in each panel are analogous to those displayed and described in Figure 3. The parameters used in this simulated scenario are in Table 1.

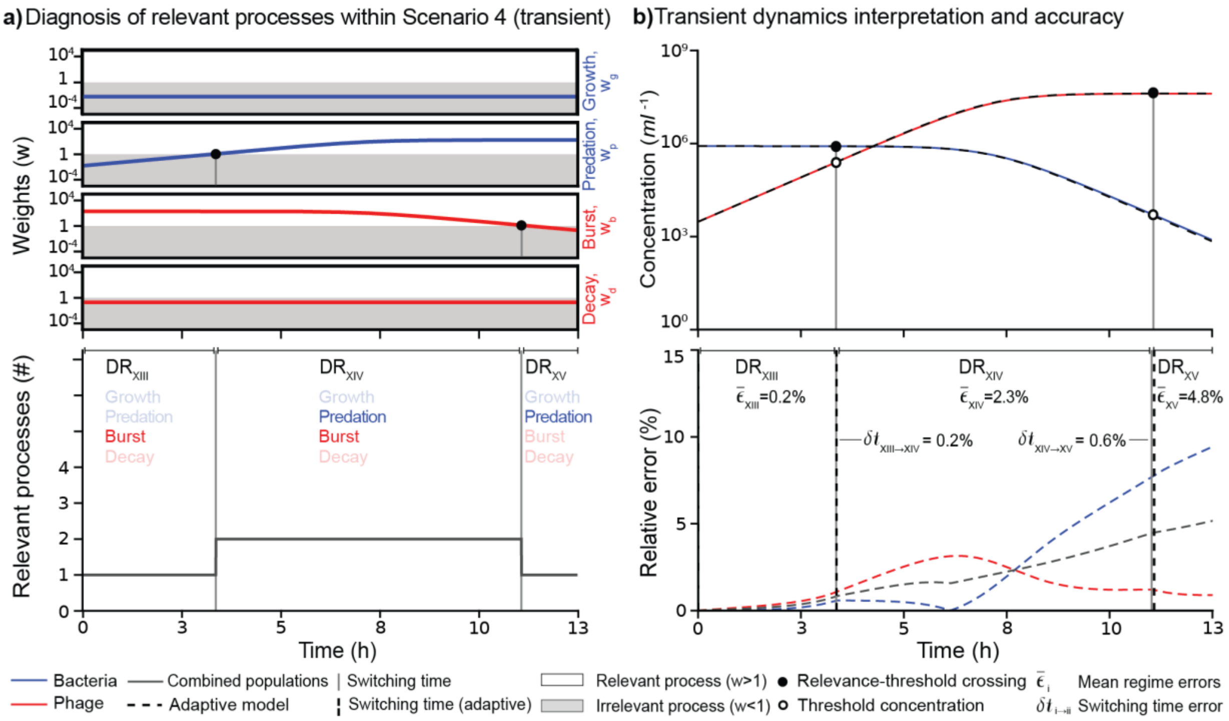


**Figure S2. Transient dynamics for Scenario 4.** a-b) Weights, relevant processes, dynamics, and relative errors for Scenario 4 (irrelevant growth rate and irrelevant decay rate) for the transient initial condition. The symbols in each panel are analogous to those displayed and described in Figure 3. The parameters for this simulated scenario are in Table 1.

#### S4. Relative Error

Statistical summary of the relative error percentages between the adaptive Boolean and full models across various scenarios and regimes. The columns include the scenario (parameters in Table 1), the regimes (Figure 2 and Tables S1-S4), the agent (phage, bacteria, and combined average), and the statistical summary of the relative error, that is, minimum (0th percentile), Q1 (25th percentile), median (50th percentile), mean, Q3 (75th percentile), and maximum (100th percentile).

**Table S5. Summary statistics of the error analysis.** The columns below display the summary statistics of the relative error (as percentages, %) of the Boolean model (which only simulates relevant processes) for the full dynamics model. The rows are organized by Scenario and Regime, following the notation of Figure 2. The column 'Agent' indicates a row that focuses on either the error in the phage population, the bacterial population, or the average error per time step for both (Combined). The summary statistics include the minimum value (Minimum), quartile 1 (Q1), median (Median), mean (Mean), quartile 3 (Q3), and maximum value for each scenario, regime, and population analyzed.

| Scenario | Regime (DR) | Agent | Minimum | Q1 | Median | Mean | Q3 | Maximum |
|---|---|---|---|---|---|---|---|---|
| 1 | I | Phage | 0.00E+00 | 1.45E-02 | 6.69E-02 | 1.05E-01 | 1.75E-01 | 3.66E-01 |
| | | Bacteria | 0.00E+00 | 1.30E-02 | 2.60E-02 | 2.60E-02 | 3.90E-02 | 5.21E-02 |
| | | Combined | 0.00E+00 | 1.37E-02 | 4.64E-02 | 6.54E-02 | 1.07E-01 | 2.09E-01 |
| | II | Phage | 3.32E-06 | 1.32E-01 | 2.69E-01 | 3.86E-01 | 5.12E-01 | 1.67E+00 |
| | | Bacteria | 5.21E-02 | 1.03E-01 | 1.57E-01 | 1.81E-01 | 2.29E-01 | 5.60E-01 |
| | | Combined | 6.29E-02 | 1.26E-01 | 1.89E-01 | 2.84E-01 | 3.70E-01 | 1.11E+00 |
| | III | Phage | 1.15E-01 | 4.82E-01 | 2.25E+00 | 2.09E+00 | 3.35E+00 | 4.07E+00 |
| | | Bacteria | 9.03E-05 | 5.21E-01 | 5.60E-01 | 2.21E+00 | 4.83E+00 | 5.67E+00 |
| | | Combined | 1.11E+00 | 1.71E+00 | 2.11E+00 | 2.15E+00 | 2.66E+00 | 2.90E+00 |
| | IV | Phage | 8.14E-02 | 1.63E-01 | 2.91E-01 | 2.95E-01 | 4.20E-01 | 5.49E-01 |
| | | Bacteria | 5.68E+00 | 6.51E+00 | 8.18E+00 | 8.76E+00 | 1.07E+01 | 1.41E+01 |
| | | Combined | 2.90E+00 | 3.33E+00 | 4.23E+00 | 4.53E+00 | 5.57E+00 | 7.32E+00 |
| | I-IV | Phage | 0.00E+00 | 1.23E-01 | 2.75E-01 | 5.96E-01 | 5.16E-01 | 4.07E+00 |
| | | Bacteria | 0.00E+00 | 1.04E-01 | 2.37E-01 | 2.57E+00 | 5.59E+00 | 1.41E+01 |
| | | Combined | 0.00E+00 | 1.40E-01 | 4.24E-01 | 1.58E+00 | 2.85E+00 | 7.32E+00 |
| 2 | V | Phage | 0.00E+00 | 1.76E-02 | 7.02E-02 | 9.17E-02 | 1.57E-01 | 2.52E-01 |

| | | | | | | | | |
|---|---|---|---|---|---|---|---|---|
| | | **Bacteria** | 0.00E+00 | 8.49E-03 | 1.62E-02 | 1.55E-02 | 2.32E-02 | 2.67E-02 |
| | | **Combined** | 0.00E+00 | 1.31E-02 | 4.37E-02 | 5.36E-02 | 9.13E-02 | 1.31E-01 |
| | **VI** | **Phage** | 3.10E-06 | 1.92E-03 | 7.90E-02 | 1.33E-01 | 2.74E-01 | 3.10E-01 |
| | | **Bacteria** | 2.83E-04 | 1.23E-02 | 2.85E-01 | 2.02E-01 | 3.51E-01 | 3.56E-01 |
| | | **Combined** | 1.29E-01 | 1.57E-01 | 1.75E-01 | 1.68E-01 | 1.80E-01 | 1.83E-01 |
| | **VII** | **Phage** | 4.18E-04 | 6.88E-02 | 7.02E-02 | 6.80E-02 | 7.11E-02 | 7.19E-02 |
| | | **Bacteria** | 3.45E-01 | 5.15E-01 | 5.60E-01 | 5.46E-01 | 5.94E-01 | 6.07E-01 |
| | | **Combined** | 1.73E-01 | 2.93E-01 | 3.15E-01 | 3.07E-01 | 3.32E-01 | 3.38E-01 |
| | **VIII** | **Phage** | 7.19E-02 | 7.38E-02 | 7.56E-02 | 7.56E-02 | 7.74E-02 | 7.92E-02 |
| | | **Bacteria** | 3.56E-01 | 4.85E-01 | 6.01E-01 | 5.90E-01 | 7.07E-01 | 7.61E-01 |
| | | **Combined** | 2.18E-01 | 2.81E-01 | 3.38E-01 | 3.33E-01 | 3.90E-01 | 4.17E-01 |
| | **V-VIII** | **Phage** | 0.00E+00 | 7.09E-02 | 7.39E-02 | 7.72E-02 | 7.68E-02 | 3.10E-01 |
| | | **Bacteria** | 0.00E+00 | 4.65E-01 | 5.63E-01 | 5.33E-01 | 6.43E-01 | 7.61E-01 |
| | | **Combined** | 0.00E+00 | 2.70E-01 | 3.18E-01 | 3.05E-01 | 3.59E-01 | 4.17E-01 |
| **3 (Transient)** | **IX** | **Phage** | 1.47E-14 | 2.35E-02 | 3.46E-01 | 5.25E-01 | 5.66E-01 | 5.79E+00 |
| | | **Bacteria** | 0.00E+00 | 1.47E-01 | 5.63E-01 | 4.23E-01 | 7.25E-01 | 7.82E-01 |
| | | **Combined** | 7.37E-15 | 8.63E-02 | 4.53E-01 | 4.74E-01 | 6.21E-01 | 3.28E+00 |
| | **X** | **Phage** | 3.02E-03 | 4.28E-01 | 1.54E+00 | 2.90E+00 | 5.79E+00 | 1.02E+01 |
| | | **Bacteria** | 2.71E-03 | 4.20E-01 | 1.65E+00 | 2.76E+00 | 5.11E+00 | 9.32E+00 |
| | | **Combined** | 7.87E-01 | 1.18E+00 | 2.32E+00 | 2.83E+00 | 4.64E+00 | 5.55E+00 |
| | **XI** | **Phage** | 1.83E-01 | 2.40E-01 | 2.57E-01 | 4.13E-01 | 7.98E-01 | 8.14E-01 |
| | | **Bacteria** | 3.88E-04 | 3.00E-01 | 3.81E-01 | 6.78E-01 | 4.93E-01 | 5.71E+00 |
| | | **Combined** | 1.19E-01 | 2.91E-01 | 3.33E-01 | 5.46E-01 | 5.74E-01 | 3.23E+00 |
| | **XII** | **Phage** | 4.17E-01 | 4.48E-01 | 4.83E-01 | 4.87E-01 | 5.24E-01 | 5.72E-01 |
| | | **Bacteria** | 3.90E-01 | 4.40E-01 | 4.81E-01 | 4.77E-01 | 5.17E-01 | 5.48E-01 |
| | | **Combined** | 4.04E-01 | 4.44E-01 | 4.82E-01 | 4.82E-01 | 5.21E-01 | 5.21E-01 |
| | **IX-XII** | **Phage** | 1.47E-14 | 2.15E-01 | 3.08E-01 | 7.21E-01 | 6.24E-01 | 1.02E+01 |
| | | **Bacteria** | 0.00E+00 | 1.52E-01 | 3.87E-01 | 7.51E-01 | 7.26E-01 | 9.32E+00 |
| | | **Combined** | 7.37E-15 | 2.27E-01 | 4.36E-01 | 7.36E-01 | 6.31E-01 | 5.55E+00 |
| **3 (Quasi-stable)** | **X** | **Phage** | 0.00E+00 | 0.00E+00 | 0.00E+00 | 0.00E+00 | 0.00E+00 | 0.00E+00 |
| | | **Bacteria** | 0.00E+00 | 0.00E+00 | 0.00E+00 | 0.00E+00 | 0.00E+00 | 0.00E+00 |
| | | **Combined** | 0.00E+00 | 0.00E+00 | 0.00E+00 | 0.00E+00 | 0.00E+00 | 0.00E+00 |
| **3 (Quasi-stable/ Carrying Capacity)** | **Carrying capacity Irrelevant** $\tau$ = 300 h | **Phage** | 0.00E+00 | 2.07E-01 | 4.76E-01 | 8.60E-01 | 1.23E+00 | 5.10E+00 |
| | | **Bacteria** | 0.00E+00 | 3.38E-01 | 8.22E-01 | 1.09E+00 | 1.70E+00 | 3.69E+00 |
| | | **Combined** | 0.00E+00 | 3.33E-01 | 7.78E-01 | 9.75E-01 | 1.48E+00 | 3.66E+00 |
| | | **Phage** | 0.00E+00 | 0.00E+00 | 0.00E+00 | 0.00E+00 | 0.00E+00 | 0.00E+00 |

| | | | | | | | | |
|---|---|---|---|---|---|---|---|---|
| | **Carrying capacity relevant** $\tau$ = 30,000 h | **Bacteria** | 0.00E+00 | 0.00E+00 | 0.00E+00 | 0.00E+00 | 0.00E+00 | 0.00E+00 |
| | | **Combined** | 0.00E+00 | 0.00E+00 | 0.00E+00 | 0.00E+00 | 0.00E+00 | 0.00E+00 |
| **4 (Transient)** | **XIII** | **Phage** | 1.52E-14 | 1.36E-01 | 2.98E-01 | 3.65E-01 | 5.47E-01 | 1.07E+00 |
| | | **Bacteria** | 0.00E+00 | 1.16E-02 | 5.25E-02 | 1.21E-01 | 1.82E-01 | 5.82E-01 |
| | | **Combined** | 7.58E-15 | 7.37E-02 | 1.75E-01 | 2.43E-01 | 3.65E-01 | 8.24E-01 |
| | **XIV** | **Phage** | 1.07E+00 | 1.29E+00 | 1.93E+00 | 2.03E+00 | 2.75E+00 | 3.15E+00 |
| | | **Bacteria** | 4.54E-04 | 5.22E-01 | 1.38E+00 | 2.62E+00 | 4.87E+00 | 7.70E+00 |
| | | **Combined** | 8.25E-01 | 1.57E+00 | 2.08E+00 | 2.32E+00 | 3.11E+00 | 4.46E+00 |
| | **XV** | **Phage** | 8.88E-01 | 8.92E-01 | 9.31E-01 | 9.73E-01 | 1.03E+00 | 1.23E+00 |
| | | **Bacteria** | 7.70E+00 | 8.20E+00 | 8.64E+00 | 8.62E+00 | 9.04E+00 | 9.44E+00 |
| | | **Combined** | 4.46E+00 | 4.62E+00 | 4.78E+00 | 4.79E+00 | 4.97E+00 | 5.16E+00 |
| | **XIII-XV** | **Phage** | 1.52E-14 | 8.76E-01 | 1.22E+00 | 1.46E+00 | 2.24E+00 | 3.15E+00 |
| | | **Bacteria** | 0.00E+00 | 2.04E-01 | 5.76E-01 | 2.66E+00 | 5.30E+00 | 9.44E+00 |
| | | **Combined** | 7.58E-15 | 6.51E-01 | 1.63E+00 | 2.06E+00 | 3.29E+00 | 5.16E+00 |
| **4 (Quasi-static)** | **XVI** | **Phage** | 0.00E+00 | 2.12E-04 | 8.49E-04 | 1.13E-03 | 1.91E-03 | 3.39E-03 |
| | | **Bacteria** | 0.00E+00 | 8.71E-02 | 1.74E-01 | 1.74E-01 | 2.62E-01 | 3.49E-01 |
| | | **Combined** | 0.00E+00 | 4.37E-02 | 8.76E-02 | 8.78E-02 | 1.32E-01 | 1.76E-01 |

#### S5. Robustness analysis

Figures S3–S6 show the robustness analysis of the adaptive Boolean model described in the Methods of the main text.

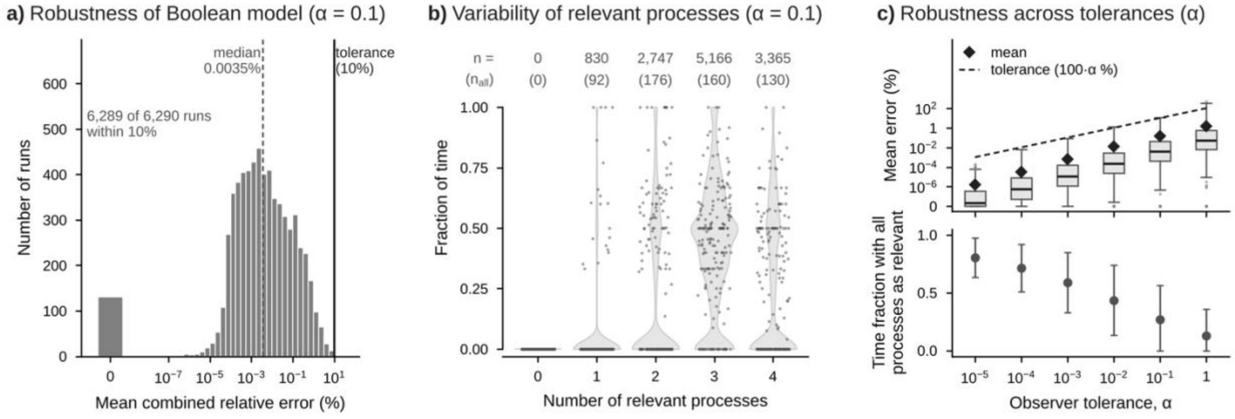


**Figure S3. Robustness analysis of the adaptive Boolean model for stopped simulations.** Latin hypercube sampling (log-uniform) of the parameters, initial concentrations, and observational time of the phage–bacteria (Lotka–Volterra) model; only simulations that were stopped before the end of the observational window (Methods of the main text) are shown, and both models are compared up to the stop time (n = 6,290 for α = 0.1). **a)** Histogram of the mean combined relative error (%) between the adaptive Boolean and full models at α = 0.1, calculated over the simulated portion of each trajectory; the bar labeled 0 counts runs with zero error (all processes relevant throughout), the dashed line marks the median, and the solid black line marks the observer tolerance (10%, corresponding to α = 0.1). **b)** Fraction of the simulated time that each run spent with 0 to 4 relevant processes (weight $w \geq 1$) at α = 0.1; violins show the distribution across runs, gray points are a random sample of 200 runs, the numbers on top (n) are the runs that spent any time with that number of relevant processes, and the numbers in parentheses ($n_{all}$) are the runs that spent all of the simulated time with that number. **c)** Top, box plots of the mean combined relative error for $\alpha = 10^{-5}$ to 1 (box, interquartile range, IQR; line, median; whiskers, 1.5 × IQR; points, values beyond the whiskers), with the mean (◆) and the observer tolerance, 100·α % (dashed line); zero errors are drawn at the bottom of the axis, labeled 0. Bottom, mean ± standard deviation of the fraction of simulated time with all 4 processes relevant. Extreme outliers (Tukey's far-out test, above Q3 + 3 × IQR of $\log_{10}$ error) are excluded from the statistics and, when present, shown in ochre.

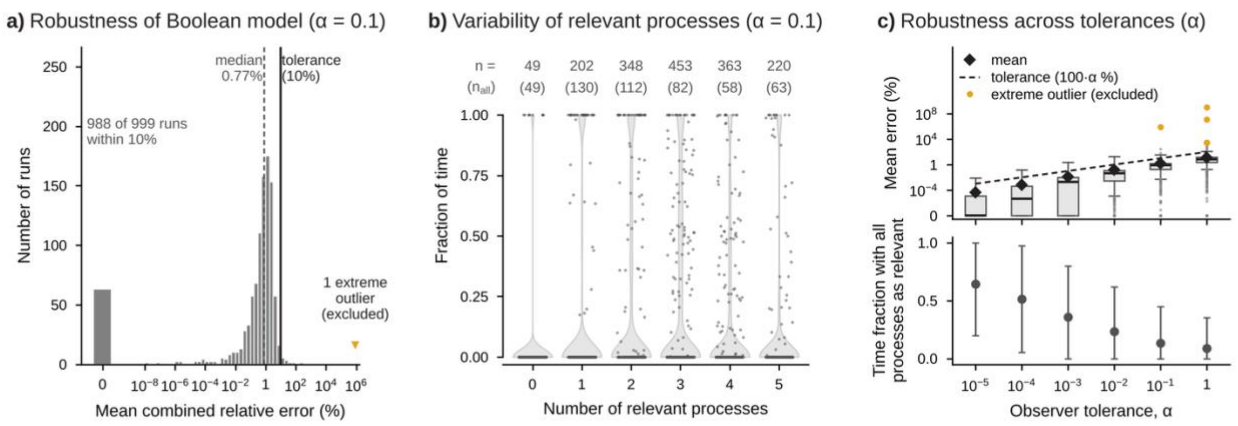


**Figure S4. Robustness analysis of the adaptive Boolean model with carrying capacity.** Latin hypercube sampling (log-uniform) of the parameters, initial concentrations, and observational time of the phage–bacteria model with bacterial carrying capacity; only simulations that remained within the physical bounds for the full observational window are shown (n = 1,000 for α = 0.1). **a)** Histogram of the mean combined relative error (%) between the adaptive Boolean and full models at α = 0.1; the bar labeled 0 counts runs with zero error (all processes relevant throughout), the dashed line marks the median, and the solid black line marks the observer tolerance (10%, corresponding to α = 0.1). **b)** Fraction of the observational time that each run spent with 0 to 5 relevant processes (weight w ≥ 1) at α = 0.1; violins show the distribution across runs, gray points are a random sample of 200 runs, the numbers on top (n) are the runs that spent any time with that number of relevant processes, and the numbers in parentheses ($n_{all}$) are the runs that spent all of the observational time with that number. **c)** Top, box plots of the mean combined relative error for α = $10^{-5}$ to 1 (box, interquartile range, IQR; line, median; whiskers, 1.5 × IQR; points, values beyond the whiskers), with the mean (◆) and the observer tolerance, 100·α % (dashed line); zero errors are drawn at the bottom of the axis, labeled 0. Bottom, mean ± standard deviation of the fraction of observational time with all 5 processes relevant. Extreme outliers (Tukey's far-out test, above Q3 + 3 × IQR of $\log_{10}$ error) are excluded from the statistics and, when present, shown in ochre.

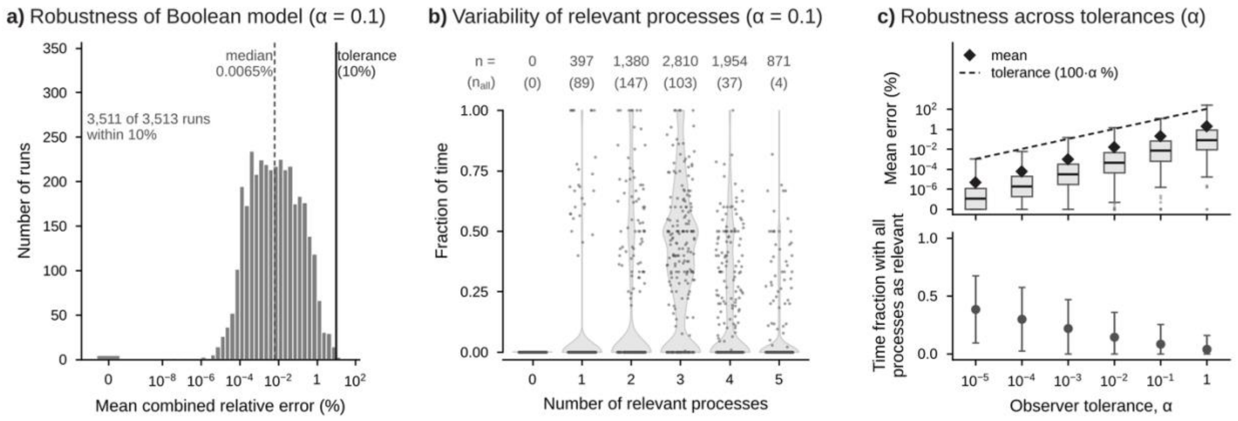


**Figure S5. Robustness analysis of the adaptive Boolean model with carrying capacity for stopped simulations.** Latin hypercube sampling (log-uniform) of the parameters, initial concentrations, and observational time of the phage–bacteria model with bacterial carrying capacity; only simulations that were stopped before the end of the observational window (Methods of the main text) are shown, and both models are compared up to the stop time (n = 3,513 for α = 0.1). **a)** Histogram of the mean combined relative error (%) between the adaptive Boolean and full models at α = 0.1, calculated over the simulated portion of each trajectory; the bar labeled 0 counts runs with zero error (all processes relevant throughout), the dashed line marks the median, and the solid black line marks the observer tolerance (10%, corresponding to α = 0.1). **b)** Fraction of the simulated time that each run spent with 0 to 5 relevant processes (weight w ≥ 1) at α = 0.1; violins show the distribution across runs, gray points are a random sample of 200 runs, the numbers on top (n) are the runs that spent any time with that number of relevant processes, and the numbers in parentheses ($n_{all}$) are the runs that spent all of the simulated time with that number. **c)** Top, box plots of the mean combined relative error for $\alpha = 10^{-5}$ to 1 (box, interquartile range, IQR; line, median; whiskers, 1.5 × IQR; points, values beyond the whiskers), with the mean (◆) and the observer tolerance, 100·α % (dashed line); zero errors are drawn at the bottom of the axis, labeled 0. Bottom, mean ± standard deviation of the fraction of simulated time with all 5 processes relevant. Extreme outliers (Tukey's far-out test, above Q3 + 3 × IQR of $\log_{10}$ error) are excluded from the statistics and, when present, shown in ochre.

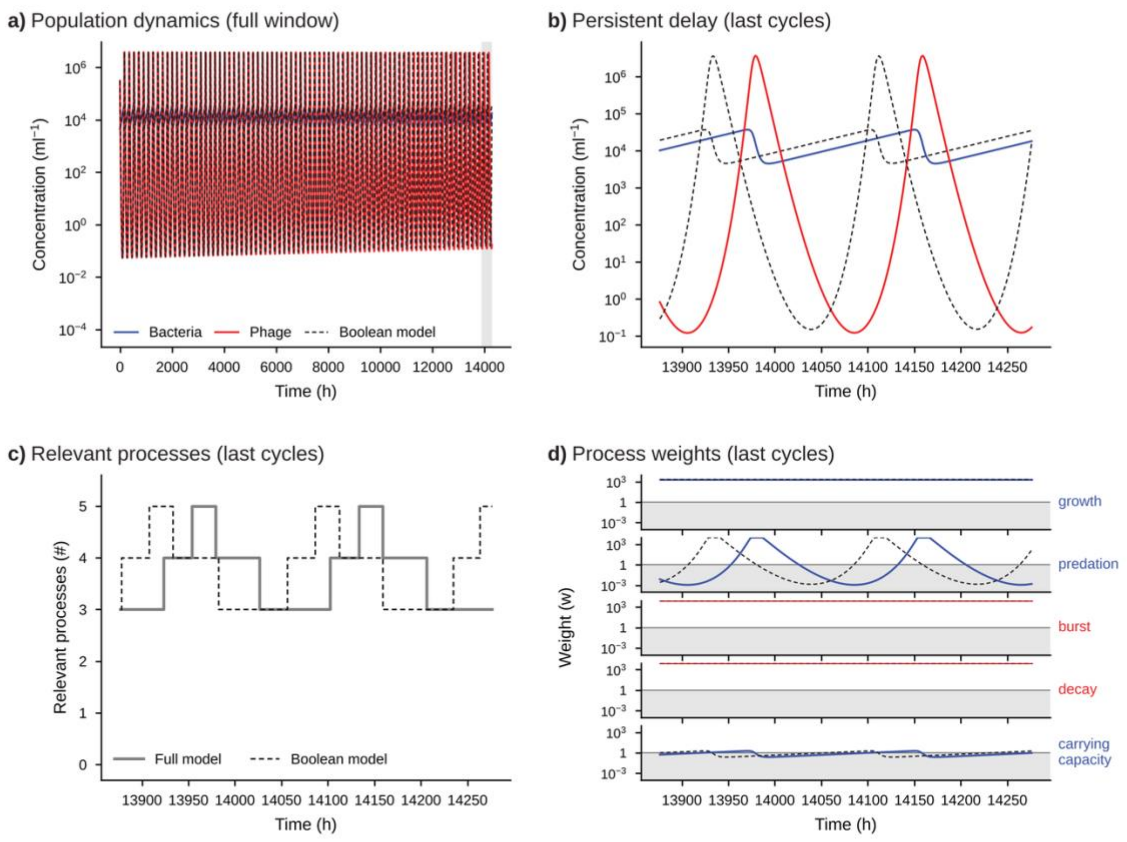


**Figure S6. Example of the extreme outlier with carrying capacity: persistent delay.** Simulation 178 of the robustness analysis of the model with carrying capacity at α = 0.1 (mean combined relative error 8.6 × $10^5$%; extreme outlier). The Boolean model gradually leads the full model by up to about 47 h at the end of the window, so phage troughs near $10^{-1}$ ml$^{-1}$ are compared with peaks near $4 \cdot 10^6$ ml$^{-1}$; the error reflects this phase shift rather than different dynamics. Blue, bacteria and bacterial processes; red, phage and phage processes; dashed black, adaptive Boolean model. **a)** Bacteria and phage concentrations over the full observational window for the full (solid) and Boolean (dashed) models; the shaded interval marks the last cycles shown in b–d. **b)** Concentrations over the last cycles, showing the delay accumulated by the Boolean model. **c)** Number of relevant processes (w ≥ 1) in the full (solid gray) and Boolean (dashed) models. **d)** Weight of each process in the full (colored) and Boolean (dashed) models; the gray band marks irrelevant weights (w < 1), and the solid line marks the threshold w = 1.

**Table S6. Robustness analysis of the adaptive Boolean model across observer tolerances.** Latin hypercube sampling (log-uniform) of the model parameters and observational time within the Table 1 ranges and of the initial concentrations within $10^3$–$10^{12}$ bacteria ml$^{-1}$ and $10^3$–$10^{14}$ phages ml$^{-1}$ (16,000 samples without and 8,000 with carrying capacity; K = $10^4$–$10^{13}$ ml$^{-1}$ with $B_0 \leq K$). For each α, the adaptive Boolean model was compared with the full model over the observational window. Error is the mean combined relative error, $\frac{1}{2}(|\Delta B|/B + |\Delta P|/P) \times 100$, averaged over time; the maximum error is its largest value over time. A run is within tolerance when the error is at most 100·α %. Samples: completed runs covered the full observational window; stopped runs ended early because a population fell below $10^{-3}$ ml$^{-1}$ or exceeded $10^{15}$ bacteria or $10^{18}$ phages ml$^{-1}$ and were compared up to the stop time; a further 8,021 samples without and 2,980 with carrying capacity stopped before two time points could be compared and were discarded, and the integration failed for 4 samples with carrying capacity. Error statistics, tolerance counts and time fractions are for 1,000 completed runs per α selected at random; far-out outliers (above Q3 + 3·IQR of $\log_{10}$ error) are included in these columns but excluded from Figures 6 and S4. All processes relevant for the whole window: share of all completed samples, and of all samples (sampled space), with every process relevant at every time point; 95% Wilson confidence intervals. Data and scripts: results/alpha-robustness-v2 in the code archive (ref. 54 of the main text).

| | **Samples** | | **Completed runs (1,000 analyzed per α)** | | | | | | **All processes relevant for the whole window** | | **Stopped runs** | |
|---|---|---|---|---|---|---|---|---|---|---|---|---|
| **α** | **Total, n** | **Completed, n** | **Median error, % (IQR)** | **90th percentile of error, %** | **Mean error within tolerance, n of 1,000** | **Maximum error within tolerance, %** | **Far-out outliers, n** | **Mean time fraction with all processes relevant** | **Completed samples, % (95% CI)** | **Sampled space, % (95% CI)** | **Analyzed, n** | **Within tolerance, %** |
| **A. Phage–bacteria model** | | | | | | | | | | | | |
| $10^{-5}$ | 16,000 | 1,689 | 0 (0–0) | <0.001 | 1,000 | 99.3 | 0 | 0.95 | 85.6 (83.9–87.2) | 9.04 (8.60–9.49) | 6,290 | 100 |
| $10^{-4}$ | 16,000 | 1,689 | 0 (0–<0.001) | <0.001 | 999 | 98.8 | 0 | 0.85 | 71.6 (69.4–73.7) | 7.56 (7.16–7.98) | 6,290 | 100 |
| $10^{-3}$ | 16,000 | 1,689 | <0.001 (0–0.0073) | 0.016 | 998 | 97.8 | 0 | 0.63 | 48.3 (45.9–50.7) | 5.10 (4.77–5.45) | 6,290 | 100 |
| $10^{-2}$ | 16,000 | 1,689 | 0.043 (<0.001–0.12) | 0.22 | 997 | 96.6 | 0 | 0.38 | 25.2 (23.2–27.3) | 2.66 (2.42–2.92) | 6,290 | 99.90 |
| 0.1 | 16,000 | 1,689 | 0.68 (0.21–1.3) | 2.2 | 997 | 96.6 | 0 | 0.19 | 9.1 (7.8–10.6) | 0.96 (0.82–1.13) | 6,290 | 99.98 |
| 1 | 16,000 | 1,664 | 8.0 (3.3–15) | 22 | 997 | 95.3 | 0 | 0.09 | 3.9 (3.1–4.9) | 0.41 (0.32–0.52) | 6,315 | 99.95 |
| **B. Phage–bacteria model with bacterial carrying capacity** | | | | | | | | | | | | |

| | Samples | | Completed runs (1,000 analyzed per α) | | | | | | All processes relevant for the whole window | | Stopped runs | |
|---|---|---|---|---|---|---|---|---|---|---|---|---|
| α | Total, n | Completed, n | Median error, % (IQR) | 90th percentile of error, % | Mean error within tolerance, n of 1,000 | Maximum error within tolerance, % | Far-out outliers, n | Mean time fraction with all processes relevant | Completed samples, % (95% CI) | Sampled space, % (95% CI) | Analyzed, n | Within tolerance, % |
| $10^{-5}$ | 8,000 | 1,505 | 0 (0–<0.001) | <0.001 | 996 | 97.1 | 0 | 0.64 | 51.4 (48.9–53.9) | 9.68 (9.05–10.34) | 3,511 | 99.97 |
| $10^{-4}$ | 8,000 | 1,505 | <0.001 (0–<0.001) | 0.0015 | 991 | 96.3 | 0 | 0.51 | 38.0 (35.6–40.5) | 7.15 (6.61–7.74) | 3,511 | 100 |
| $10^{-3}$ | 8,000 | 1,505 | 0.0020 (0–0.0093) | 0.020 | 992 | 95.3 | 0 | 0.36 | 23.7 (21.6–25.9) | 4.46 (4.03–4.94) | 3,511 | 99.94 |
| $10^{-2}$ | 8,000 | 1,505 | 0.044 (0.0031–0.13) | 0.25 | 985 | 93.8 | 0 | 0.23 | 12.8 (11.2–14.5) | 2.40 (2.09–2.76) | 3,511 | 99.94 |
| 0.1 | 8,000 | 1,503 | 0.77 (0.18–1.7) | 2.9 | 988 | 92.0 | 1 | 0.13 | 6.9 (5.7–8.2) | 1.29 (1.06–1.56) | 3,513 | 99.94 |
| 1 | 8,000 | 1,489 | 7.2 (2.5–15) | 25 | 988 | 94.8 | 4 | 0.09 | 2.8 (2.0–3.7) | 0.51 (0.38–0.69) | 3,527 | 99.97 |
