## Supplementary figures and images for "Emerging transient dynamics at finite observation scales"

### Combined_Weights_and_Capacity_cc.pdf

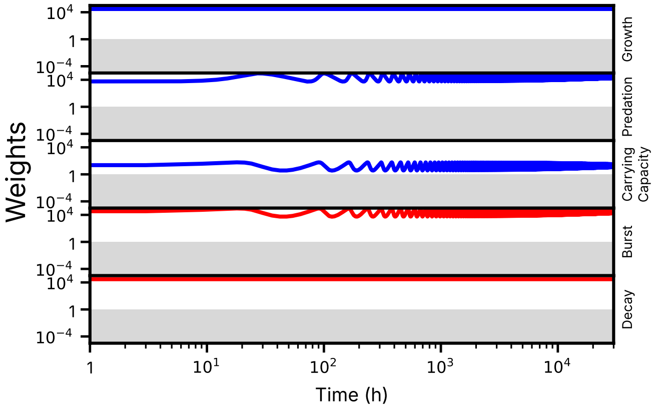

Weights
$10^4$
1
$10^{-4}$
Growth
Predation
Carrying Capacity
Burst
Decay
1
$10^1$
$10^2$
$10^3$
$10^4$
Time (h)

### Combined_Weights_and_Capacity_cc.pdf

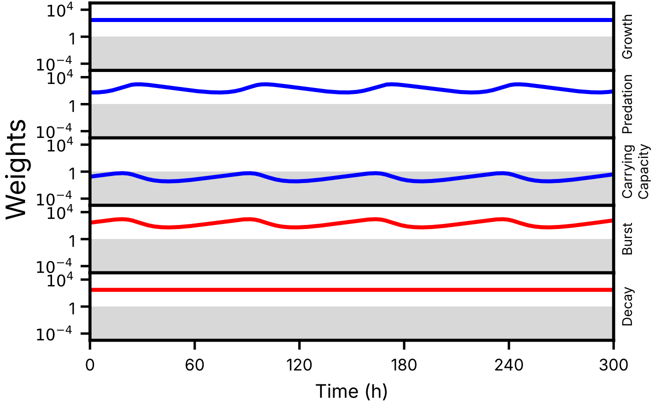
Weights
$10^4$
1
$10^{-4}$
Growth
Predation
Carrying Capacity
Burst
Decay
0
60
120
180
240
300
Time (h)

### Combined_Weights_and_Capacity_cc.png

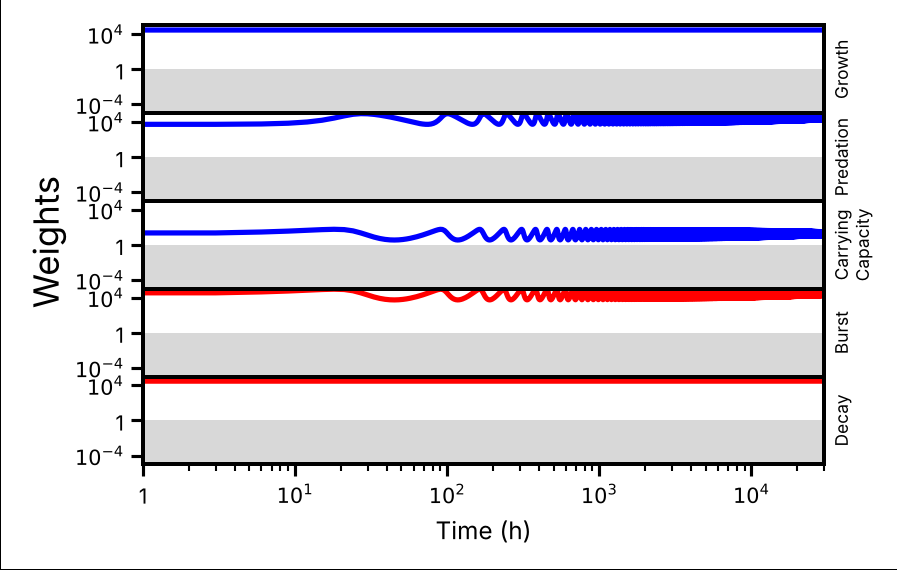

### Combined_Weights_and_Capacity_cc.png

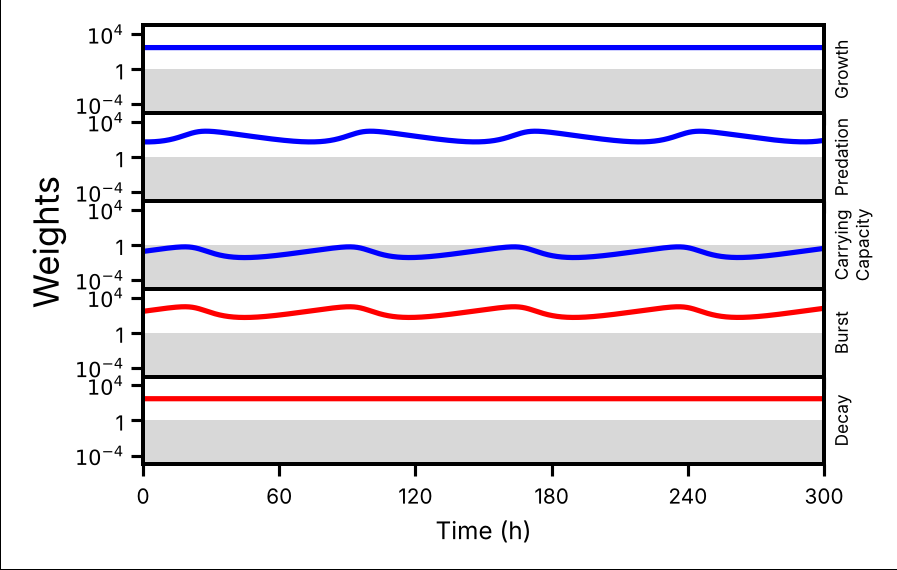

### phase.pdf

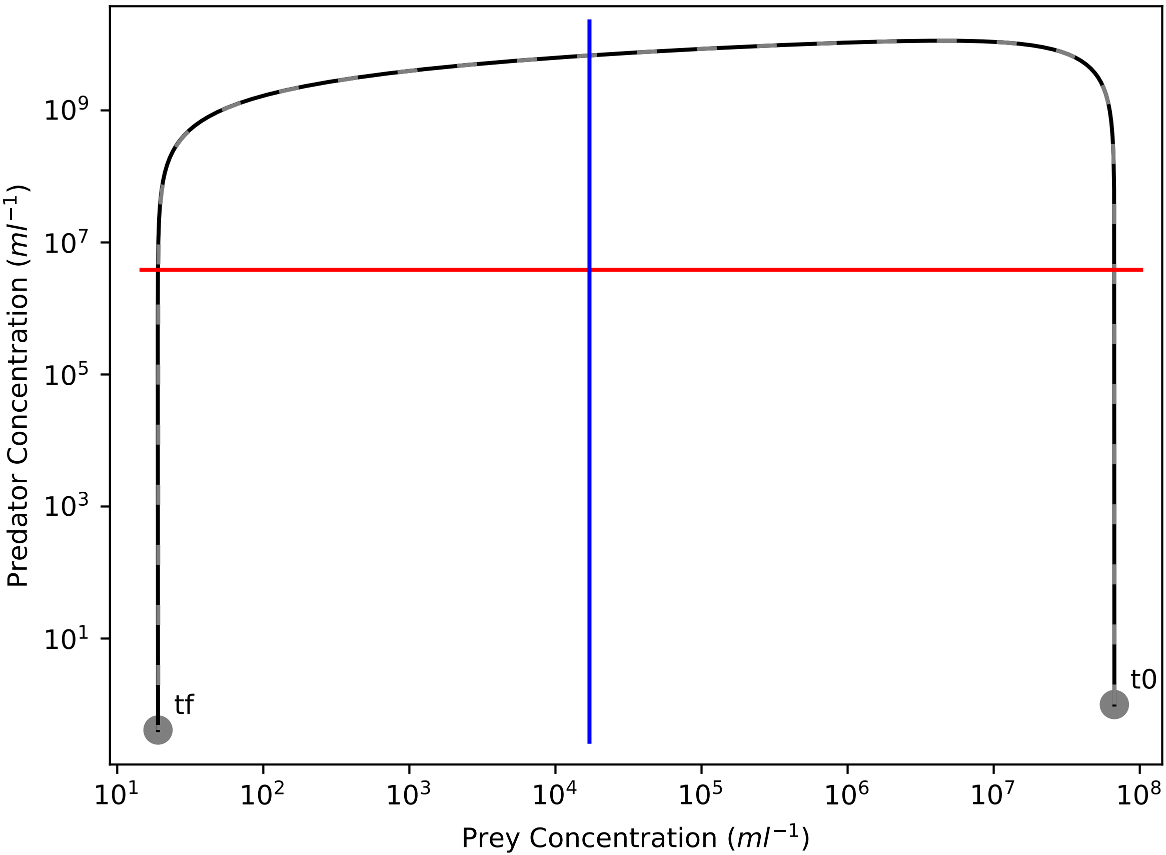

Predator Concentration ($ml^{-1}$)
$10^9$
$10^7$
$10^5$
$10^3$
$10^1$
tf
t0
$10^1$
$10^2$
$10^3$
$10^4$
$10^5$
$10^6$
$10^7$
$10^8$
Prey Concentration ($ml^{-1}$)

### phase.pdf

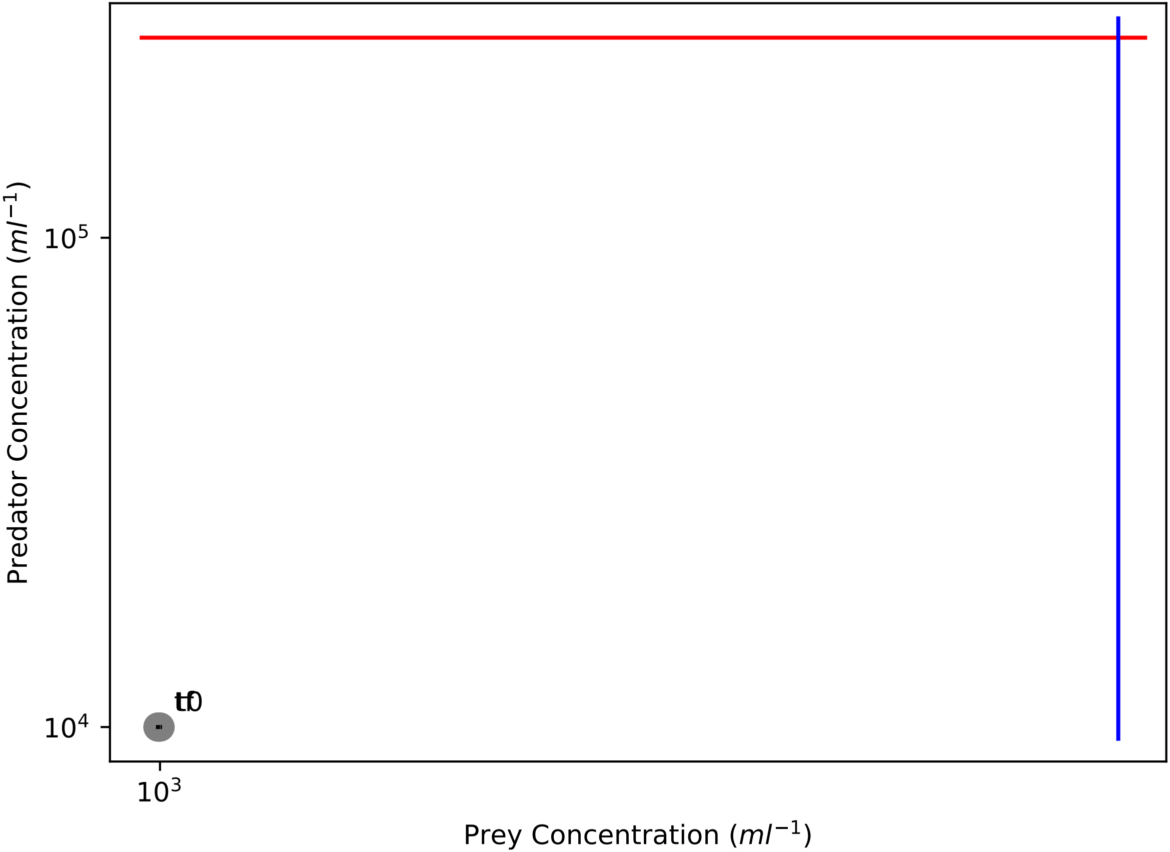

Predator Concentration ($ml^{-1}$)
$10^5$
$10^4$
tf
t0
$10^3$
Prey Concentration ($ml^{-1}$)

### phase.pdf

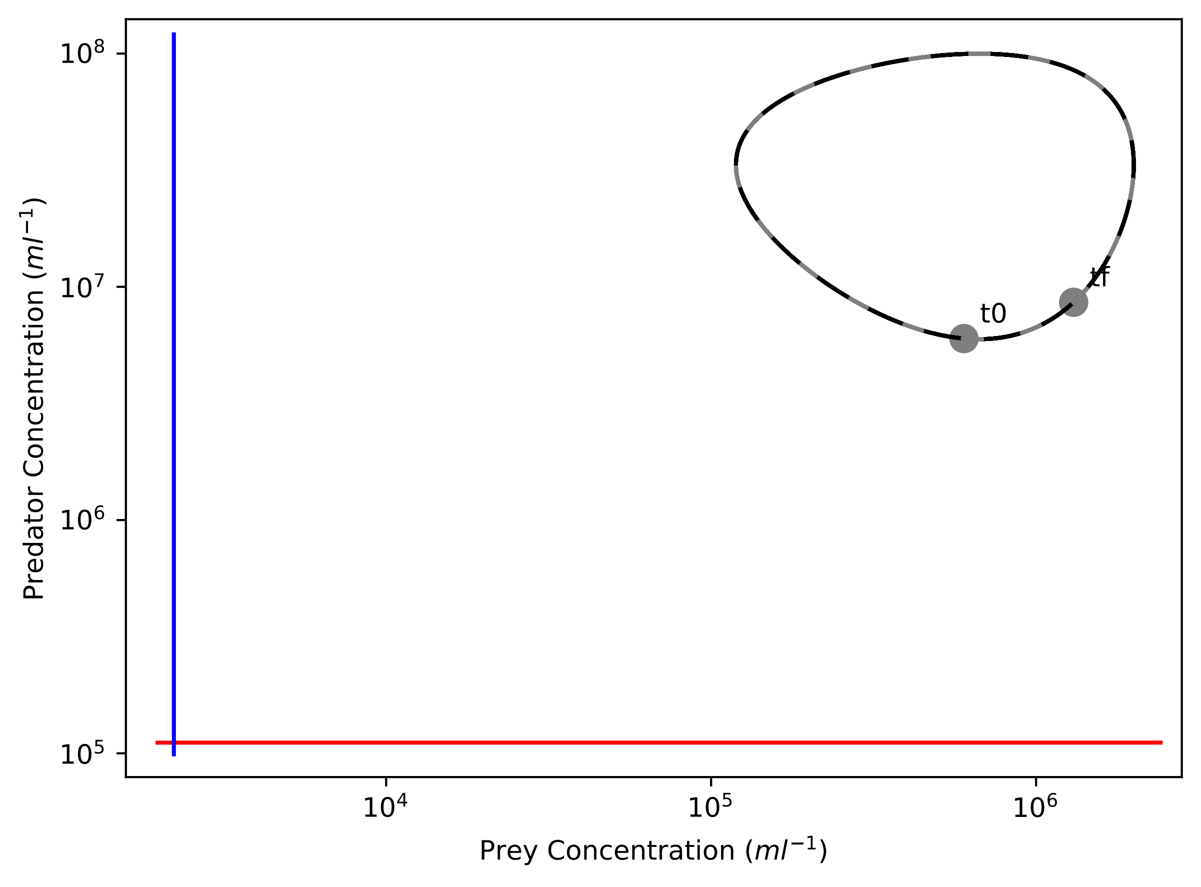

Predator Concentration ($ml^{-1}$)
$10^8$
$10^7$
$10^6$
$10^5$
t0
tf
$10^4$
$10^5$
$10^6$
Prey Concentration ($ml^{-1}$)

### phase.pdf

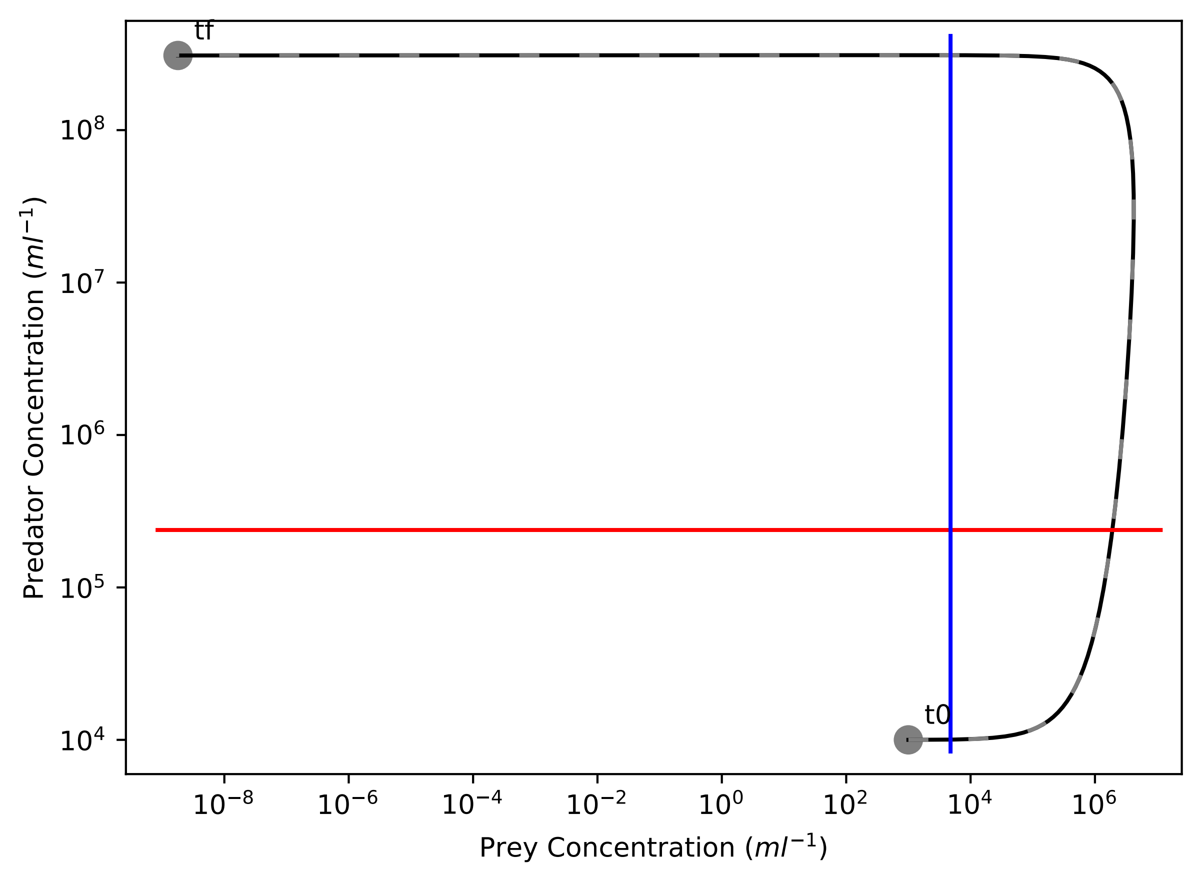

tf
t0
$10^8$
$10^7$
$10^6$
$10^5$
$10^4$
Predator Concentration ($ml^{-1}$)
$10^{-8}$
$10^{-6}$
$10^{-4}$
$10^{-2}$
$10^0$
$10^2$
$10^4$
$10^6$
Prey Concentration ($ml^{-1}$)

### phase.pdf

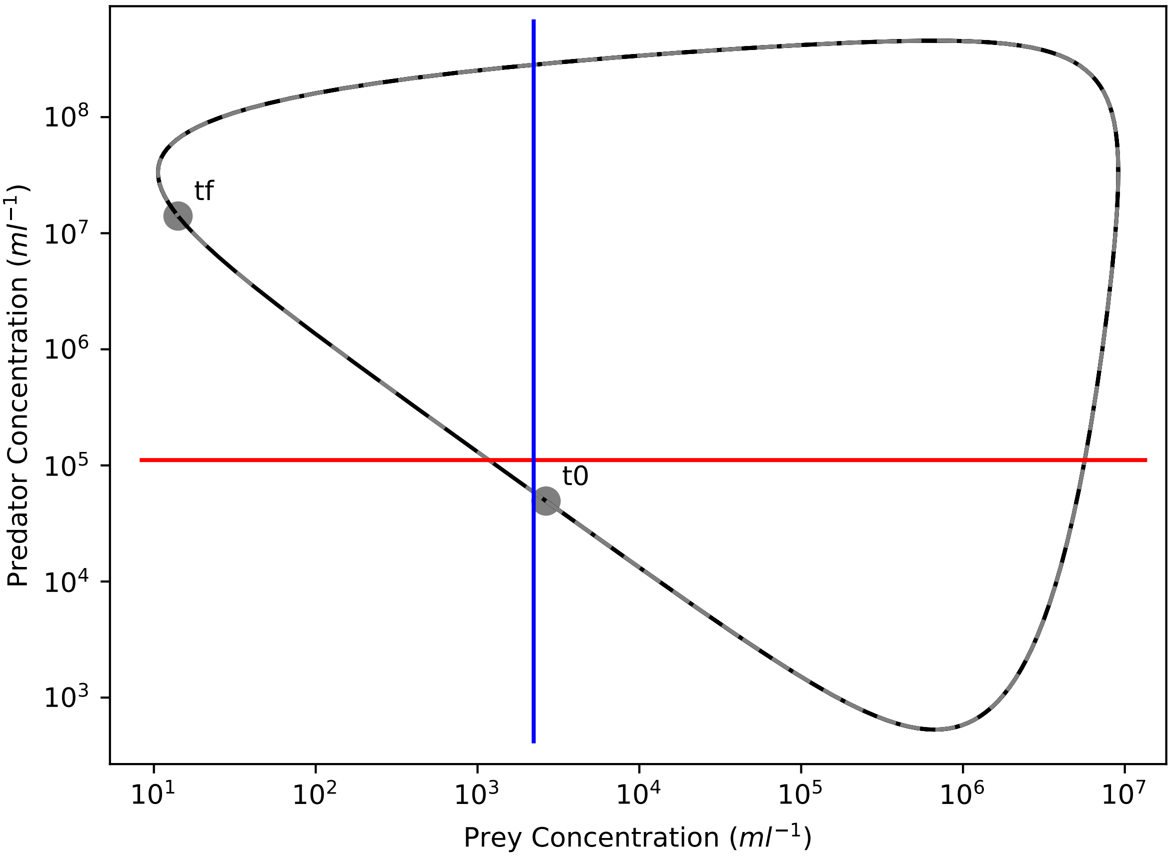

Predator Concentration ($ml^{-1}$)
$10^8$
$10^7$
$10^6$
$10^5$
$10^4$
$10^3$
tf
t0
$10^1$
$10^2$
$10^3$
$10^4$
$10^5$
$10^6$
$10^7$
Prey Concentration ($ml^{-1}$)

### phase.pdf

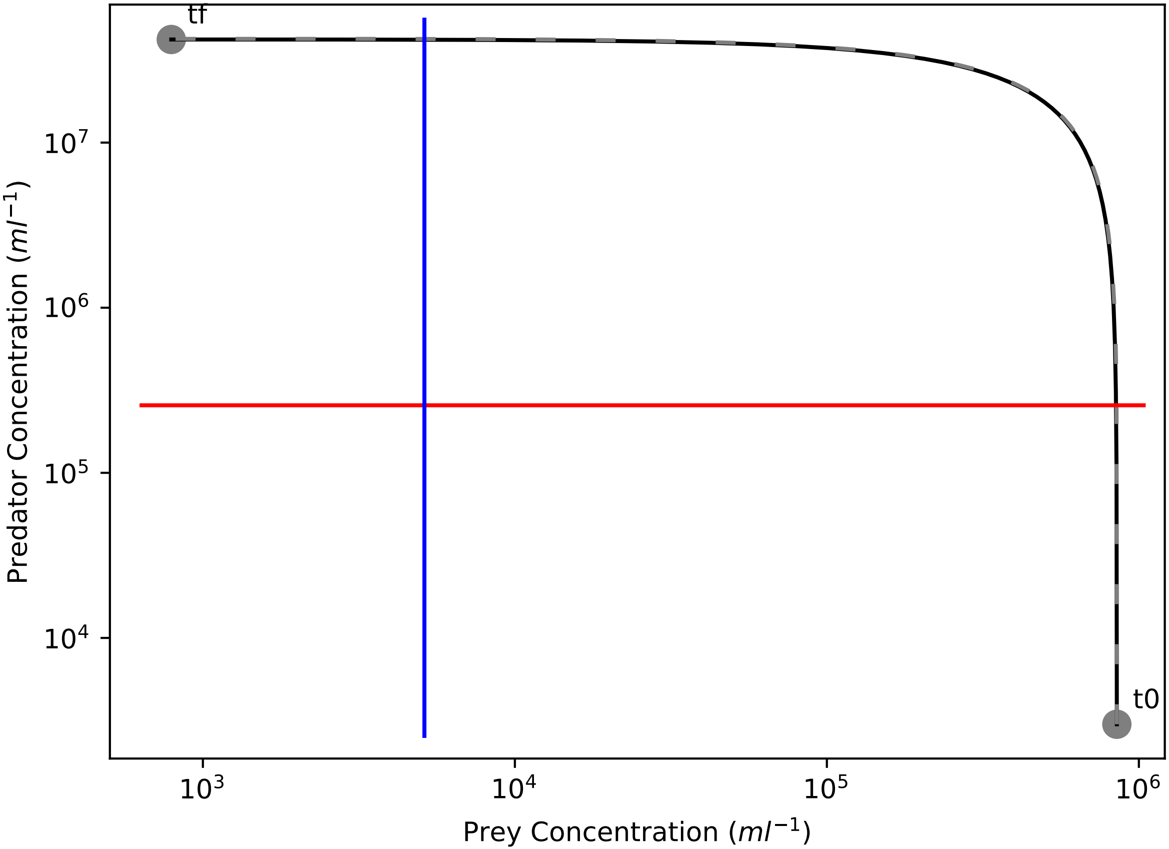

tf
t0
$10^7$
$10^6$
$10^5$
$10^4$
$10^3$
$10^4$
$10^5$
$10^6$
Predator Concentration ($ml^{-1}$)
Prey Concentration ($ml^{-1}$)

### phase_comparison.png

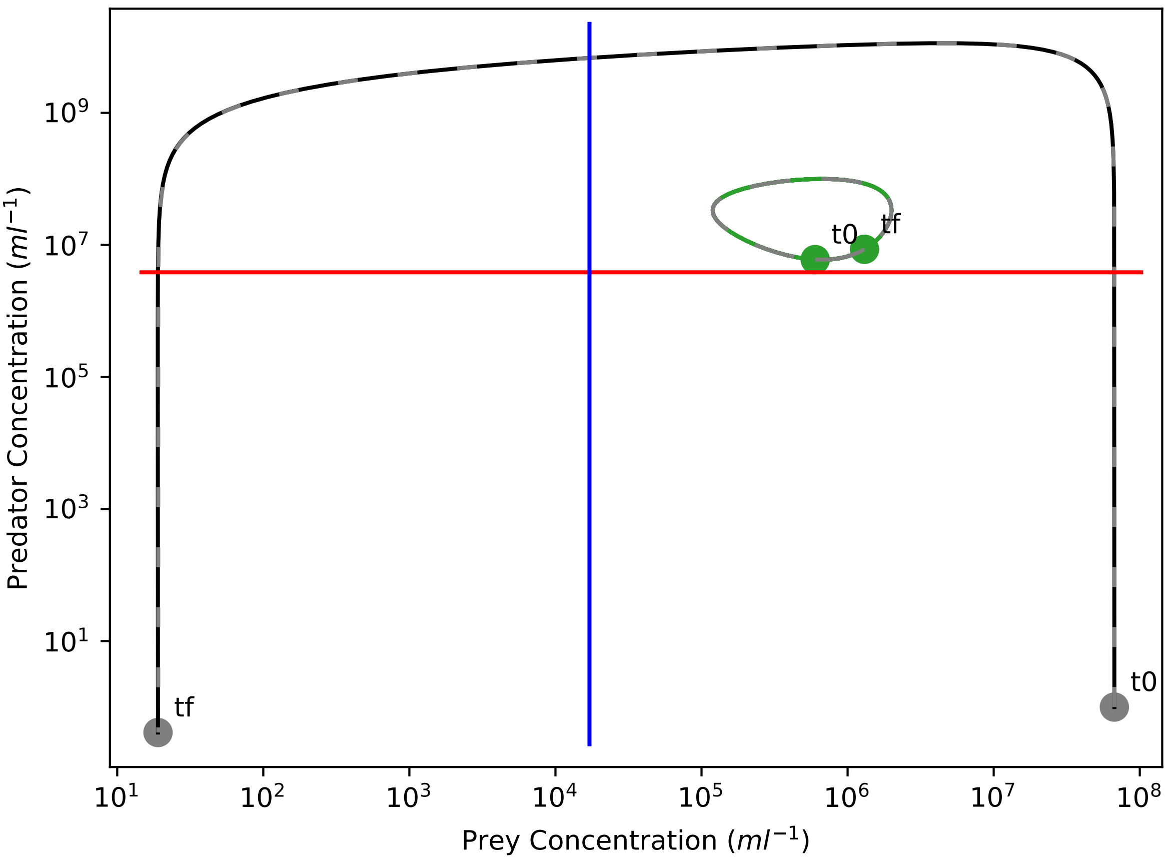

Predator Concentration ($ml^{-1}$)
$10^9$
$10^7$
$10^5$
$10^3$
$10^1$
t0
tf
tf
t0
$10^1$
$10^2$
$10^3$
$10^4$
$10^5$
$10^6$
$10^7$
$10^8$
Prey Concentration ($ml^{-1}$)

### population_comparison_graph.png

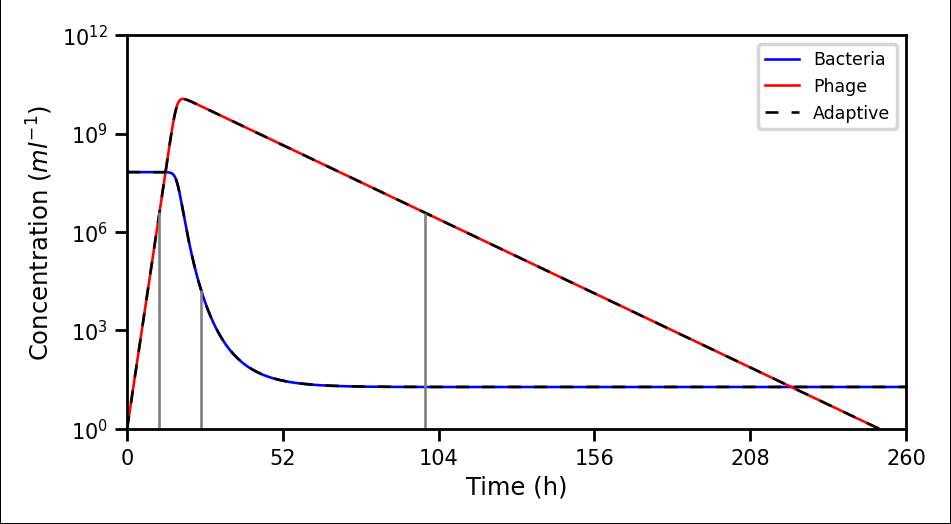

### population_comparison_graph.png

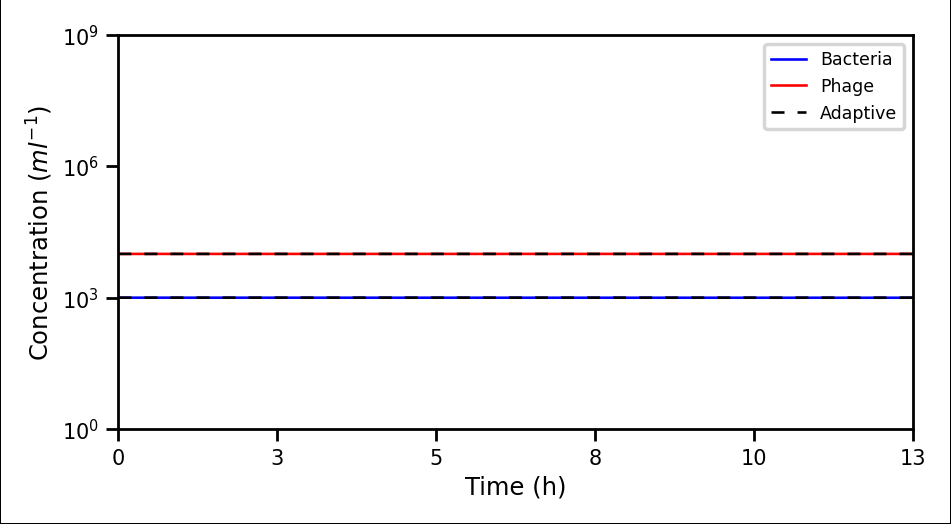

### population_comparison_graph.png

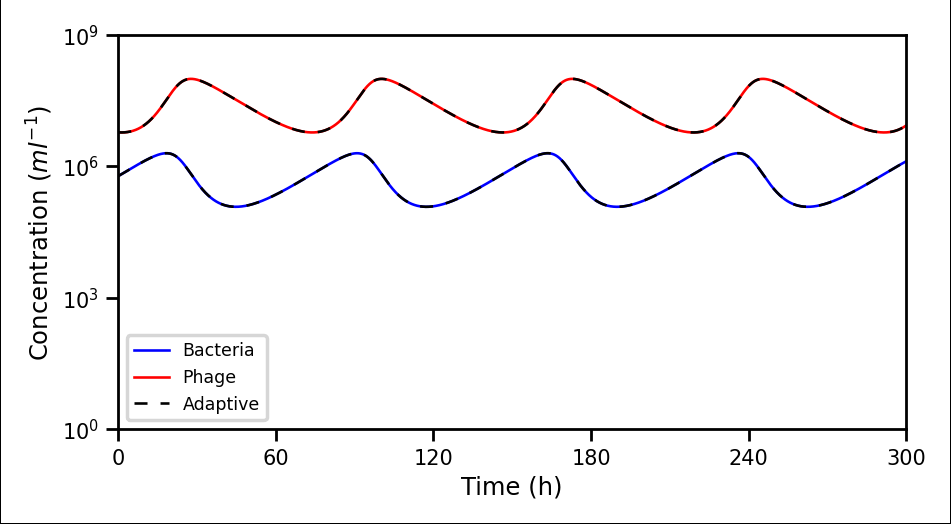

### population_comparison_graph.png

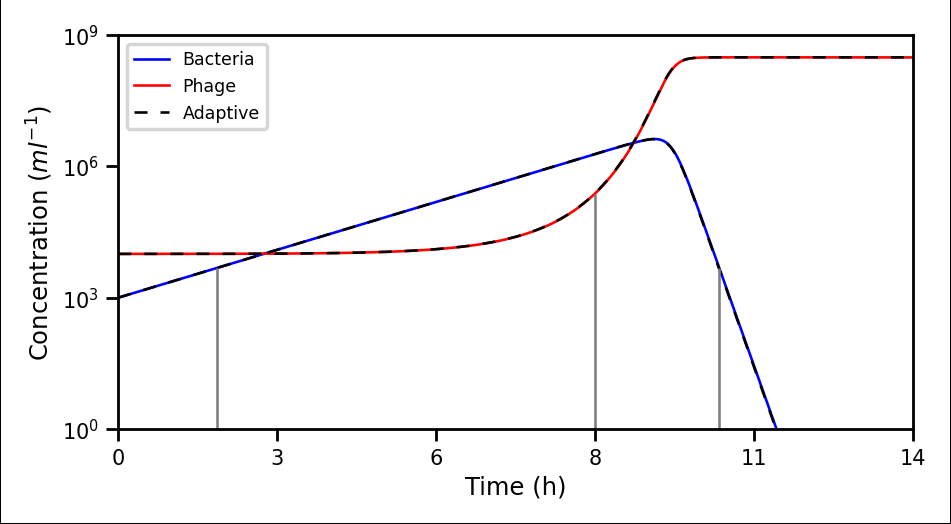

### population_comparison_graph.png

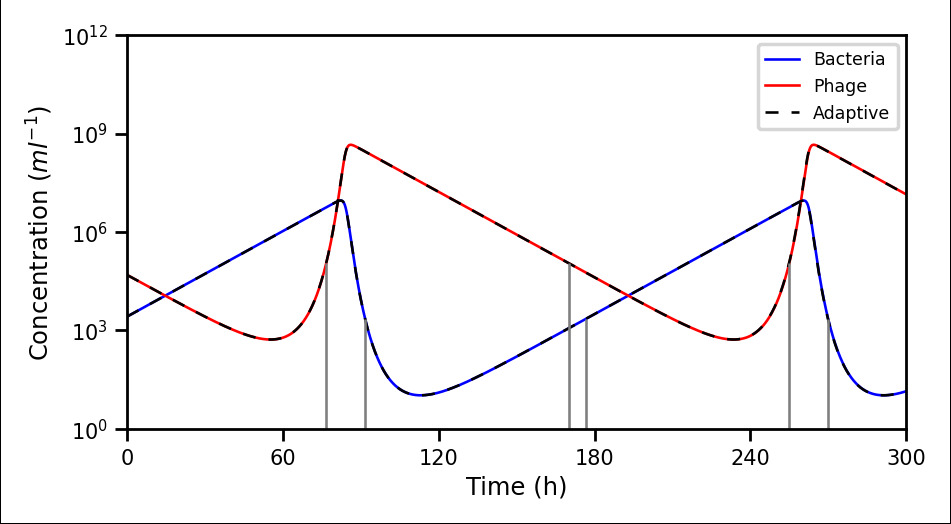

### population_comparison_graph.png

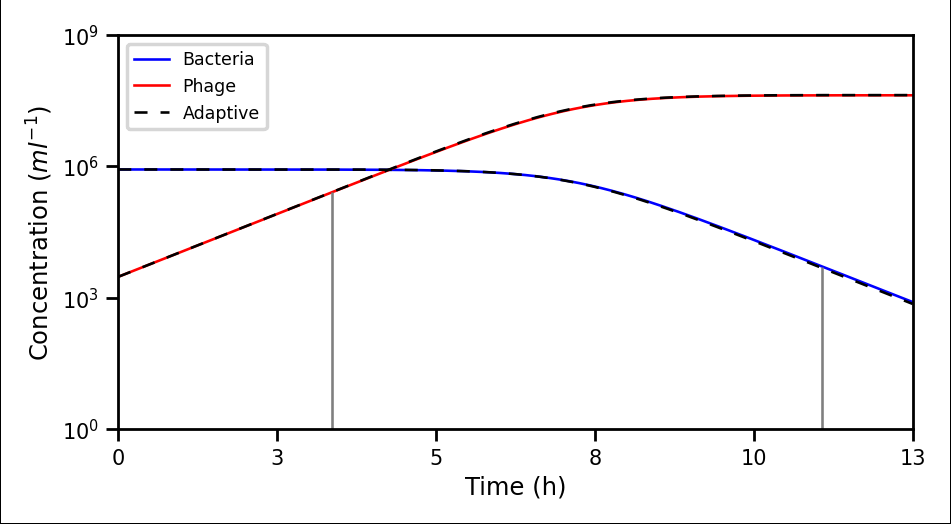

### population_comparison_graph.png

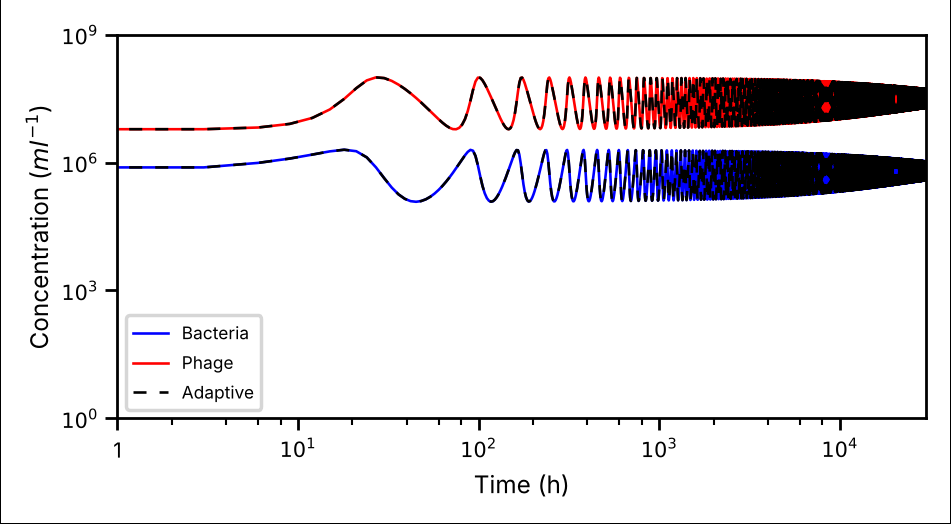

### population_comparison_graph.png

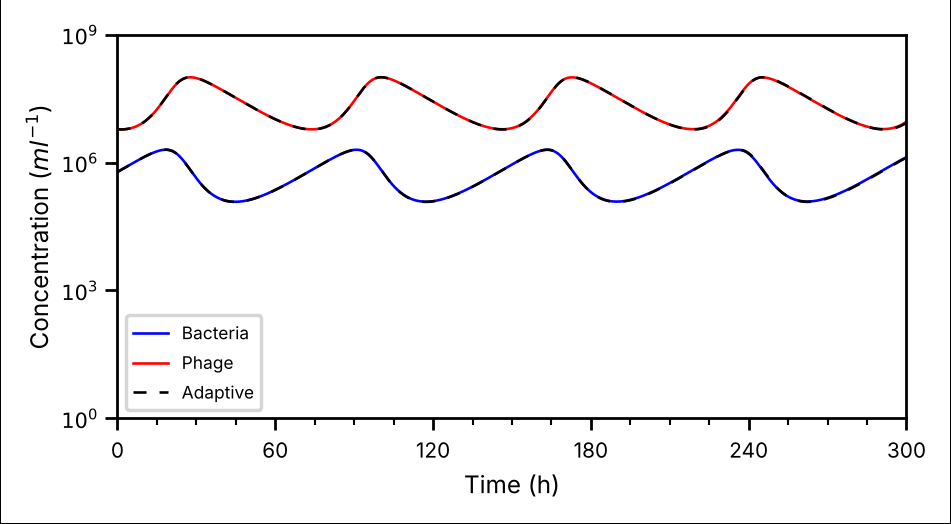

### Primary_Weights.png

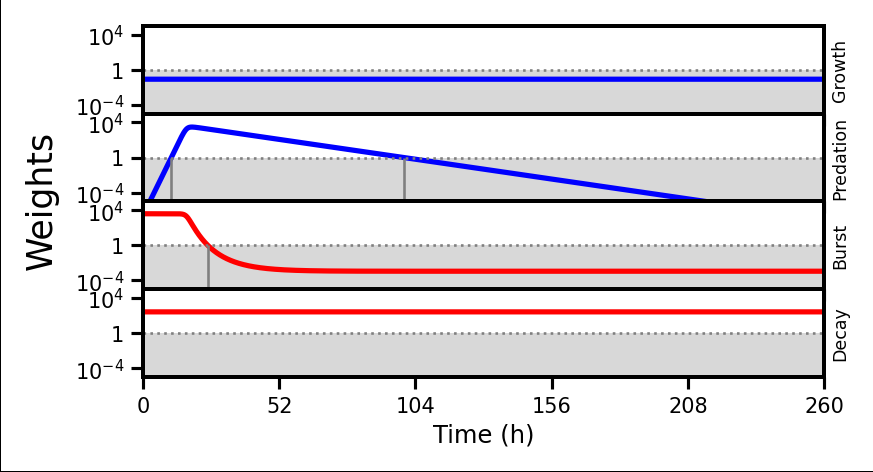

### Primary_Weights.png

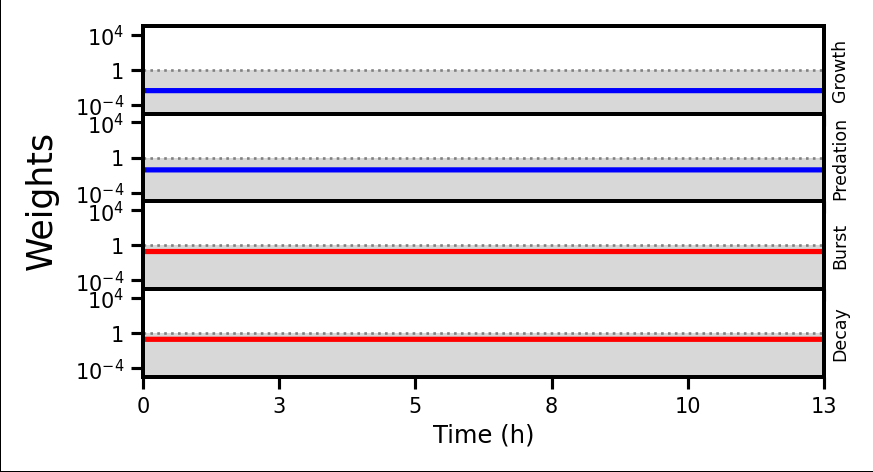

### Primary_Weights.png

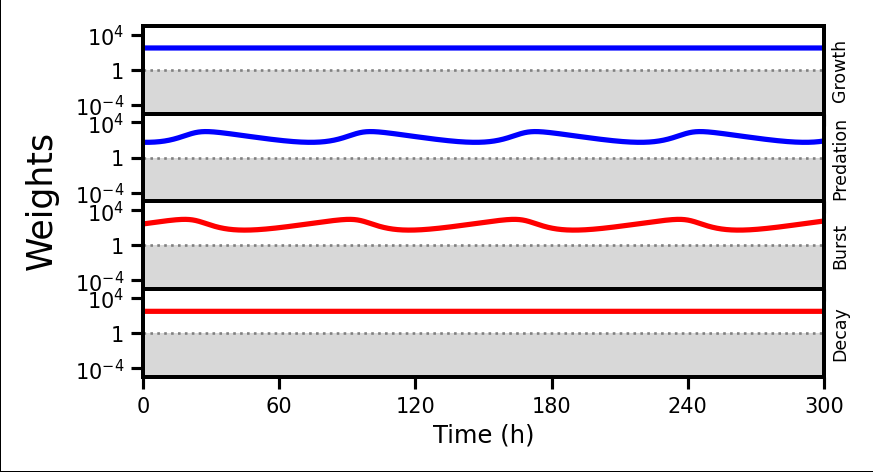

### Primary_Weights.png

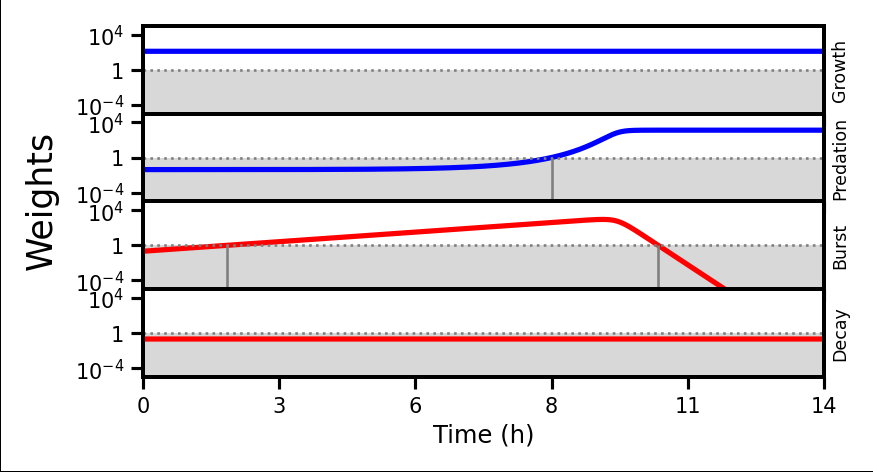

### Primary_Weights.png

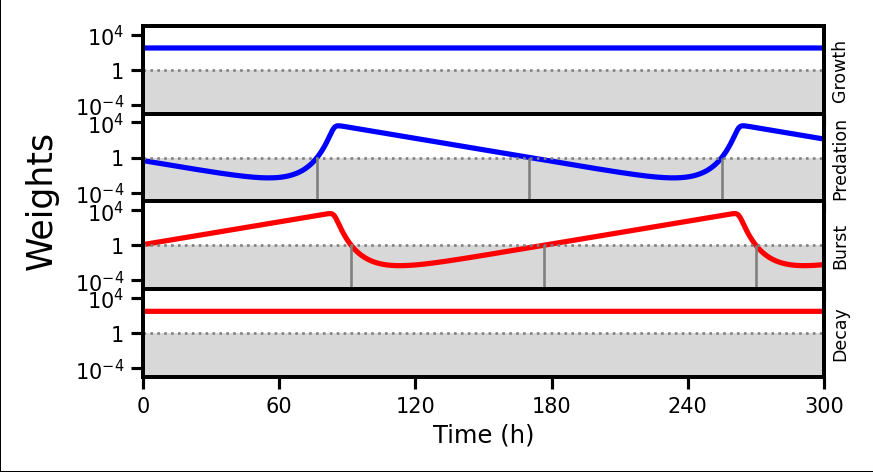

### Primary_Weights.png

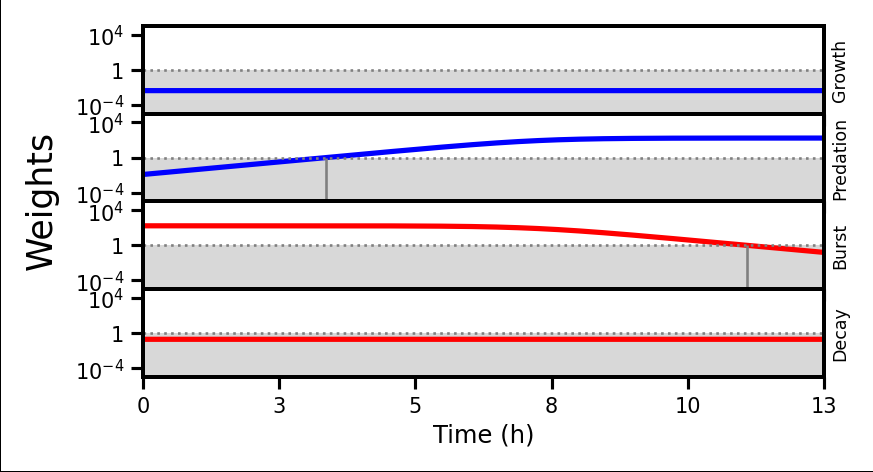

### processes_graph.png

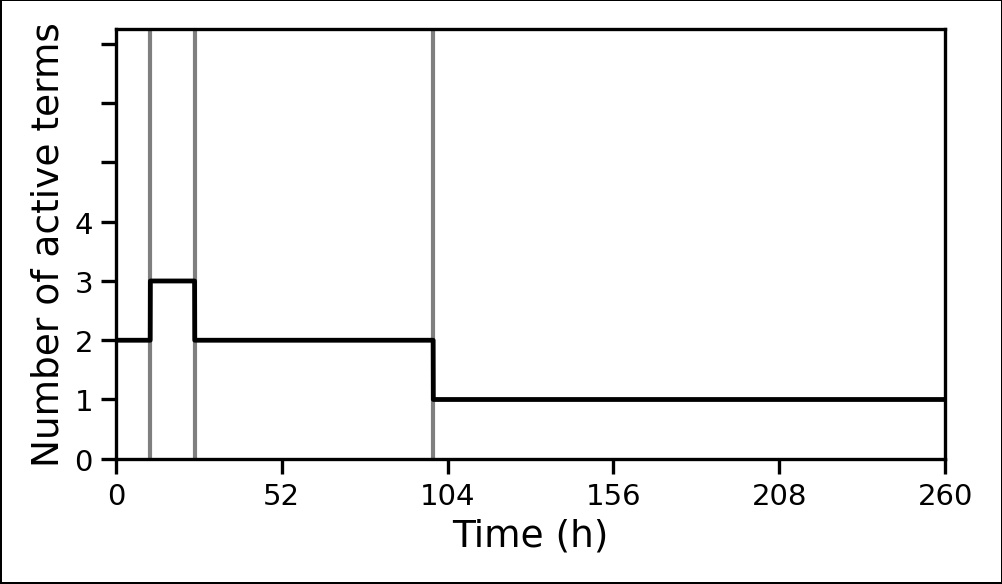

### processes_graph.png

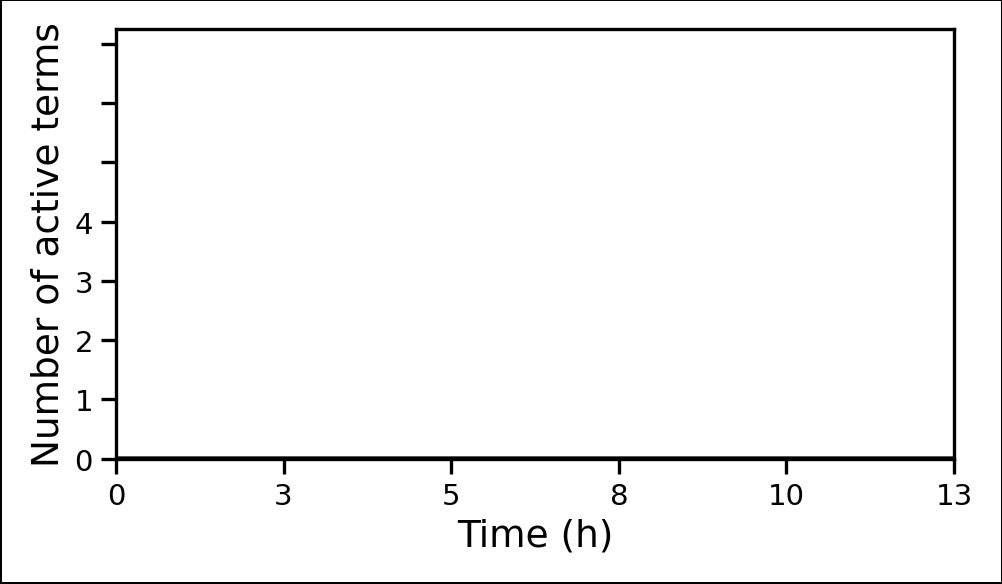

### processes_graph.png

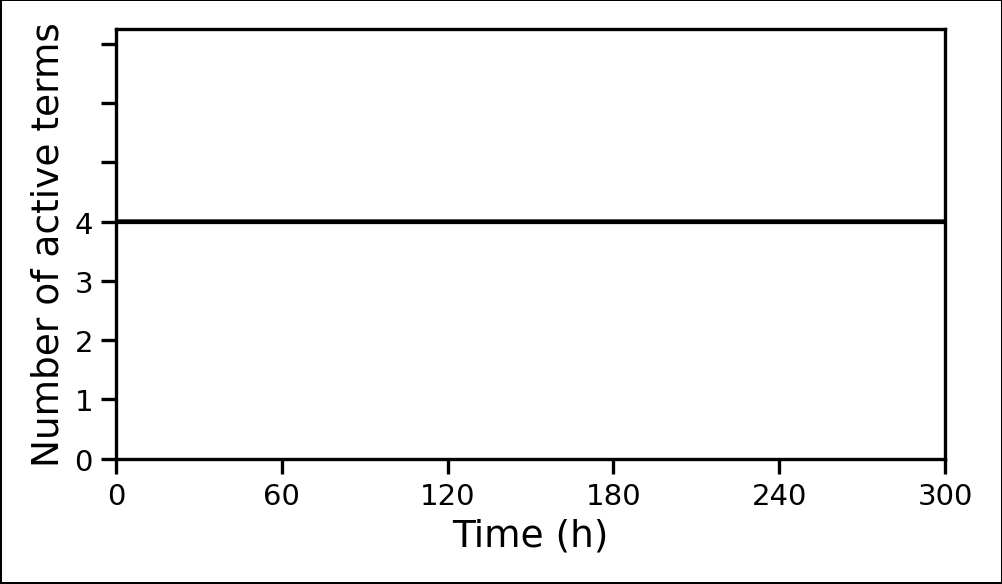

### processes_graph.png

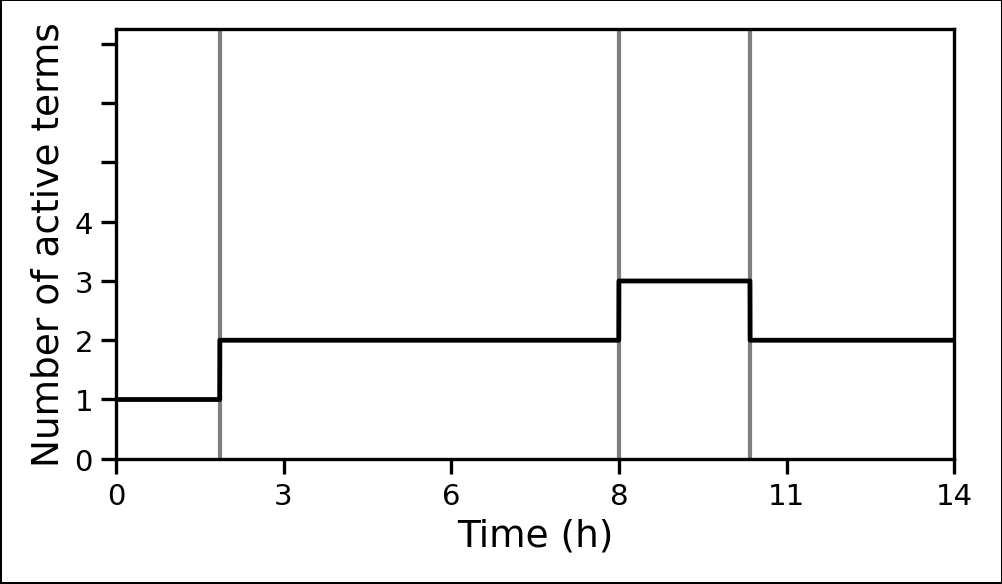

### processes_graph.png

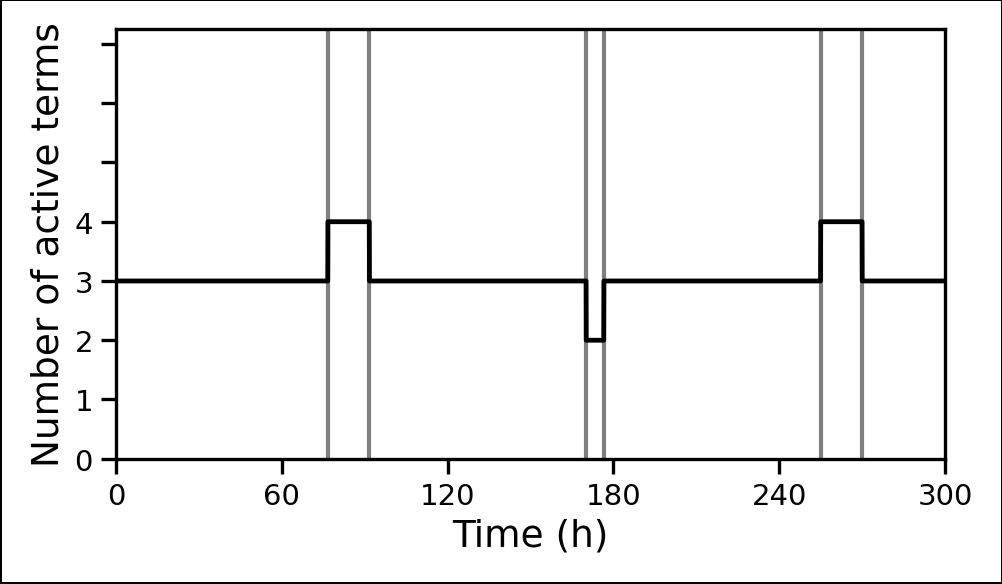

### processes_graph.png

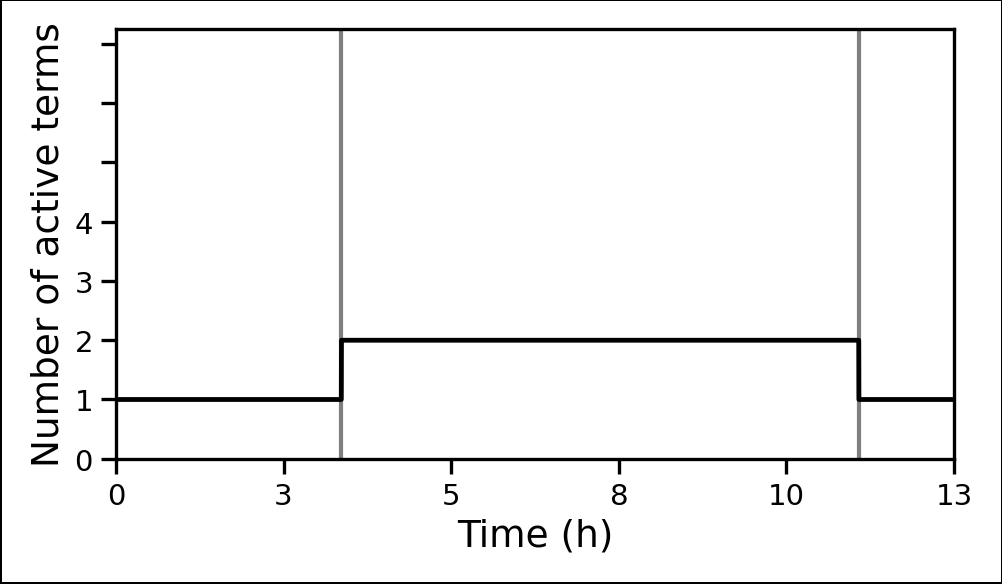

### Regime Category Versus Time.png

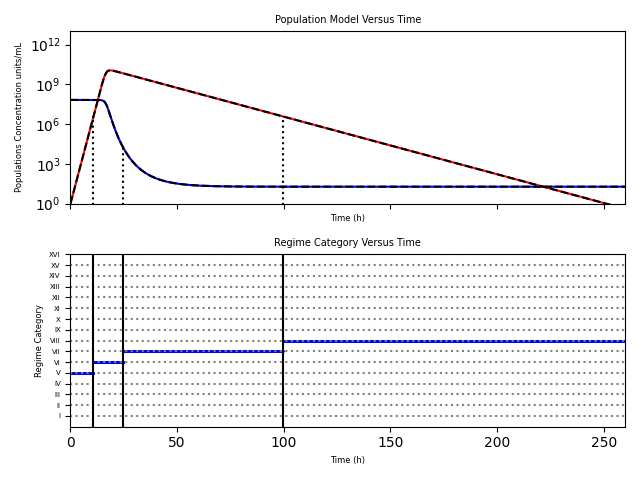

### Regime Category Versus Time.png

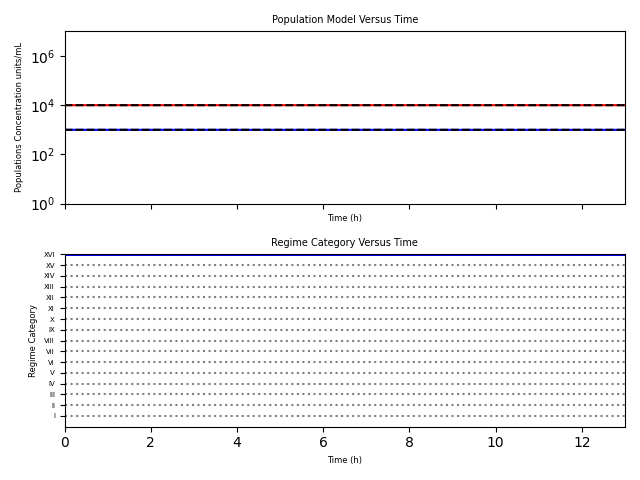

### Regime Category Versus Time.png

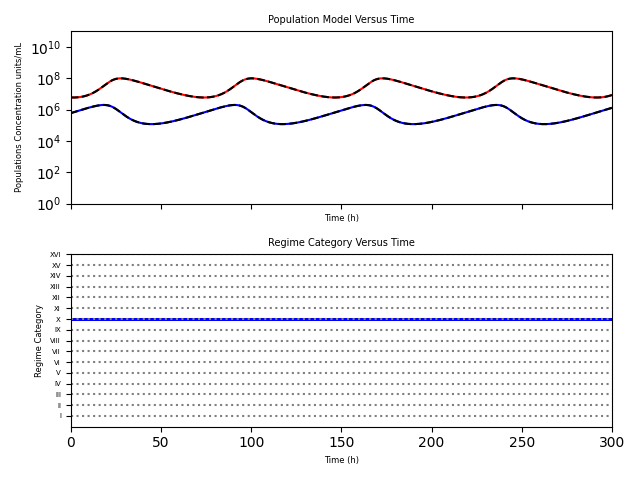

### Regime Category Versus Time.png

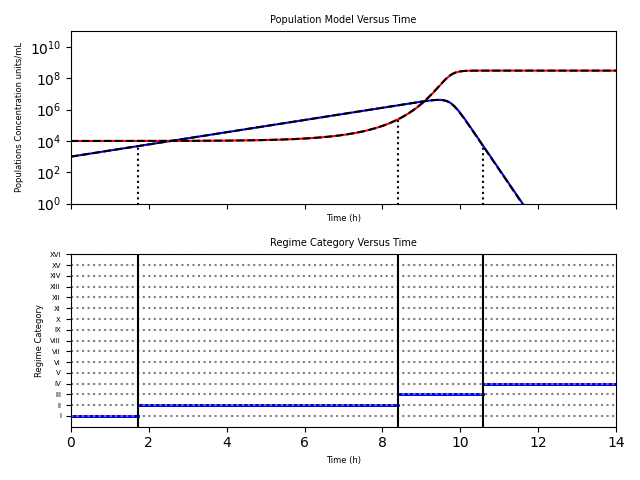

### Regime Category Versus Time.png

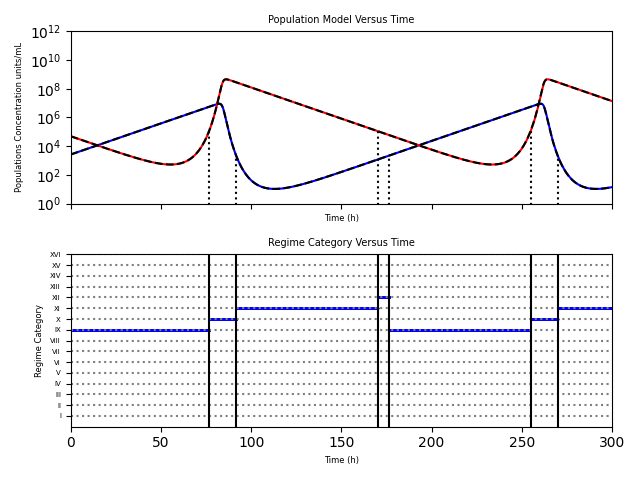

### Regime Category Versus Time.png

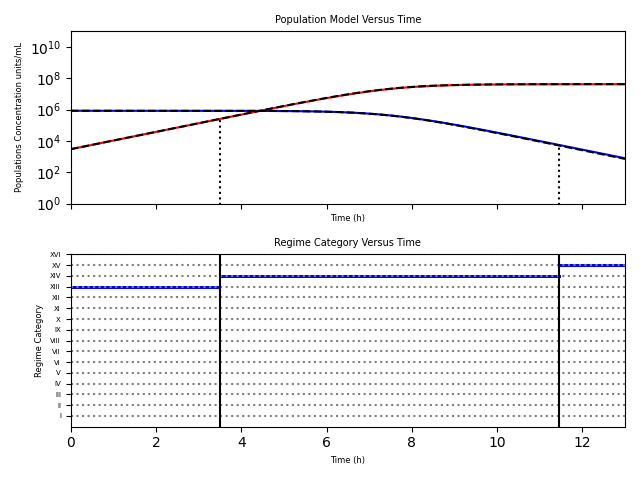

### relative_error_with_inlay.png

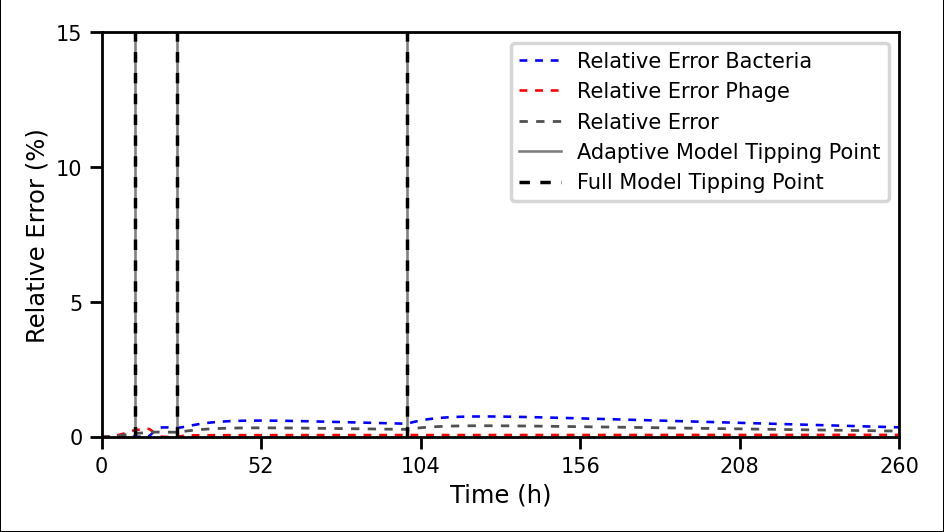

### relative_error_with_inlay.png

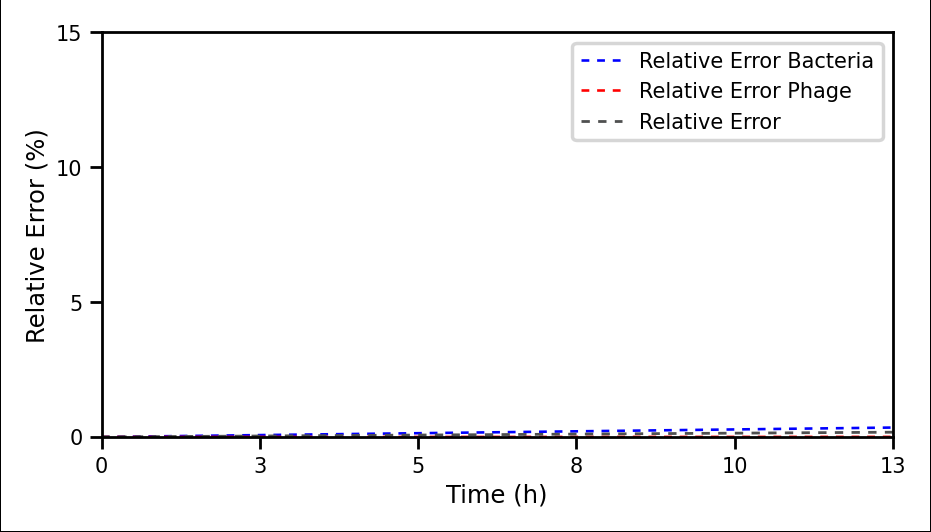

### relative_error_with_inlay.png

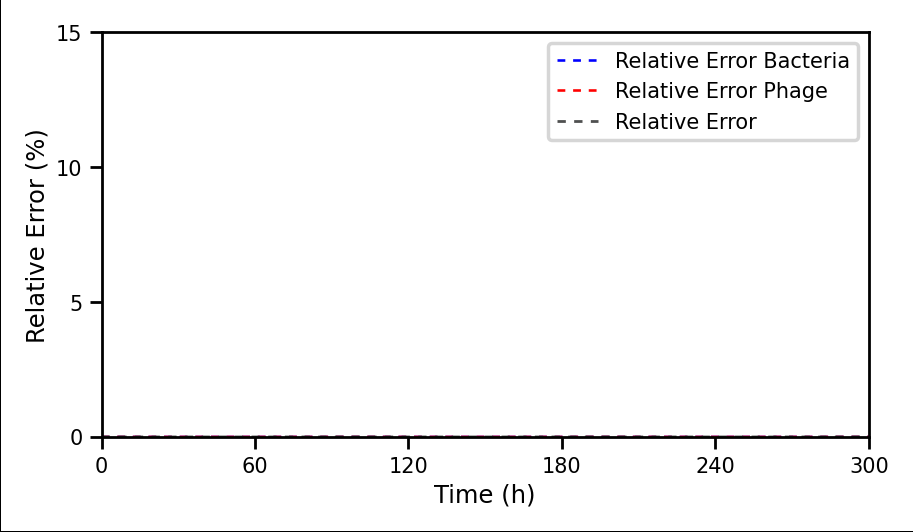

### relative_error_with_inlay.png

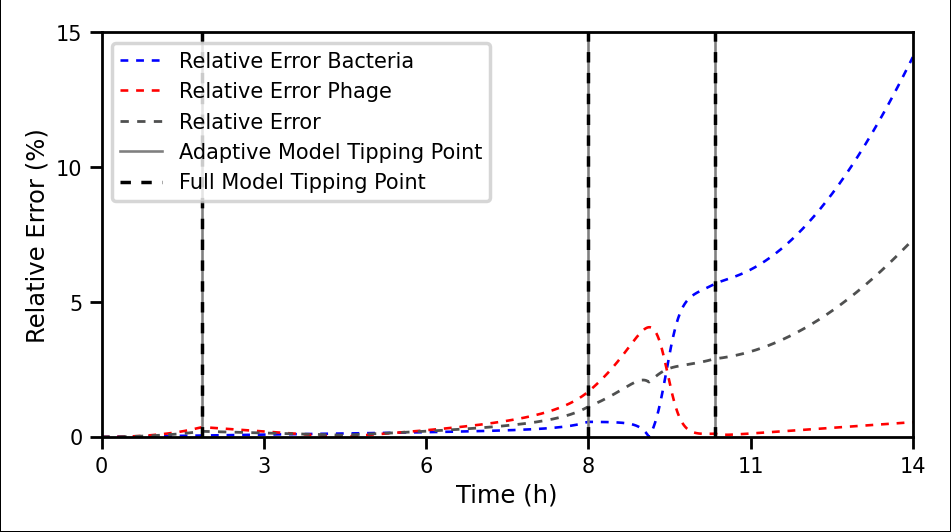

### relative_error_with_inlay.png

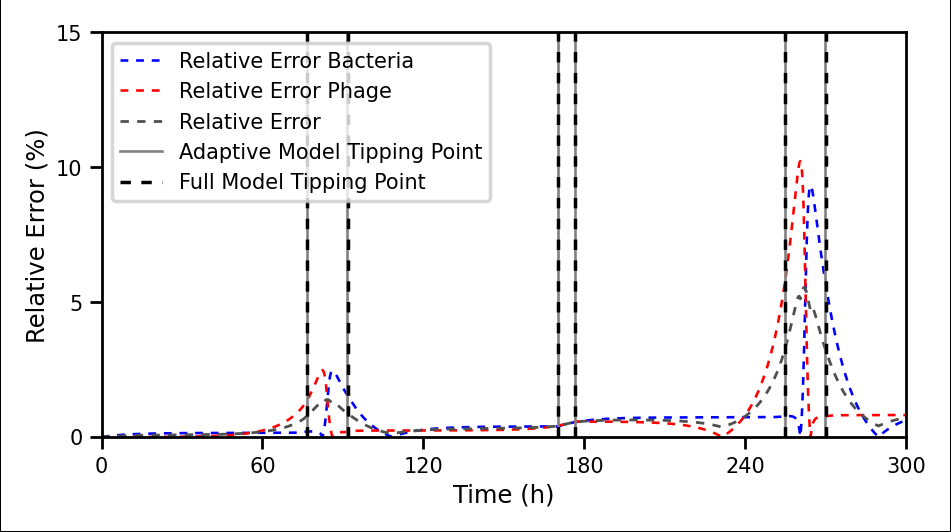

### relative_error_with_inlay.png

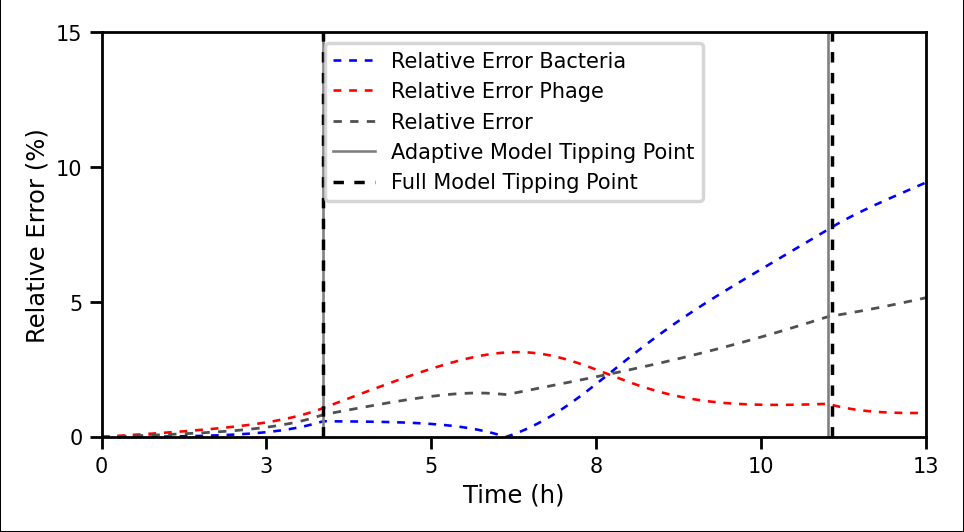

### relative_error_with_inlay_cc.pdf

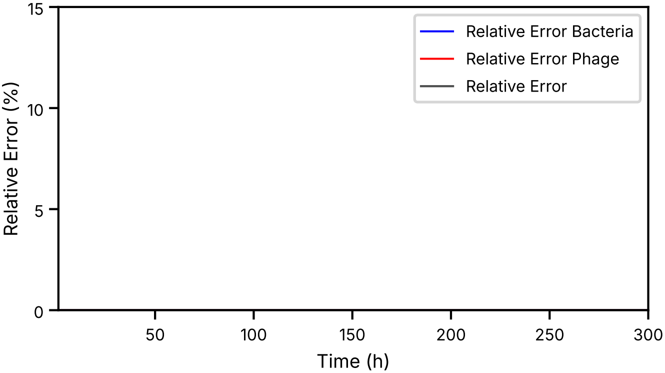

Relative Error Bacteria
Relative Error Phage
Relative Error
Relative Error (%)
0
5
10
15
50
100
150
200
250
300
Time (h)

### relative_error_with_inlay_cc.pdf

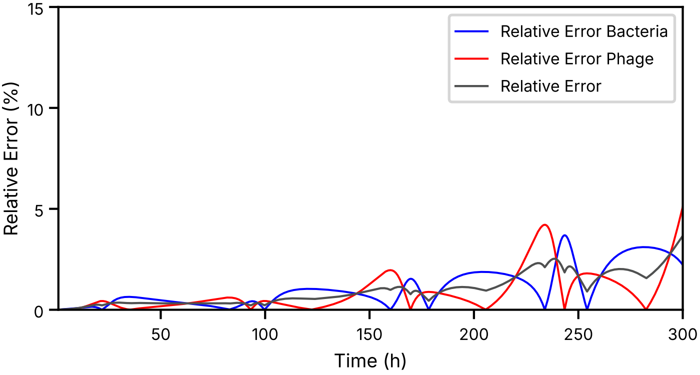

Relative Error Bacteria
Relative Error Phage
Relative Error
Relative Error (%)
Time (h)
0
5
10
15
50
100
150
200
250
300

### relative_error_with_inlay_cc.png

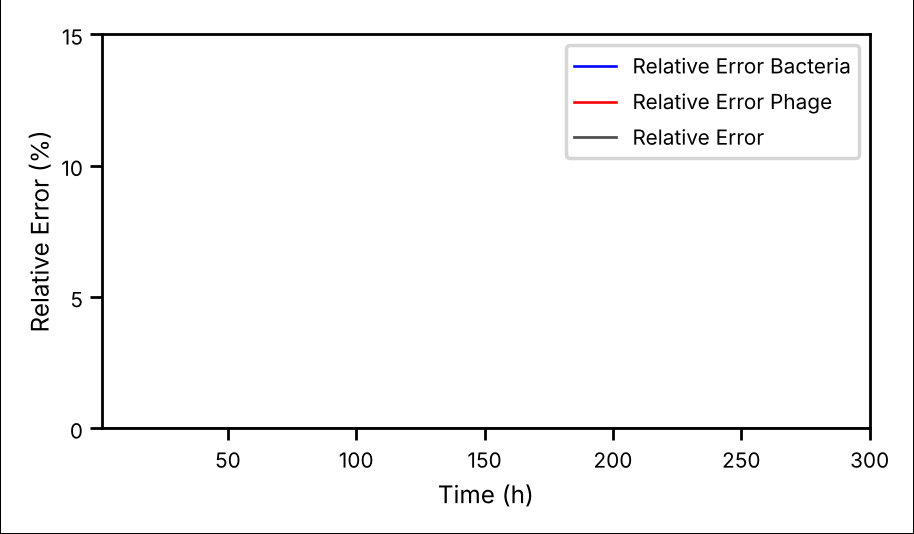

### relative_error_with_inlay_cc.png

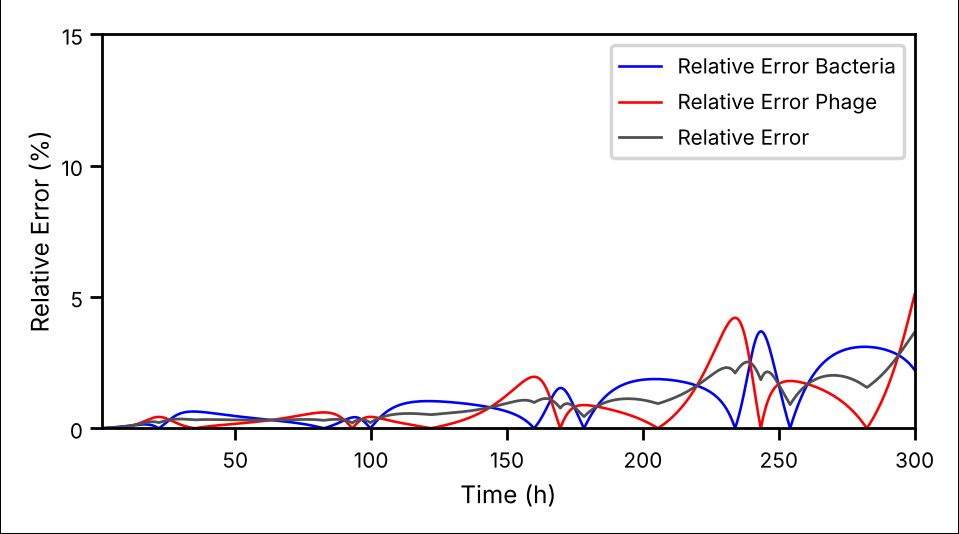
